# A comprehensive computational model of animal biosonar signal processing

**DOI:** 10.1101/2020.12.28.424616

**Authors:** Chen Ming, Stephanie Haro, Andrea Megela Simmons, James A. Simmons

**Affiliations:** Department of Neuroscience and Carney Institute for Brain Science, Brown University Providence, RI 02912; Graduate Program in Speech and Hearing Science and Technology, Harvard University, Boston, MA 02115; Department of Cognitive, Linguistic and Psychological Sciences and Carney Institute for Brain Science, Brown University Providence, RI 02912

## Abstract

Computational models of animal biosonar seek to identify critical aspects of echo processing responsible for the superior, real-time performance of echolocating bats and dolphins in target tracking and clutter rejection. The Spectrogram Correlation and Transformation (SCAT) model replicates aspects of biosonar imaging in both species by processing wideband biosonar sounds and echoes with auditory mechanisms identified from experiments with bats. The model acquires broadband biosonar broadcasts and echoes, represents them as time-frequency spectrograms using parallel bandpass filters, translates the filtered signals into ten parallel amplitude threshold levels, and then operates on the resulting time-of-occurrence values at each frequency to estimate overall echo range delay. It uses the structure of the echo spectrum by depicting it as a series of local frequency nulls arranged regularly along the frequency axis of the spectrograms after dechirping them relative to the broadcast. Computations take place entirely on the timing of threshold-crossing events for each echo relative to threshold-events for the broadcast. Threshold-crossing times take into account amplitude-latency trading, a physiological feature absent from conventional digital signal processing. Amplitude-latency trading transposes the profile of amplitudes across frequencies into a profile of time-registrations across frequencies. Target shape is extracted from the spacing of the object’s individual acoustic reflecting points, or glints, using the mutual interference pattern of peaks and nulls in the echo spectrum. These are merged with the overall range-delay estimate to produce a delay-based reconstruction of the object’s distance as well as its glints. Clutter echoes indiscriminately activate multiple parts in the null-detecting system, which then produces the equivalent glint-delay spacings in images, thus blurring the overall echo-delay estimates by adding spurious glint delays to the image. Blurring acts as an anticorrelation process that rejects clutter intrusion into perceptions.

**Author summary:** Bats and dolphins use their biological sonar as a versatile, high-resolution perceptual system that performs at levels desirable in man-made sonar or radar systems. To capture the superior real-time capabilities of biosonar so they can be imported into the design of new man-made systems, we developed a computer model of the sonar receiver used by echolocating bats and dolphins. Our intention was to discover the processing methods responsible for the animals’ ability to find and identify targets, guide locomotion, and prevent classic types of sonar or radar interference that hamper performance of man-made systems in complex, rapidly-changing surroundings. We have identified several features of the ears, hearing, time-frequency representation, and auditory processing that are critical for organizing echo-processing methods and display manifested in the animals’ perceptions.

## Introduction

Echolocation is a real-time active acoustic sense used by bats and cetaceans for spatial orientation and perceptual imaging [1–3]. Sound production mechanisms, the design of the broadcast signals, the sense of hearing, and the neural mechanisms of the auditory system are harnessed into a single cohesive biosonar system [4–6]. Orchestration of these components is a product of evolution rather than engineering design [7]. Nevertheless, in its similarity to radar and sonar, no other animal system more invites thinking about inspiration from biology to technology [8]. The lessons derived from understanding evolved echolocation can be applied to manmade design. Additionally, the strong underpinnings for radar and sonar in applied math and engineering can reflect back to improve our understanding of the biological system as if it were a product of design [9, 10]. Here, we describe a sufficiently detailed computational model of biosonar that it can contribute to both directions of inspiration [11, 12].

Echolocating animals produce sounds that travel outwards, impinge on objects, and form echoes that convey information back to the animal about those objects [2, 3]. For most bats, sounds are generated in the larynx and vocal tract and broadcast into the air through the open mouth or the nose (a few species use tongue-clicks instead); for dolphins, sounds are generated in a pair of air-filled sacs called phonic lips near the blowhole on the top of the head, and then the sounds are projected into the water through a specialized structure in the forehead [13–16]. For bats, both the broadcasts and their echoes are received by the external ears and conducted through the middle ears to the inner ears, while for dolphins they are received through sound-conducting channels in the lower jawbones and conducted to the inner ears [14,16,17]. After reception has occurred, the neural mechanisms for the sense of hearing serve as the receiver in biosonar [17–21]. The model presented here is intended to emulate the signal processing that guides movements and forms perceptions of objects and their place in the surrounding scene. We are updating an earlier, bioinspired computational model of echolocation in bats to arrive at a more auditorily-plausible processing scheme that can now be applied to both bats and dolphins [12, 22]. The expanded model is focused on generating output that emulates echo delay accuracy and resolution in both echolocating species [23–32]. New research on the bat’s resistance to interference from background, or *clutter*, echoes [33] expands our earlier model to include the question of whether echolocation leads to true perceptual “images” and whether the content of the percepts images reveals an internal arithmetic of their component parts.

The parent model is Spectrogram Correlation and Transformation (SCAT) [22]. It consists of a single channel of processing to mimic monaural processing by bats. For future work, we will develop a binaural version of the model to examine methods for reconstructing target shape, but for the present the monaural version is what we present here. Based on processing in big brown bats, it incorporates features of broadcast signals, echo reception, peripheral auditory coding, and the reorganization of neural response profiles from acoustic features to object features. The SCAT model provides a conceptual and computational framework for peering into the gap between biological and technological descriptions of biosonar performance. It belongs to a family of computational auditory models that use time-frequency concepts [34] to account for aspects of auditory perception [35]. The model is deterministic, with clearly defined computational pathways and no internal stochastic processes. This conforms with experimental findings that the echo delay accuracy of bats approximates that of a matched filter, or crosscorrelation receiver [36, 37], at moderate and high echo signal-to-noise ratios (the Cramér-Rao region) [24] as well as at low signal-to-noise ratios (the side lobe region) [38]. Echo delay resolution by bats also corresponds to a version of the model’s output [25, 26]. Analysis of these results indicates that noise produced internal to the auditory system does not have a deleterious impact on the bat’s performance. Accordingly, for reasons explained further below, we do not insert conventional integrate-and-fire neurons into the model, especially at early stages where sound transduction occurs and internal noise is most likely to affect performance adversely. Critical stochastic parameters such as refractory times [39], spike time distributions [40], lowpass filtering [41], and converging inputs in early neuronal circuits [42] are potential sources of added temporal variability for registering the timing of sounds. However they are manifested in bats, they do not degrade echo delay accuracy beyond that determined strictly by external acoustic signal-to-noise ratios. Moreover, the values for these parameters of neural responses are not actually known in bats. To fill in this gap, we plan to use the SCAT model as a tool for estimating outside bounds for these neuronal parameters prior to attempting experimental measurements. This keeps the model as presented here squarely in the realm of signal processing, not as neuromorphic modeling.

### Principles of wideband echolocation

#### Echo bandwidth and delay

Most echolocating animals emit wideband biosonar sounds containing ultrasonic frequencies that span up to several octaves (40-150 kHz in bottlenose dolphins (*Tursiops truncatus*); 20-100 kHz in big brown bats (*Eptesicus fuscus*) around a center frequency comparable to the bandwidth around the center frequency [2,3,13–16]. While a sizeable minority of bat species emit narrowband, essentially constant-frequency sounds, too, the SCAT model is wideband by its nature and does not take them into consideration. These systems are truly wideband in the sense that their spectrum is situated in the baseband, anchored at the zero origin of the frequency scale [36, 37]. The core purpose of wideband sonar or radar is to determine the time-delay of echoes with great accuracy by exploiting the presence of many frequencies to enhance estimations of delay [36, 37]. In a process called *ensonification* (illuminating with sound), the transmitted sounds travel outward from the animal to impinge on objects in the surrounding scene and return as echoes [13–16]. Both the outgoing broadcasts and the returning echoes are received by the ears and processed by the auditory system to guide locomotion and perceive objects in the surrounding scene [1–6,14,16]. All of the frequencies in echoes return to the ears at the same time to register the range to the target from echo delay (delays of 5.8 ms/m in air, 1.4 ms/m in water) [2,3,14]. Both big brown bats and bottlenose dolphins can detect changes in echo delay as small as 1 μs [28–32]. For big brown bats, the acuity for detecting changes in echo delay depends quantitatively on having as many of these frequencies as possible [43, 44].

#### Echo amplitude

The other acoustic features of echoes relevant to sonar are their amplitudes and their spectra, which behave differently than delay [14,45,46]. Delay is entirely determined by distance, and it is not affected by frequency. In contrast, several aspects of ensonification influence overall echo amplitudes and echo spectra. Both echo delay and echo amplitude are single-valued features. Each depends on distance, which of course correlates with echo delay, while amplitude additionally depends on object size [46–50]. Amplitude is crucial for determining whether echoes are detected at all [51, 52], but detection is affected also by both echo strength and echo delay [47–49]. In contrast to delay and amplitude, echo spectra consist of amplitude across frequencies, and so intrinsically are multi-valued features. Their individual frequency-by-amplitude values depend on several factors, including target distance, direction, size, and shape [2,3,14,45–54]. To perceive individual objects embedded in sonar scenes, the animals have to unravel echo delay, amplitude, and spectrum for the target and separate those from contributions due to its location, as well as screen out the influence of different background objects in the surrounding.

Echo amplitudes are weakened by outward and returning travel due to the sound spreading out from the source, impinging on an object, and then reradiating, or spreading out again going back to the animal. For small targets in air or water, assuming an open scene, echo attenuation is about 12 dB for each doubling of target range (spherical spreading losses of 6 dB going out, 6 dB coming back) [45–47]. Smaller objects return weaker echoes based on the size of the reflective surface the object presents to the incident sound, called its acoustic *crosssection* [36,37,48]. The acoustic size of a target depends on the wavelengths of the incident sounds in relation to the object’s dimensions [36,37,54]. For bats operating at 25-100 kHz, a sphere 2 cm in diameter reflects an echo that is about 20 dB weaker compared to a flat, mirror-like surface at the same location [50]. Bats and dolphins separate cross-sectional size from distance by exploiting their access to the independent feature of delay to determine distance [47–49].

For big brown bats, broadcasts have amplitudes 10 cm in front of the open mouth of 120-130 dB SPL [1,2,48]. Each broadcast is projected forward in a beam rather than uniformly to the sides [55], so the outgoing sound is received weaker by the side-placed ears than the sound actually ensonifying the scene towards the front. Simultaneous with vocalization, middle-ear muscle contractions attenuate the self-heard broadcast by an additional 30-35 dB from its already somewhat lower amplitude at the external ear and the eardrum to about 70-75 dB at the inner ear [47–49]. This brings the otherwise very intense broadcast down into the range of amplitudes for echoes returning from an insect-sized 1-2 cm sphere located about one meter away [50]. At this close range, the bat aims its full band of frequencies directly onto the target to ensonify it so that all of the available bandwidth impinges on the object [56]. This prevents the echoes from undergoing spectral changes from factors other than shape [11,43,57]. Following vocalization, while the broadcast is traveling outward to impinge on objects at different distances, the middle ear muscles relax in a graded fashion so that hearing for echoes becomes progressively more sensitive as time passes. The rate of hearing improvement is 11-12 dB less attenuation caused by middle ear muscles for each doubling of echo delay or target range, out to a delay of about 10-12 ms, which corresponds to a target distance of almost 2 m [47]. Going in the more natural (opposite) direction from long to short range as though during aerial interception, as the bat approaches a small target, the echoes become stronger by 12 dB per halving of range, while the bat’s sensitivity for hearing the echoes becomes less sensitive by 11-12 dB per halving of range. These two countervailing processes stabilize the strength of the echoes at a fixed *sensation level* (decibels above hearing threshold) once the target is nearer than 2 m [47]. Even though echoes increase progressively in strength with further declining target range during the bat’s approach, concurrent reduction in hearing sensitivity occurs as delay shortens, so that the bat cancels out changes in echo amplitude due to distance, leaving changes caused only by fluctuations in the target’s reflectivity from one echo to the next [53]. These fluctuations represent the animation of the target, and they are passed through to the inner ear because the middle-ear gain control only adjusts for echo delay. During pursuit, or when following a moving object, big brown bats aim the head and sonar beam (cast through the open mouth) directly at the target with surprising accuracy [58, 59]. The bat’s flight path is coordinated with the aim of the broadcast beam as it homes in to intercept the target while following a curving, oblique track [60]. Keeping the transmitted sound beam locked onto the target while tracking its direction prevents any wavering of amplitude or spectrum from excursions of the target to the left or right of the beam’s axis. Collectively, these processes compartmentalize information about echo amplitudes and spectra associated with the object’s location separately from its size and shape. They are part of the suite of processes that makes biosonar an active perceptual system [61, 62].

#### Echo spectrum

The previous section describing sonar beam aim and delay related gain control emphasized eliminating unwanted variability of echo features during aerial interceptions. Echo spectra affect the bat’s perceptions, too. The first factor affecting the echo spectrum is sound absorption, due to frequency-dependent propagation path-length [2,14,45,46]. As a rough rule of thumb, for bats operating at 25 to 100 kHz, atmospheric absorption accumulates at rates from about 0.1 to 1 dB per meter, in proportion to frequency. For dolphins operating at 40 to 150 kHz, underwater absorption is about 0.01 to 0.04 dB/m [3,3,14,46]. The largest consequence is to limit the maximum operating range of echolocation because the steady accumulation of small amounts of absorptive attenuation eventually equals and then overwhelms spreading losses, which extinguishes the detectability of echoes [52]. For example, big brown bats can detect insect-sized spheres of 0.48 to 1.9 cm diameter at distances of 3 to 5 m [47], whereas bottlenose dolphins can detect fish-sized spheres of 2.5 to 7.6 cm diameter at distances of 73 to 110 m [3,14,51]. The operating range is over 20 times farther for the dolphin, due to lower sound absorption coefficients in water compared to air. External to the animals, in spatial terms, the size of the relevant sonar scene for small objects is about 100 m for dolphins [51], but it is barely 5 m for bats [47]. Internal to the animals, however, in terms of echo delay, the difference is considerably smaller because the velocity of sound is roughly five times faster in water than in air. For big brown bats, the maximum effective echo delay is about 30 ms for small targets [45]. For dolphins, the equivalent effective delay is under 150 ms. The delay space thus is only 5 times larger for dolphins than for bats. This distinction is important because the scale of the neuronal computations needed to support the acoustics of echolocation depends significantly on the size of the auditory representation of the scene. A 20-fold difference suggests that the scope of the auditory mechanisms for echolocation is likely to be quite different between bats and dolphins, whereas a 5-fold difference may not require so radical a difference in auditory processing. There is another, more specific implication, too: Because atmospheric or underwater sound absorption is lower at the lowest frequencies in the broadcasts, long-range detection depends on picking up the low-frequency end of the broadcast spectrum, around 25 kHz in big brown bats and 40-50 kHz in bottlenose dolphins (Fig 1). Unsurprisingly, the hearing of both bats and dolphins is very sensitive at these frequencies [2,3,49,63]. Below, we address computational implications of the emphasis on low frequencies for registering the arrival of echoes [64].

**Fig 1.**
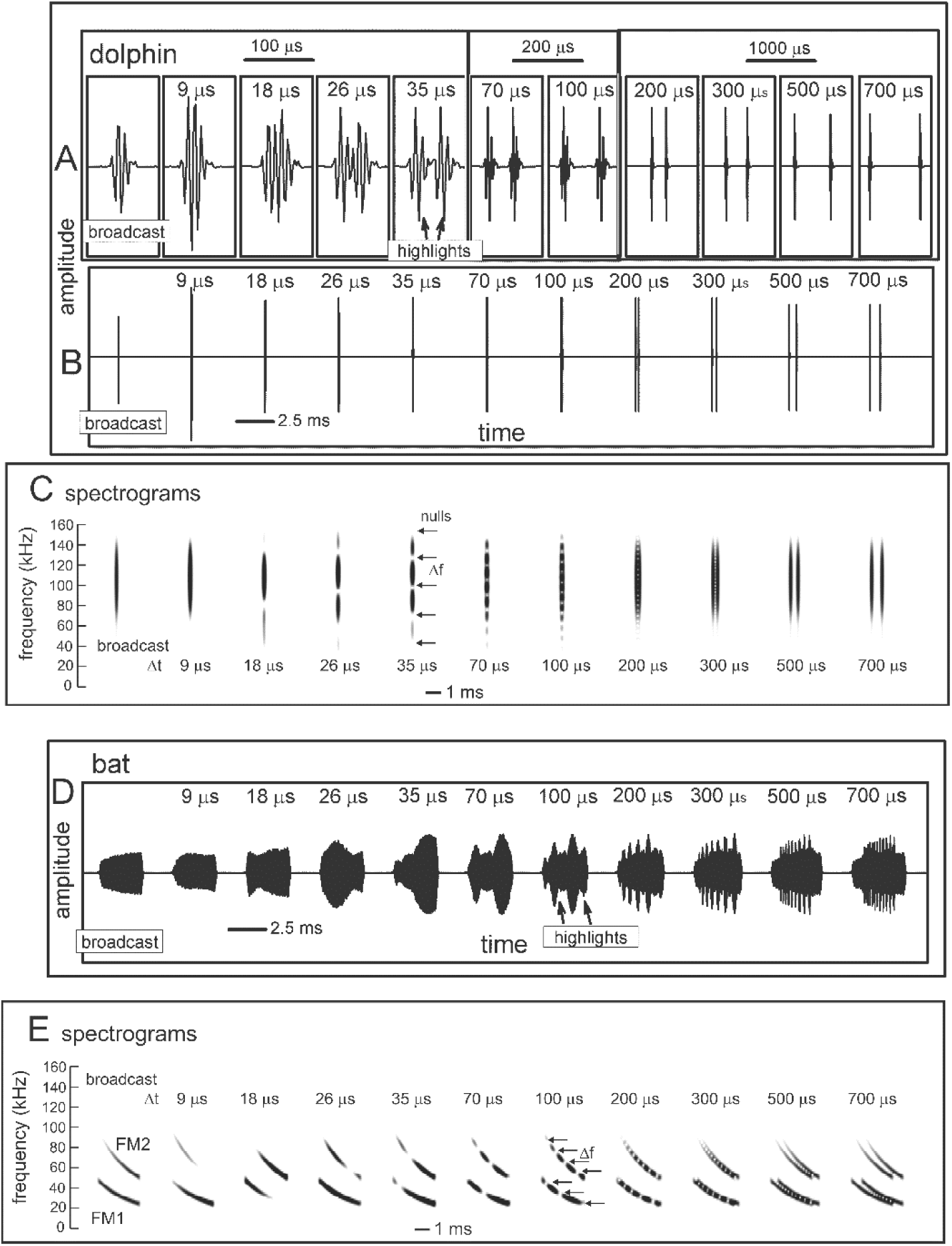
Wideband biosonar broadcasts and echoes. (**A,B**) Time waveforms and (**C**) spectrograms of a bottlenose dolphin echolocation click (broadcast, left) and a series of two-glint echoes with glint-delay spacings (Δt) from 9 to 700 μs (left to right). The click contains only three prominent waves and is about 50 μs long. In the expanded-time waveforms (A), for glint-delay intervals as short as 26 to 35 μs, both reflections are visible as highlights (labeled), and for further spacings they pull apart completely. In contrast, for the spectrograms (C), which have an integration-time of 250 μs, the glint reflections overlap and interfere with each other, creating a single vertical spectrogram ridge that has nulls or ripples in its profile at frequency spacings (Δf) equal to the inverse of the glint time spacings (*e.g*., at 35 μs Δt, the Δf is 29 kHz). Although the two glint highlights are visible in the time waveforms (A) from 26 to 700 μs, in the spectrograms (C) they pull apart only at 300-700 μs. (**D**) Time waveforms and (**E**) spectrograms of a big brown bat FM echolocation chirp (broadcast, left) and a series of two-glint echoes with glint-delay spacings (Δt) from 9 to 700 μs (left to right). The bat chirp is 3 ms long with 1^st^ and 2^nd^ harmonics (FM1 sweeping from 55 to 25 kHz; FM2 sweeping from 90 to 50 kHz). The width of the dark ridges in the FM spectrograms (E) is the integration-time (about 300-350 μs). The duration of the bat chirp is longer than any of the glint-delays, so the time waveforms (D) are completely overlapped from 9 to 700 μs. The highlights in the time waveforms (labeled) are not individual reflections, as is the case for the dolphin click echoes, but instead represent peaks in the interference of the two reflections. The spectrograms (E) have interference nulls at frequency spacings (Δf) equal to the inverse of the glint time spacings (*e.g*., at 100 μs Δt, the Δf is 10 kHz). The spectral nulls appear as well-defined ripples in the dolphin echo spectrograms (C) for time spacings as short as 26-35 μs because all the frequencies appear at the same moment, so the spectrogram ridge is a continuous vertical stripe. For the bat chirps, the dispersion of frequencies along the sweeps and their bifurcation into 2 harmonics (E) obscures the nulls as ripples until the glint-delay spacing is long enough that two or more nulls are present in each harmonic (100 μs). For spacings of 500 and 700 μs, the spectrogram ridges pull apart to form separate echoes.

Echoes arriving from longer distances undergo significant lowpass filtering on top of greater spreading losses, whereas echoes from objects at short ranges more faithfully convey all the frequencies in broadcasts [45,46,49]. Longer-delay echoes can be discriminated from shorter-delay echoes just from the difference in delay because they do not overlap, but the pervasive loss of their higher frequencies due to the lowpass nature of the farther reaches of the scene offers another way to distinguish them. Lowpass echoes have narrower bandwidths, and to the extent that the content of the bat’s perceptions is enhanced by having more bandwidth [43, 44], the resulting perceptions will be less acute, or more blurry, for distant objects [33,65,66], which has implications for auditory computations [11, 12] that are incorporated into the SCAT model described here. This is more important than it seems because both bats and dolphins often search for small targets that are relatively near while in the background are larger, often spatially-extended sources of echoes at longer ranges. While the foreground contains objects of interest that potentially return all the frequencies in the broadcast, the background constitutes clutter that can mask the presence of a target unless it is discriminated against. Exploiting lowpass filtering on echo spectra is an effective means to mitigate clutter interference [33].

A second factor involving the echo spectrum has to be considered here: It is true that long-range clutter can be suppressed simply because the echoes arrive much later, while a target returns echoes at shorter delays. But that depends on the bat or the dolphin emitting each biosonar sound and waiting until all of its echoes return before emitting the next sound. If circumstances require more rapid emissions, such as coping with nearby collision hazards or rapid target movements [64–69], then suppressing long-range clutter requires more complicated mitigation strategies involving the spectrum of echoes [33] which is incorporated into the SCAT model [12].

The third factor affecting the echo spectrum is broadcast beaming. Both bats and dolphins actively track the movements of a nearby target by centering the object on the beam [2,3,14,58,59], thus ensuring that ensonification involves the full broadcast spectrum [56]. As a result, changes in the echo spectrum during tracking reflect properties of the target itself. Off the beam’s axis, the strength of the incident sound is weaker, with higher frequencies falling off to a greater degree than lower frequencies because the beam is not as wide at higher frequencies [2,3,13,14–16,55]. Thus, the spectrum of echoes arriving from objects located off to the sides lack the higher frequencies because these are less strongly projected away from the beam’s axis. By definition these objects are clutter; the animal already has selected one object of interest for its attention, at least momentarily, and its echoes have to be isolated for perception without interference from clutter [33]. Off-axis clutter is more serious than distant clutter because many echoes could arrive from off to the sides at about the same time as echoes from the target, and the lowpass character of clutter echoes is used to help mitigate the inevitable masking effect [57].

The fourth factor affecting the echo spectrum is the target’s structure [1–3,11,14,50,53; see also 36]. Along with cross-sectional acoustic size described above, targets are characterized by the geometric arrangement of their individual reflecting points, called *glints* [70]. In air, the acoustic impedance mismatch that occurs as sound impinges on any solid material is so large that straightforward reflection predominates [2,14,50]. Negligible energy penetrates into the material, and the target does not react to ultrasonic sounds by resonating. In acoustic terms, then, objects are portrayed by the geometry of the reflecting glints. In the earliest behavioral tests, bats distinguish between airborne targets that differ in their glint structure [71, 72]. Using different physical targets designed to have glints at different distances [2,3,73–76] or virtual targets with electronically-controlled glint delays [77–82; see 83, 84], bats perceive shape as a graded feature measured out by the spacing of the glints. For echolocation in air, considering insects as typical targets, the principal body parts (head, wings, and abdomen) scatter the incident sound very strongly [53]. Big brown bats perceive these glint reflections as having distinct arrival-times that represent the separation of the glints from each other along the distance axis [77]. Moreover, they perceive glint spacings of insect-sized virtual targets as a class: A median glint-delay spacing of 100 μs (1.7 cm target glint spacing) is classified as similar to a range of other spacings from 35 to 300 μs (about 0.5 to 5 cm) [85].

For underwater echolocation, relating the target’s features to the structure of its echoes is more complicated. Except for the large impedance mismatch at the interface to the gas-filled swim bladder of fish, where glint-like reflections predominate [3,46,86,87], the impedance mismatch at the boundary between water and different solid materials is smaller, so the initial reflection is weaker [3]. Appreciable acoustic energy also enters the material to propagate inside the target and reflect off any internal impedance boundary. A variety of different behavioral experiments have examined the ability of bottlenose dolphins to perceive differences in targets that vary in size, material composition, thickness, and shape [3,14,27,46]. Because the targets have acoustic impedances closer to that of water, sound not only returns from features on the outside, such as surfaces and edges, but also penetrates inside to reflect from internal boundaries as the sound moves through the material. Particularly definitive experiments have used geometric shapes such as hollow metal cylinders with different diameters, wall materials, and wall-thicknesses suspended in the water in front of the animal [3,27,46]. Echoes from these objects consist of reflections from the front surface of the nearest wall and the front surface of the back wall, on the far side of the water-filled center, both of which are water-to-metal acoustic interfaces. These reflections are separated by the sound path across the cylinder’s hollow center. In addition, sound enters the wall material itself, travels inside the wall, and encounters the back boundary of each wall, which is the opposite metal-to-wall interface. Each boundary gives rise to an internal reflection that scatters back in the direction of the incident sound and eventually returns as one of several highlights in the echo as a whole. Further complications arise if the cylinder is not suspended perpendicular to the axis of ensonification, with its broadside to the dolphin. If it is tilted relative to the incident sound, each corner or distinct edge reflects its own mini-echo. The aggregate echo from the cylinders contains several closely-spaced glint reflections from the front and internal walls of the cylinders. The experimentally-manipulated diameter, material composition, and wall thickness determine the time separations among the reflections, which create highlights in the aggregate echo from the target. Most importantly, when expressed in terms of glint reflections, discrimination of wall-thickness and cylinder size by dolphins corresponds closely with discrimination of the spacing between reflective surfaces by bats [27, 46].

### Constraints on Modeling Biosonar

Several factors identified from experiments with bats, and in several instances corroborated from experiments with dolphins, stand out as distinct from conventional engineering methods for sound transduction, signal processing, and output display. These constraints are described below.

#### Biosonar sounds

Fig 1A illustrates the wideband clicks emitted by bottlenose dolphins (broadcast, left). The dolphin click is a sharp transient about 50 μs in duration containing frequencies from 40 to 150 kHz; due to the short duration, these frequencies appear as a continuous vertical stripe in the spectrogram (broadcast in Fig 1C). Fig 1D illustrates the FM chirp emitted by big brown bats (broadcast, left). The chirp is 3 ms long with frequencies dispersed in two prominent downward frequency sweeps, the 1^st^ harmonic (FM1) from 55 to 25 kHz and the 2^nd^ harmonic (FM2) from 90 to 50 kHz (Fig 1E, left). In spectrograms, the width of the dark ridges or stripes relates to the integration time used to compute the time-frequency slices of the signals (about 250 µs for the dolphin click, 300-350 μs for the bat chirp). These values were chosen to approximate the integration time of sound reception by the animals’ inner ears [3]. For the dolphin click, the spectrogram’s horizontal width (Fig 1C, left) reflects the time-smearing that occurs during calculation of the spectrogram slices. In the FM chirp, the frequencies are not instantaneous but are dispersed in time along the sweeps. At any moment, the frequency dwells slightly longer due to the finite sweep rate, so the spectrogram ridge is slightly wider than for the dolphin click, and the slices are calculated with a slightly longer integration time (Fig 1E, left). These factors influence both the conventional spectrograms used here for illustrative purposes and the SCAT model’s auditory spectrograms.

Nearly all of the roughly 1000 different species of echolocating bats emit FM signals ranging from 1 to 20 ms in duration [2,13,14]. Many of these species also emit constant frequency (CF) signals in conjunction with their FM signals. The CF components typically are either short in duration (1-5 ms) and capable of being omitted, or long (10-50 ms) and obligatory. The SCAT model addresses only processing of the wideband FM components. Moreover, although the FM sweeps are nearly always downsweeping, their shape varies from linear with frequency and time to exponential with frequency and time [13, 14]. The sounds transmitted by big brown bats have curvilinear sweeps (Fig 2) exponential with frequency and time but reasonably well approximated as logarithmic downsweeps or as hyperbolic downsweeps (*i.e*., upsweeps that are linear with period) [88]. Here, for simplicity we use logarithmic downsweeping FM signals as representative of big brown bats. The presence of the harmonics (FM1, FM2 in Fig 1E) is not a common feature of sonar or radar transmissions, but it is a natural property of the bat’s laryngeal sound production mechanism [2,15,16]. Although sound production by both bats and dolphins is driven pneumatically, differences between the bat’s laryngeal and the dolphin’s nonlaryngeal vocalizing structures ultimately reflect anatomical solutions that accommodate the acoustic impedance differences between air and water. Similarly, the radically different sound reception pathways used by bats through the external, middle, and inner ears and by dolphins through the lower-jaw and fat channels to the inner ear accommodate the acoustic properties of air and water [2,3,14,15]. In spite of these differences, after sound reception occurs, the inner ear and auditory systems of bats and dolphins share several response properties [27, 46]. For simplicity, we assume for our modeling purposes that they are amenable to the same computational approach.

**Fig 2.**
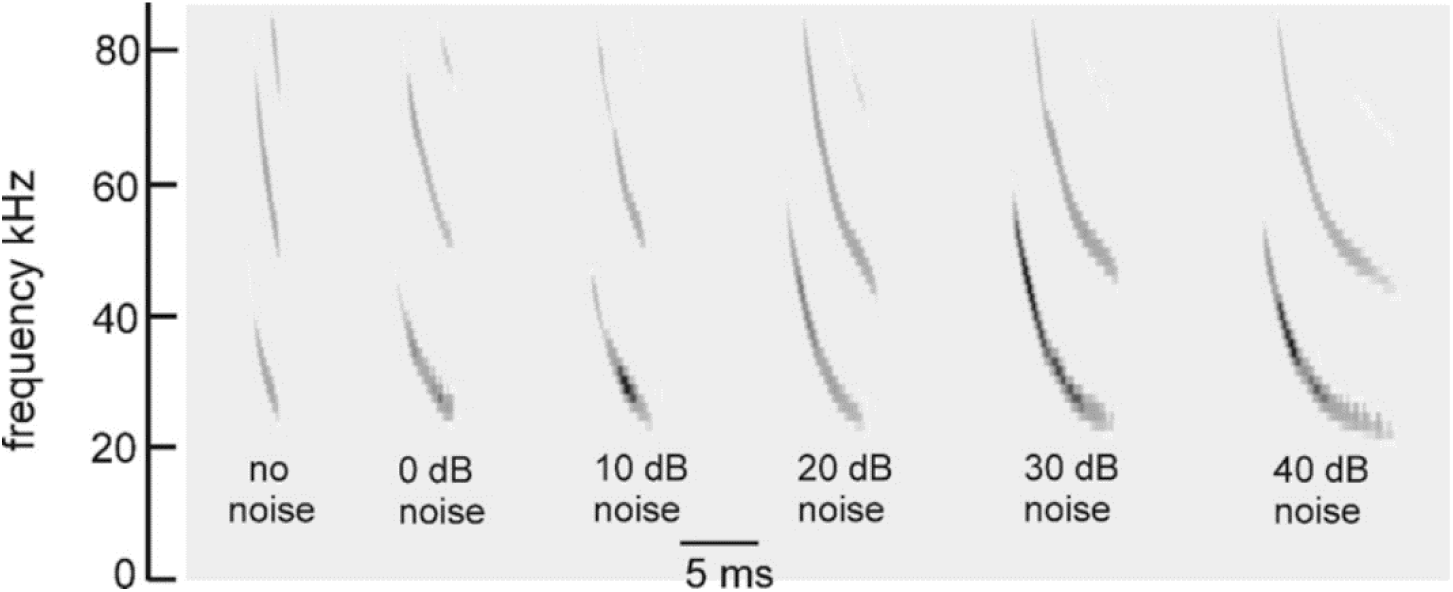
Time-frequency adaptability of FM bat biosonar sounds. FM pulses recorded from a big brown bat performing in psychophysical tests of virtual echo delay discrimination with electronically-generated echoes [38]. Increasing levels of wideband noise (X axis) were added to the echoes, which induced the bat to increase the duration of its broadcasts from 1-2 ms with no noise (far left) to 6-7 ms in intense noise (far right). Performance in the task nevertheless remained the same in terms of percentage error responses showing that echo reception and delay perception adapt to changing duration of signals.

#### Signal adaptability

An important constraint on echo processing is imposed by the *changes* the animals make in their outgoing signals. For both bats and dolphins, the most obvious change is in broadcast repetition-rates, or interpulse intervals, when approaching objects or pursuing prey [1–3,13–16]. They emit their biosonar sounds in trains, at repetition rates from 5-10 Hz to 150-200 Hz. This affects biosonar modeling because the initial SCAT model [22] and most similar models (see below) assume that each broadcast is followed by an undisturbed epoch of time when echoes return and are processed before the next sound is sent out to gather another set of echoes. Echoes can be matched to their corresponding broadcasts because no echoes of earlier broadcasts are intermingled with echoes of the current broadcast. However, for a big brown bat engaged in close maneuvering through complex surroundings, the premium on emitting sounds at short intervals to rapidly update its view of nearby collision hazards outweighs the need to wait for echoes to arrive from longer distances to fill in the whole scene [67, 68]. Overlap of echo streams from successive broadcasts creates ambiguity about matching echoes to their corresponding broadcasts, a serious problem for radar and sonar systems [36, 37]. As presented here, the model emits broadcasts and receives echoes using intervals long enough to fully segregate successive echo streams. Elsewhere, the question is addressed of how the big brown bat and the SCAT model adapt to what happens when new broadcasts are emitted while echoes of the previous broadcast still are coming back [64,67–69].

Changes in the broadcast waveform from one emission to the next also affect processing. Indeed, they are the primary computational motivation for using a time-frequency, spectrogram-like representation of signals [10]. Successive bottlenose dolphin clicks are relatively stable in both wave structure and duration (Fig 1A, left) [3,14,15]. They change systematically in amplitude according to the distance to the target, decreasing in sound pressure to compensate for the increasing strength of echoes at shorter ranges [14,15,48,89]. They are part of a larger pattern of changes in both broadcast amplitude and hearing sensitivity that stabilize echoes within the auditory system in the presence of changing echo strength in the external acoustic situation [49]. Other changes include shifts in the shape of the envelope of the spectrum to emphasize lower or higher frequencies while the dolphin is performing a task. In contrast, echolocating bats normally change the duration of their FM broadcasts over a wide range from milliseconds or tens of milliseconds down to 0.3-0.5 ms [1,2,13–16]. The big brown bat’s FM chirps change in duration considerably as the bat searches for, tracks, and intercepts a moving target or maneuvers in its surroundings [53,56,61,62,90]. Moreover, because the chirps are FM, changes in duration alter their time-frequency structure dramatically by changing the slope along the sweeps and shifting individual frequencies to different locations inside the overall duration [88]. The SCAT model is designed to account for echo-delay perception in the presence of these changes without requiring the internal computational structure to change each time the broadcast is changed. While performing the same delay discrimination task in different levels of masking noise, big brown bats adjust their broadcasts over a 6:1 span of durations (Fig 2) [38]. This variability exposes a crucial feature of wideband methods: The essence of wideband processing is retention of a template of the broadcast signal internal to the receiver for comparison with incoming echoes [36, 37]. The process of comparison yields an estimate of each echo’s delay. If all broadcasts have the same waveform, only one template needs to be stored. The same hard-wired receiver can operate on any broadcast because they all are the same. This might still be feasible if only a small number of discretely-different broadcasts are used; several templates can be stored alongside each other inside the receiver and called up one-at-a-time when any particular broadcast is selected. But, because the bat’s broadcast changes continuously along time-frequency dimensions depending on external acoustic conditions (Fig 2), with a continuous set of possible waveforms in the repertoire instead of one or even a few exemplars, the receiver has to update its template to take into account each new broadcast. Alternatively, it could accept the suboptimal use of a single echo-processing template for all broadcasts and perform less well for most of them. Here, the significance behind auditory reception of the broadcast as well as the echoes comes into play. The bat hears each of its broadcasts and builds a neural representation of echoes that incorporates the broadcast as a reference [2,91–93]. The simultaneous presence of a time-frequency representation of the broadcast followed by the same kind of time-frequency representation of echoes inside the auditory system provides a means for a tailored comparison of the individual broadcasts with the echoes, a criterion essential for wideband receiver design and built into the SCAT model and related models.

#### Glints and biosonar echoes

Echoes returning from individual targets of interest, such as insects (for bats) or fish (for dolphins), typically contain several discrete replicas of the incident broadcast reflected from the target’s prominent acoustic glints (wings, body, abdomen for insects [50, 53]; gas-filled swim bladder for fish [3,14,86,87]). The geometric arrangement of the glints is the wideband sonar equivalent of the object’s shape [70]. In one sense, glint structure also is a different kind of size—how much distance is spanned by the glints, not just how intense is the echo. Although overall echo amplitude depends on the target’s cross-section, which is a measure of size, the spatial spread of the glints interacts with amplitude to form perceptual size, which appears to correct glint representations for amplitude changes [77]. Fig 1 illustrates a series of echoes containing two glint reflections separated by different intervals from 9 to 700 μs. In water, for the dolphin clicks, these correspond to glint spacings from 0.75 to 60 cm. In air, for the bat chirps, these correspond to glint spacings from 0.15 to 12 cm.

For the dolphin clicks in Fig 1, the time waveforms of the resulting 2-glint echoes contain sharply-delineated time-domain highlights corresponding to the glint reflections (*e.g*., at 35 μs) that are clearly visible as separate peaks. Considering just the time waveforms, the separation (Δt) of the highlights characterizes the target’s shape acoustically (Fig 1A, 26 to 700 μs glint delays) [3,27,14,46,86,87]. These highlights are visibly separated even for glint-delay spacings as short as 26 μs and 35 μs. However, auditory reception transforms echoes onto auditory spectrograms, which obscures the discrete highlights for the two glints as events distributed in time unless they are far enough apart to evade the compressing effect of the integration-time of the inner-ear’s receptors (for dolphins, about 250 μs [3]). Following conversion into spectrograms, the highlights in time (Δt) (Fig 1A), appear as ripples in frequency (Δf) along the otherwise continuous vertical ridge in the spectrograms (Fig 1C). For example, in the 2-glint echo with the time separation (Δt) of 35 μs, the frequency spacing (Δf) of the ripples is 29 kHz, the inverse of the time separation. Only when the glint-delay separation is 300 μs or more do the two glint reflections of the dolphin click pull apart in the spectrograms to become visible as separate sounds (*i.e*., vertical ridges). The 250 μs integration-time determines the boundary between having the highlights visible at longer separations and having them merged at shorter separations. Given the evidence that the time separation of the glint reflections is an important perceptual dimension for dolphins [3,14,27,46,86,87], the migration of their representation from the time axis to the frequency axis has to be treated as a major constraint on biosonar modeling. Registration of the glints has to be sought in the echo spectrum instead of separate representation of each glint. Dolphins distinguish between targets by their glints, but do they perceive the time separation Δt or the frequency separation Δf (see below)? In experiments with passively presented, spectrally-shaped sounds, not actual echoes, the bottlenose dolphin discriminates between rippled and smooth wideband noise for ripple patterns with intervals (Δf) from 70 kHz down to 2-5 kHz, corresponding to glint delays (Δt) from 15 μs up to 200-500 μs [94].

Moreover, they classify ripple patterns into two subcategories—coarse ripples, or macro spectral patterns with nulls too far apart to appear as distinct ripples composed of multiple nulls (more than 15 kHz for Δf or less than 70 μs for Δt), and fine ripples or micro spectral patterns with enough adjacent nulls to be unmistakable as a distinct pattern (from 15 kHz down to 4 kHz for Δf or 70 to 250 μs for Δt [79]). In the spectrograms of Fig 1C, the pattern of ripples is easily visible for glint delays from 35 to 200 μs, which occupies mainly the micro spectral feature. In new experiments that present dolphins with actual echoes, not passively presented sounds, the boundary between time and spectral representation of two glints, and categorization of coarse and fine spectral ripples has been explicitly demonstrated [95, 96]. In bats, the conversion of glint time separations into spectral ripples is reversed so that the bat perceives the glints themselves in spatial terms [77,84,97].

Just as dolphins perceive the ripples in the spectra of echoes, so do big brown bats [85]. For the bat FM chirps (Fig 1D), the test broadcast (left) is 3 ms long. Over the range of glint spacings from 9 to 700 μs, the two glint reflections never become separate in any of the time waveforms at all. Instead, they overlap in time at all of the illustrated spacings. There are highlights (*i.e*., peaks) in the envelopes of the echoes (*e.g*., at 100 μs), but these are not the two glint reflections themselves but modulations of the time waveform caused by interference between the two reflections. The visible peaks and dips in the envelopes of the 2-glint chirp echoes occur at locations along the FM sweeps where the frequency corresponds to a peak or null in the spectrum at any given glint-delay spacing. This is more obvious in Fig 1E, where the locations of the nulls shift along the frequency axis of the spectrograms according to the delay separation. As an example, at 100 μs time separation (Δt), the nulls are spaced 10 kHz apart (Δf), which is the inverse of the time difference. The spread of frequencies in time along the FM sweeps and the separation of frequencies into two harmonic bands makes it harder to discern the regularity of the nulls as evenly-spaced ripples, however. In Fig 1E, not until the delay separation has increased to 100 μs do enough nulls fit into FM1 by itself for them to appear as ripples that follow along the FM sweep. Thus, the conditions for recognizing the pattern of the nulls is different for dolphin click echoes, with ripples more easily noticed at time separations as short as 25 μs, than for the fragmented bat chirps, where the ripples become noticeable only around 100 μs or more.

This simple visual analogy about the salience of the ripples reveals a factor that has to be addressed for biosonar modeling. For the dolphin click, the ripples in Fig 1C form a distinct pattern at glint delays as short as 35 and even 26 μs because the spectrograms all appear as a single, concentrated and uninterrupted vertical ridge. For the bat chirp, the dispersal of adjacent nulls both in time along the FM sweeps and in frequency into two separate harmonics poses a problem: A model that finds the ripples in the bat echoes has to sort out the nulls not only at different *frequencies* but at different *times*. In the SCAT model and related examples [12,22,23] and in FM radar generally [36, 37], this difficulty is solved by dechirping the FM sweeps so their frequencies are compressed in time to remove the tilt of the sweeps and counteract the dispersion of successive frequencies along the sweeps. This effectively turns the sweeps into a vertical ridge in spectrograms, much like the dolphin clicks (Fig 1). This is especially important in view of the wide range of adaptability in broadcasts used by bats (Fig 2). Sounds and their echoes that have different distributions of frequencies across time have to be converted into a common format to achieve the same delay acuity in spite of different FM sweep rates used by bats. Information about the timing of the glints, which is the acoustic equivalent of target shape, is folded into the spectrograms and appears along its frequency dimension as ripples instead of highlights in time. The glint highlights still represent the object’s shape, even though they have disappeared into the ripples. However, the FM sweeps spread the ripples across the time axis of the spectrograms, too, and dechirping aligns them for successful extraction. Following dechirping, recovering the glint-based target shape requires reversing the transformation of the interfering glint reflections back into the actual timing of the glints so they can be reattached to target range. This is the purpose of the SCAT model.

## Methods

### Original SCAT model

The SCAT model [22] is a time-frequency method for processing sonar echoes [34, 36]. It belongs to a family of time-frequency approaches to signal analysis and processing that have wide applications [34,98–100]. For creating spectrograms, the model adopts a bank of parallel bandpass filters tuned to different frequencies to mimic sound reception by the inner ear [35, 93]. The original version of the SCAT model [22] had two computational components. The first component was Spectrogram Correlation—a system for forming pulse and echo spectrograms and then determining echo delay from the time that elapses between the broadcast and the echo at each frequency. The utility of spectrogram correlation for determining echo delay has been demonstrated in a number of different implementations of the model [23,101–104]. Simulations of delay estimation show that its accuracy is comparable to matched filtering [24]. The second component of the model was Spectrogram Transformation—a method for converting the pattern of interference peaks and nulls across frequencies into a reconstruction of the glint-delays themselves [22]. It was inspired by work that demonstrated the utility of spectrograms of echoes for representing details of targets [105]. The first version was a simple analysis-and-synthesis approach: The FM signal was broken down into 81 parallel frequency channels to mark the time-of-occurrence of each frequency in the broadcast and then in the echo, to be used for spectrogram correlation. At the same time, the time-marking event in each frequency channel was used to trigger a rapidly-decaying burst of oscillations that corresponded to the impulse response of each frequency channel. For each channel, the burst consisted of a series of cycles at the frequency of that channel that declined in amplitude over about ten cycles. Because all of these bursts began in phase with each other (they were triggered by the instantaneous occurrence of their corresponding frequencies, but dechirped, not with their time offsets in the FM sweeps), they were added together as synchronized parallel time waveforms to form the aggregate impulse response of the parallel frequency channels as a complete system. The resulting signal amounts to the crosscorrelation function of the echo, *i.e*., the output of a matched filter [23, 24].

This early version incorporated the amplitudes at different frequencies (the nulls and peaks in the spectrograms) as the starting amplitudes of the individual frequency-tuned bursts. The glint structure of the echo emerged as a consequence of some of the side peaks in the composite, summed impulse response not canceling out when the bursts were summed. These side peaks mark the glint delays themselves. Subsequent iterations of the model adopted several alternative methods for reading the glint interference pattern out of the spectrograms to estimate the glint delay spacing underlying the interference pattern [101–104]. Subsequent behavioral and neurophysiological research has refined our understanding of auditory mechanisms in bats, focusing attention particularly on how peaks and nulls in the echo spectrogram are encoded in neural responses [106–108]. The new version of the model described here incorporates these developments, in particular shifting the mechanism for detecting and using the nulls into the frequency domain and making the transformation back into the glint delays by exploiting the inverse relation between the delays in time and the null spacings in frequencies. The underlying principle is a set of spectrogram templates for different ripple patterns that are matched to specific echoes to estimate glint delays. A version of the model using templates has proven effective [25], but in the new model the templates remain more dispersed computationally in keeping with auditory frequency representations in bats [18–21,107,109–113].

### Block diagram of the SCAT Model

Fig 3 is a block diagram of the present version of the SCAT model. In this and subsequent examples, the signals being processed consist of a single prototypical broadcast followed by several echoes. The original version of the model used only single FM pulse-echo pairs [22]; here, we use multiple echoes following each single broadcast as a convenience for batch processing of multiple echoes to illustrate the model’s output for different echoes in a single display format. As shown schematically here, the input is an FM chirp at top left followed by an echo at delay t that contains two glints separated by Δt. Reception occurs through a bank of parallel bandpass filter—161 gammatone filters, center frequencies from 20 to 100 kHz spaced at 0.5 kHz intervals, with equivalent rectangular bandwidths, or ERBs, of 4 kHz. These parameters were chosen to approximate frequency tuning sharpness in big brown bats [109–113]. Gammatone filters are commonly used for modeling auditory receptor tuning [35]. Furthermore, the transduction units are very complicated, creating a hybrid of spectrogram and waveform representation that figures prominently in auditory perception, including SCAT (see below) [114, 115]. The filterbank that we use here reproduces an outstanding feature of auditory transduction—many, closely spaced center frequencies and a high degree of overlap between adjacent filter frequency responses (almost 90% of each filter’s band relative to the adjacent filter’s band). In conventional terms, this looks like oversampling in frequency; an engineered filterbank might have filters spaced apart so their ERB band edges just touch rather than overlap substantially. The simple assumption is that frequency is the dimension being sampled, and the filters are spaced apart to cover the desired frequency band without too many, unnecessary filters. A time history for the amplitude of individual frequencies emerges from the bandpass filters. In normal circumstances the filter outputs would be full-wave-rectified and then subjected to suitable lowpass smoothing to set the scale for following variations of frequency over time. As described below, the auditory system’s filterbank creates a richer content for auditory spectrograms than just sampling of frequencies. First, the filter outputs are half-wave-rectified, which is not conventional: Why discard half of the signal’s energy carried by the negative-going cycle peaks before lowpass smoothing? Second, the cutoff frequency for lowpass smoothing is high enough to allow details of the cycle-by-cycle phase in the low-frequency segment of the waveform to slip past and become an integral part of the overall time-frequency representation. How this content is used depends on the algorithms that follow the bandpass filters, which we take up before returning to the content.

**Fig 3.**
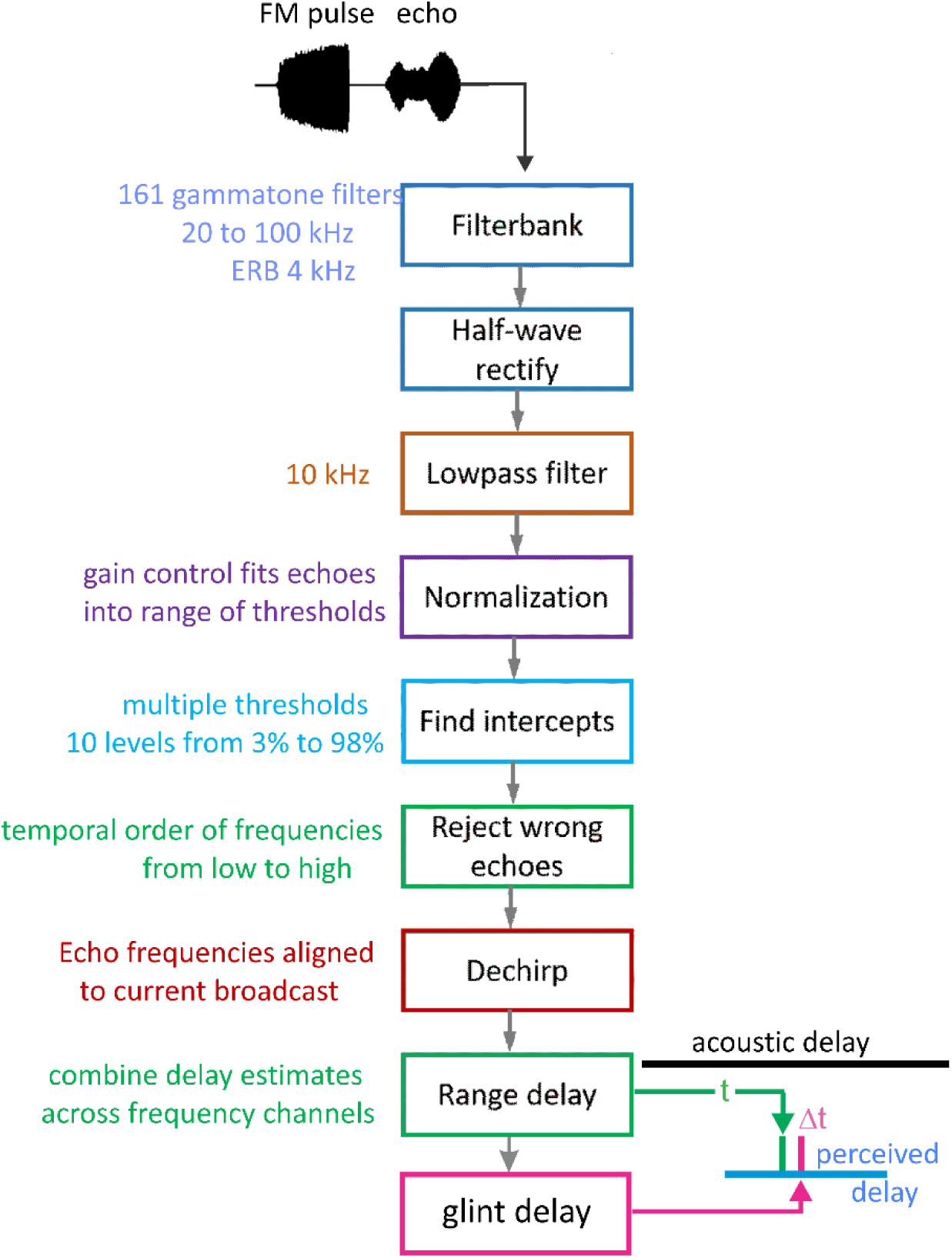
Block diagram of SCAT model. The input is an FM pulse followed by an echo at delay (t) with two glint reflections separared by (Δt). These signals are segmented into parallel frequency channels by 161 bandpass filters (4^th^ order gammatone filters, center frequencies from 20 to 100 kHz, equivalent rectangular bandwidth, ERB, 4 kHz), then half-wave rectified and lowpass filtered (10 kHz cutoff). Outputs are then normalized to fit into 10 equally spacedthreshold levels (3% to 98% full-scale). Processing started at the lowest frequencies in the broadcast in bats 23-25 kHz); if these particular frequencies are absent, then echoes are rejected and processing is not started [64]. To remove the slope of the FM sweeps, echo frequencies are dechirped on the time-frequency plane by setting zero origin of time for individual bandpass frequency channels to threshold times of the pulse, resulting in corresponding thresholds of echoes arrayed at longer delay corresponding to target range. These threshold times are then analyzed by two parallel pathways—range delay from total echo delay (t) determined by time between pulse and echo thresholds combined across all broadcast frequencies, and glint delay (Δt) from spectral notch detection followed by extraction of time spacing for glint reflections before merging of range delay and glint spacing onto a common perceived delay scale (blue, at output).

Subsequent SCAT processing takes place separately in the individual frequency-tuned channels of the time-frequency representation created by the bandpass filters. The transduction processing stages are half-wave rectification followed by lowpass filtering at 10 kHz to partially smooth the envelopes of the filtered, rectified signals without destroying all phase information. The choice of 10 kHz for the lowpass cutoff comes from simulations of echo delay accuracy using bat-like FM broadcasts followed by echoes at different signal-to-noise ratios [24]. Big brown bats can detect changes in echo delay of fractions of a microsecond at moderate to high signal-to-noise ratios [29, 39]. These simulations reveal that delay accuracy determined by spectrogram correlation fails to achieve the bat’s performance if lowpass filtering of bandpass filter outputs uses cutoff frequencies below 10 kHz. Following lowpass filtering and half-wave-rectification, the amplitudes of the signals are normalized to impose a gain control that fits the full-scale amplitude of each broadcast into the dynamic range of ten equally spaced thresholds from 3% to 98% of the full-scale amplitude range. This thresholding scale is derived from neural responses recorded from the brainstem and midbrain auditory pathway in big brown bats: Response thresholds vary from close to the bat’s hearing thresholds (∼5-15 dB SPL) to as much as 50-60 dB above hearing thresholds across ultrasonic frequencies [111–113]. Such a wide dynamic range refers to sound pressures of stimuli in neurophysiological experiments. It does not consider the awake, active big brown bat’s peripheral gain control from middle ear muscle contractions synchronized to vocalization [17,18,47]. These reduce reception sensitivity by a minimum of 30-35 dB, squeezing the directly-received amplitude of broadcasts into a narrower dynamic range that tops out at 30-35 dB above detection thresholds at different frequencies. Then, graded relaxation of these same muscles ramps up sensitivity at the same rate that echo amplitude declines with target distance. The time course of relaxation tracks the component of echo amplitude related to distance to keep echoes inside the dynamic range from detection threshold to about 30-35 dB sensation level [47]. We mimic this compression of the effective dynamic range of echolocation with gain normalization followed by ten threshold crossing levels. They are a simplified way to establish quantification of amplitudes for the model’s simulation of echolocation. By this stage, the model in Fig 3 has created 161 parallel time-frequency-amplitude channels that each carry a single time value for both the pulse and also echo threshold crossings at each frequency in each sound. Next, the representation of each echo is checked to determine if the lowest frequencies in the broadcast are present. For big brown bats, frequencies of 25-30 kHz are required to initiate processing for the delay of any echo [64]. If these frequencies are absent, the echo is rejected. If the lowest frequencies are present, delay processing begins at these lowest frequencies and proceeds in order from lowest to highest. In Fig 3, to dechirp the broadcast-echo stream, the time-of-occurrence of each broadcast frequency at each threshold level is set as time zero for estimating echo delay. Dechirping removes the time-frequency slope of the FM sweeps using the broadcast as the reference for each frequency’s delay estimate, thus aligning all broadcast frequencies to the same zero-delay origin for determining echo delay.

To determine range (echo) delay, the times-of-occurrence of corresponding frequencies are registered at the ten threshold levels for each broadcast and its echoes in each frequency channel (Fig 3). The time intervals that elapse between each threshold event at each frequency in the broadcast and the corresponding frequencies and thresholds in the echo are accumulated across frequencies to estimate overall echo delay (t). This combined delay estimate represents the output of the spectrogram correlation process of SCAT (green arrow and delay in Fig 3). Such a simple range-delay estimate occurs only for a single, 1-glint target returning an echo that is just a delayed replica of the broadcast. Its echo spectrum is flat, unaffected by long-range or off-side lowpass filtering, and it is not affected by the object’s shape.

For complex objects consisting of several glints, the returning echo contains several mini-replicas of the incident broadcast that represent the placement of the glints in range by their individual delays. Recovering those delays is the purpose of spectrogram transformation. Because the interglint delays are shorter than the integration time of the gammatone filters in the filterbank, the glint reflections merge together into a composite echo that represents range delay by its overall delay and the object’s glints by its spectrum. In a stage of SCAT processing parallel to estimating glint delay (t), the glint delay (Δt) is extracted by estimating and then inverting the frequency spacing of interference peaks and nulls (Δf) that is reciprocally related to the desired time separations (Δt). To illustrate this process, the schematic echo in Fig 3 contains two such glints. The small glint delay spacing (Δt) determined from the interference spectrum is directed onto the same perceptual delay axis used to display the range delay (t) (in Fig 3, green mark for t, red mark for Δt). Big brown bats perceive both types of delays on a common psychological or cognitive scale, indicating that target location and shape are both “spatialized” from their acoustic origins when translated into perception [25, 77].

### Structure of algorithms in the SCAT model

Fig 4 illustrates the sequence of representations developed at each stage of the SCAT model for an FM pulse followed by an echo arriving at a delay of 6 ms (t), corresponding to a range of 1 m. First, in Fig 4A, each sound is transformed into its spectrogram by the filterbank and displayed along the resulting vertical frequency axis. Again, this target contains two glints, with the second glint 17 mm farther away than the first glint, resulting in two reflections 100 μs apart (Δt). The interference spectrum of the echo thus has peaks and intervening nulls separated by 10 kHz intervals (Δf). These appear as a regular pattern of ripples along the two harmonic ridges of the spectrogram (Fig 4A). The individual frequency slices in the computed spectrogram contain short, half-wave-rectified and partially-smoothed bursts emerging from the bandpass filters. They are subjected to ten threshold crossings in parallel, scaled so the broadcast occupies all ten threshold levels (3% to 98% full scale). Three representative threshold crossings are shown as blue dots in Fig 4A, connected by horizontal lines that indicate their corresponding range delays. For example, there is a blue mark in the broadcast at 40 kHz in FM1 followed by a similar blue mark at the same frequency in the echo. The horizontal blue arrow connecting them is the range delay of 6 ms recorded for the frequency of 40 kHz. A similar pair of lighter-blue dots marks the broadcast and the echo at 70 kHz in FM2. Again, the length of the lighter-blue horizontal arrow marks the range delay of 6 ms at 70 kHz. There is also one threshold crossing marked by blue dots at 34 kHz in FM1. However, there is an interference null in the echo spectrogram at this frequency, while both 40 and 70 kHz correspond to peaks in the echo spectrogram. The blue dot marking the time-of-occurrence of 34 kHz in the echo is not aligned on top of the spectrogram ridge; instead, it is displaced to the right (*i.e*., to a longer delay) by nearly one millisecond. This sideways overestimation of echo delay (horizontal blue arrow at 34 kHz) is caused by the lower amplitude of the spectral null at 34 kHz relative to the flanking peaks at 30 and 40 kHz. The anomalously-long delay for registering range delay at 34 kHz is caused by *amplitude-latency trading* (amplitude-latency trading; purple arrow and lines in Fig 4A), a phenomenon of auditory sensory transduction that retards the time-of-occurrence, or latency, of neural responses that mark the occurrence of frequencies with lower amplitudes. Measured behaviorally from the perceived lengthening of echo delay, amplitude-latency trading is 15-25 µs of added latency for each decibel of echo attenuation [29, 77]. Measured from neural responses, amplitude-latency trading varies from 15-200 μs for every decibel of attenuation [77,109,116–120]. We use 25 µs per decibel of attenuation in SCAT because the intention of the model is to reconstruct the bat’s perceptions. The latency shifts come from both the slight retardation of threshold crossings that occur when amplitude decreases, a straightforward waveform phenomenon, and changes in the combination of excitation and inhibition that occur at synapses between neurons in the auditory pathway, a physiological phenomenon [115, 116]. The lengthened latency at 34 kHz in Fig 4A reflects the lower amplitude there; the diagram illustrates how latency becomes the surrogate of amplitude for purposes of SCAT computations. It transposes numerical values of lower echo amplitude at specific frequencies into longer response latencies at those frequencies, which can be processed using differences in the timing of neural responses between neurons—a hallmark of auditory processing [35].

**Fig 4.**
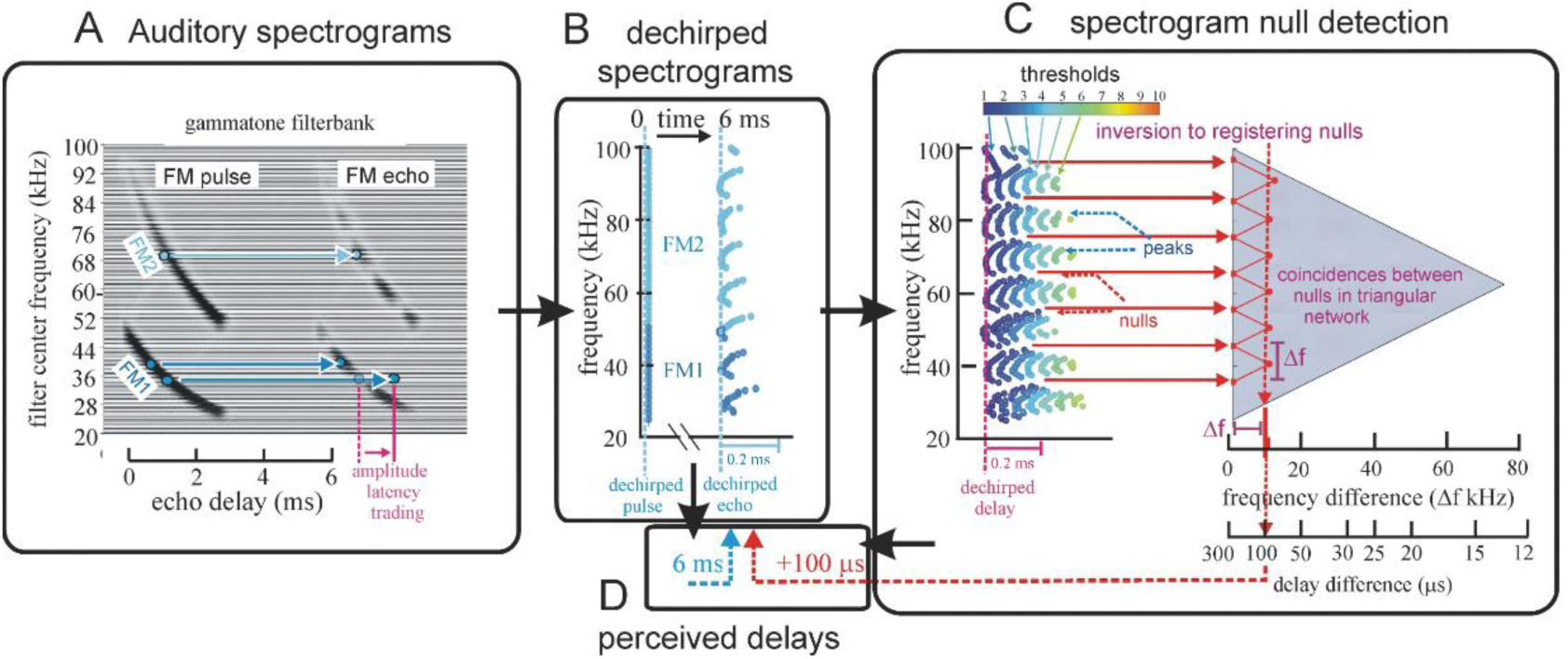
Algorithms in SCAT model for processing of FM pulses. (**A**) Initial reception of biosonar broadcast and returning echo. The FM pulse contains two harmonic sweeps FM1, FM2) and is followed 6 ms later by 100-μs two-glint FM echoes containing multiple interference nulls at frequencies 10 kHz apart (reciprocal of 100-μs glint spacing) caused by overlapping glint reflections. The model computes spectrograms with 161 parallel gammatone bandpass filters tuned to center frequencies of 20-100 kHz. Filter outputs are half-wave rectified, lowpass-filtered at 10 kHz, and thresholded with 10 amplitude levels. In each channel, the time that elapses between crossings from the same threshold in the chirp and the echo (horizontal arrows, blue circles) marks delay measurements. At frequencies where echo and broadcast spectrograms have the same amplitude, crossings register echo delay from times-of-occurrence accurately. If the echo is weaker, crossings across all frequencies are later due to amplitude-latency trading, and the delay estimate is longer. At frequencies with interference nulls, echo amplitude is locally weaker than at surrounding peak frequencies. At nulls, crossing is later due to amplitude-latency trading (red time offset). (**B**) Delay is estimated frequency-by-frequency using the pulse-to-echo elapsed times. Threshold crossings in the pulse mark the start of the delay estimate (time zero). Across frequencies, time intervals between pulse and echo crossings are aligned on pulse thresholds at zero time, which dechirps the FM sweeps to make vertical row of time marks. Only crossings from one threshold are shown in A and B to illustrate time marks (blue circles). Time of echo thresholds creates a similar vertical row of marks in each channel, modified by amplitude-latency trading. The threshold marks at the nulls occur later (to the right), causing the dechirped echo to have a scalloped appearance. The leftmost, leading edge of the curved thresholds marks the echo’s 6 ms delay. (**C**) Inversion of representation from echo amplitudes across frequencies to echo nulls across frequencies. Close-up view on left shows dechirped echo threshold marks for 6-7 activated thresholds (#1 up to 7 out of 10 levels on color bar). This representation is densely populated, coming from all the time-frequency values that exceed different threshold levels spread across about 0.2 ms from lowest threshold (dark blue) to highest threshold (light green). Spectrogram amplitudes track along the thresholds as clusters where they exceed thresholds; nulls have marks only at lowest thresholds because their amplitudes are weak (dark blues), and the track of these threshold events is curved, extending to longer times (rightward) due to amplitude-latency trading. Between peaks, where the thresholds are clustered, there are voids at the center of nulls where none of the thresholds are crossed. Locations of nulls are extracted from scalloped pattern of thresholds across frequencies. These longer latencies and the voids are transformed into representation of the nulls (red horizontal arrows), which is a sparse representation due to inversion from the dense representation of amplitudes that exceed thresholds. This peak-to-null inversion is key to subsequent processing: The late or absent responses at nulls trigger new responses that progress to next stage, a triangular network of model neurons that registers the nulls and connects adjacent nulls with triangular connections at different frequency spacings set by frequency separation between filters in the filterbank. Frequencies of nulls are marked in red dots at the left of the triangular network in 0.5 kHz frequency steps, the same as the gammatone filters. The frequency differences between nulls form a zig-zag pattern of coincidence responses that register the frequency spacing of adjacent nulls (Δf) by their right-most triangular apex points in the zig-zag. These points are read out of the triangular network by the vertical alignment of the triangular apex points (vertical dashed red arrow) projected down onto the horizontal frequency difference scale. This yields an estimate for the average frequency spacing of the nulls (Δf = 10 kHz). The corresponding reciprocal of 100 μs is registered on the horizontal delay difference scale and the spacing of the glint reflections in the echo. (**D**) The 100-μs glint delay estimate from the triangular network in C (red arrow) is attached to the 6 ms overall echo delay estimate from the thresholds in B (blue arrow) to form an image of the target’s range and shape.

The algorithms used to determine the echo’s range delay are explained in Fig 4B. They dechirp the broadcast and stack all of the broadcast’s threshold crossings at time zero (threshold level #3 is illustrated by blue dots; dark blue for frequencies of 25-50 kHz in FM1, light blue for frequencies of 50-100 kHz in FM2). Then, range-delay processing is referred to time zero in each frequency band and at each threshold band. The overall range delay is determined by combining the time intervals between broadcast and echo threshold crossings at each frequency in first threshold level. In Fig 4B, the pattern of threshold-times for the broadcast is a vertical row of dots; this is the effect of dechirping. The pattern of threshold-times for the echo is not so uniform because the local reductions in amplitude at frequencies of the nulls causes amplitude-latency trading, which moves them to longer latencies, to the right, at the nulls compared to the intervening peaks. The scalloped appearance of the dechirped echo spectrogram provides the raw material for finding the nulls, but it also complicates the process of determining the range delay. Two aspects of this process deserve mention: First, the overall echo delay of 6 ms is shown as corresponding to the leftmost (*i.e*., earlier) leading edge of the repetitively-curved threshold-crossing events evoked by the echo. These early threshold marks represent the most valid source of range delay information because they are relatively unaffected by variations in amplitudes across the echo spectrogram, particularly the nulls (Fig 4A). The longer threshold marks are discounted for determining delay because they are retarded due to amplitude-latency trading. They are not without value, however, because they show where the nulls are located, which brings up the second aspect of the computations illustrated in Fig 4B. Here, the echo contains two glints 100 μs apart (Δt), putting interference peaks and their intervening nulls in the spectrogram 10 kHz apart (Δf). The salient feature of the dechirped echo in Fig 4B, which comes from the interference between the glint reflections, is the repetitive, curled pattern of peaks and nulls of the threshold crossings moving upward at different frequencies. The interference *peaks* are shown at the leftmost (*i.e*., earlier) blue dots (at frequencies of 30, 40, 50, 60, 70 80, and 90 kHz), while the dots curve to the right (*i.e*., later) as the lower amplitude of the *nulls* induces progressively more amplitude-latency trading, eventually with no threshold crossing at the frequency of the individual nulls themselves. The resulting scalloped appearance of the dechirped echo illustrates the value of amplitude-latency trading as an unconventional computational strategy; it renders the locations of the nulls easily recognizable just from the timing of the threshold crossings by the rightward curves and the holes in the pattern.

Fig 4C shows all ten threshold levels used to represent the dechirped echo. Six of the full ten thresholds are included as colored dots for the dechirped echo (horizontal color bar labels thresholds from 1 to 10). At the echo’s spectral peaks, the threshold crossings stretch over a time span of about 0.2 ms as they follow the amplitude envelopes of the half-wave-rectified, lowpass filtered outputs of the bandpass filters. Their spread reflects the role of signal amplitude in determining when thresholds at different levels are crossed. The nulls that are interspersed between the peaks are recognizable by the longer latencies of the threshold events, which give the plot its characteristic scalloped appearance. The centers of the nulls are voids where none of the thresholds are crossed. At this stage, the SCAT model adopts a process previously identified in neurophysiological experiments [106, 107] whereby the dense, multidimensional representation illustrated by the threshold events in Fig 4C is converted into a sparse representation of the frequencies of the nulls (horizontal red arrows in Fig 4C) [11]. The frequencies where nulls are registered form entries on a scale at the left side of a triangular network of coincidence-detecting connections. This triangular network determines the frequency spacing of the nulls from the diagonal connections on the surface of the triangle that link individual null frequencies by a zig-zag line (red). The frequency spacing is represented by how far into the triangular network from left to right the apex points of the coincidences are located. In this example, the frequency spacing of the nulls is 10 kHz, and the coincidences are aligned vertically on the column of points where the diagonal lines correspond to 10 kHz. This spacing is converted into its reciprocal, 100 μs, to display the glint spacing itself. In Fig 4D, the 100-μs glint spacing is dragged from the triangular network to be attached to the previously-determined range delay of 6 ms and registered as a second glint reflection 100 μs later. Formation of this composite but nevertheless all-time-valued image is the purpose of the model.

### Content of SCAT auditory spectrograms

A key feature of SCAT processing is exploiting fine structure in the time-frequency representation derived from the bandpass filterbank. Spectrograms ordinarily are made by passing a series of short-term Fourier transforms along the time axis of the signal to be analyzed. The resulting segments of the signal’s spectrum are stitched together to trace the evolution of frequencies over time [34,98,105]. The display’s resolution of frequency and time is determined by the size of the time window used for the individual segments, although it is common to have adjacent segments overlap, usually by 50%, to improve the smoothness of the frequency traces. Each time-by-frequency pixel of a spectrogram display is unitary; fine details of the waveform on a scale smaller than the pixel are lost because the amplitude of the spectrum is expressed by squared magnitude values. Auditory spectrograms are made using a bank of parallel bandpass filters designed to mimic the sharpness of tuning for inner-ear receptors and their associated afferent auditory nerve fibers [35]. Even the amount of frequency tuning of each frequency channel is a complicated matter, however, because the “receptor” is a group of hair cells that are mechanically yoked together to form an active mechanism with gain control and amplification quite unlike any familiar engineered device [114]. Then, the dynamics of auditory nerve excitation transforms the time-series signals passing through the bandpass filters, with their half-wave-rectification and lowpass smoothing, into a curious hybrid of spectrogram and time-domain representation [115]. Subsequent levels of auditory processing spread the features of this new representation across a variety of neural response patterns that are selective for sound frequency, duration, delay, and spectral profile [93, 116].

Fig 5 illustrates the richness of the content of auditory spectrograms [35] in the context of SCAT. First, Fig 5A shows the frequency response curves for a subset of 32 bandpass filters out of the total number of 161 filters in Fig 4. The center frequencies are 2.5 kHz apart instead of 0.5 kHz, but even with this subset, the degree of overlap is almost complete. Any two adjacent filters have responses only slightly shifted apart, in contrast to the overlap of the curves. This points to a different description for the purpose of the bandpass filters. Instead of merely sampling the frequency axis in order to determine the spectrum of wideband echoes, the much higher density and close packing of the auditory bandpass filters is also involved in registering details of the time-series waveform of echoes. Fig 5B shows a subset of the filter outputs for the same FM broadcast and echo illustrated in Fig 4, including the half-wave-rectification and lowpass smoothing. The timing of the threshold crossings for threshold #4 is shown by red dots. What stands out in the waveforms is the prominence of the cycle peaks at the lower frequencies of the 1^st^ harmonic, FM1, compared to the smooth envelopes coming out of the higher frequencies of the 2^nd^ harmonic, FM2. The echo portrayed in Fig 5B has two glint reflections separated by 100 μs, and the frequencies of the interference nulls are 10 kHz apart. The plots are dechirped to align the threshold crossings for the broadcast at time zero for purposes of determining echo delay. The nulls appear as excursions of the threshold crossings for the echo waveforms to the right, with amplitude-latency trading here just due to the displacement of the crossings. Fig 5C shows heatmaps for the waveforms emerging from the broadcast to illustrate how the SCAT spectrograms go beyond a conventional spectrogram to include the structure of the signals at lower frequencies. In human hearing [35], this structure forms the basis for perceiving the pitch of complex sounds, including vowels in speech and notes in music. Echolocating bats and dolphins perceive changes in the phase of biosonar echoes as corresponding changes in echo delay [29–32,121]. The traces of half-wave-rectified cycles passing by the lowpass smoothing suggest how the SCAT model can be used to explain this capability. Overlap of the bandpass filter responses ensures that the signals emerging from adjacent bandpass channels are very similar, enough so that tracking small differences in the filtered output waveforms provides for sensing changes in phase between channels, which is effectively the local slope of the sweep for FM signals.

**Fig 5.**
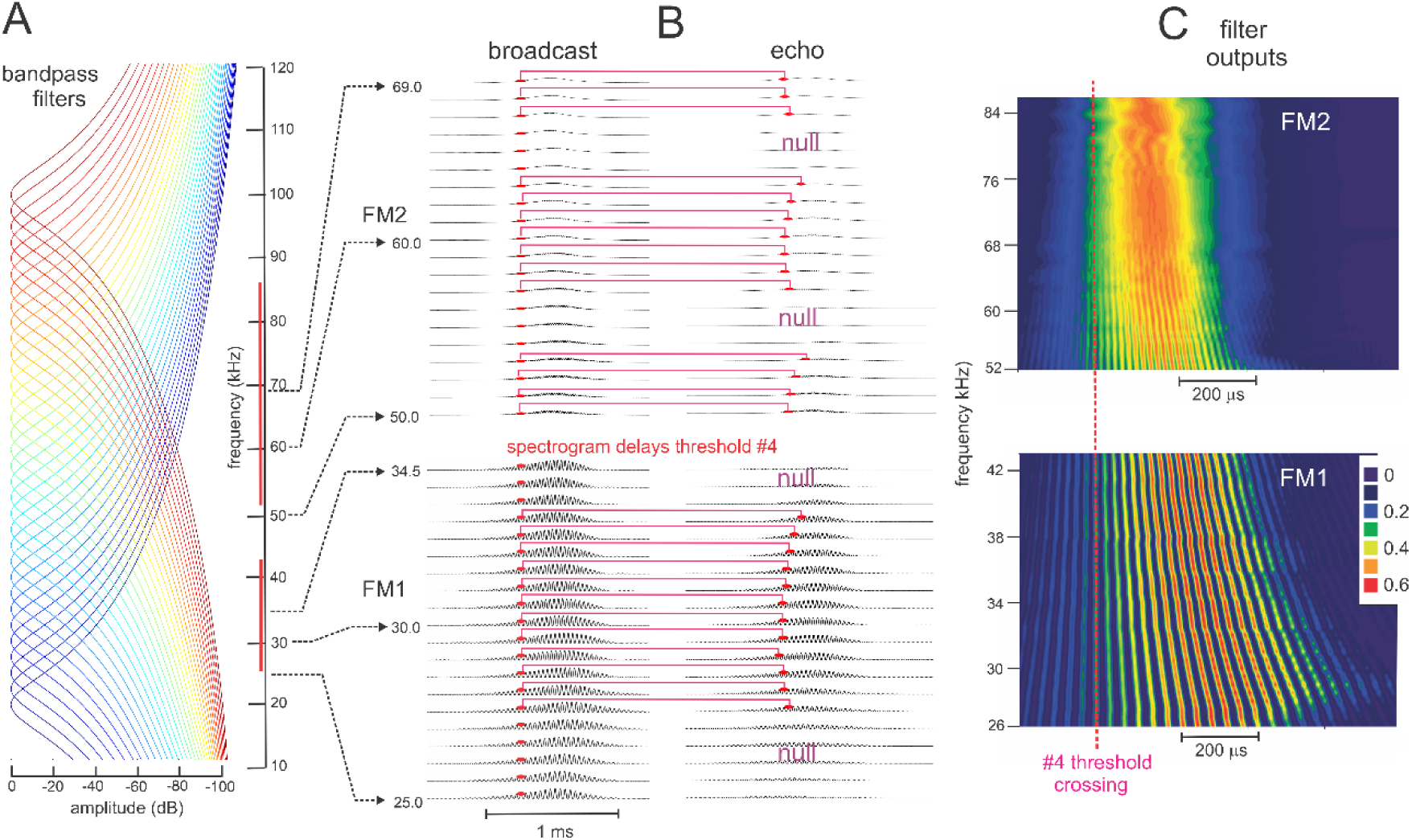
Fine structure of SCAT spectrograms. (**A**) Frequency response curves for subset of 32 parallel bandpass filters from the SCAT input system (frequencies from 20 to 100 kHz at 2.5 kHz intervals, a subset of the full filterbank of 161 filters at 0.5 kHz intervals). (**B**) Dechirped filter outputs (half-wave-rectified, 10 kHz lowpass filtered) for FM broadcast and 100 μs two-glint echo at selected FM1 and FM2 frequencies. Crossings for threshold #4 shown as red dots; dechirping aligns threshold crossings in broadcast at time zero. Horizontal red lines show spectrogram delays in each frequency channel. Curved appearance illustrates effect of amplitude-latency trading (ALT) on individual delay values near nulls in echo spectrum. (**C**) Heat maps show sequence of peaks in filter outputs for 1^st^ harmonic bands of 26-43 kHz in FM1 and corresponding 2^nd^ harmonic frequency bands of 52-84 kHz in FM2 (marked by vertical red lines in A). Both plots are dechirped by alignment to #4 threshold crossings. The seeming oversampling of frequencies by closely-spaced bandpass filters in A reveals a detailed representation of the cycle-by-cycle structure for adjacent frequency bands in FM1 in spite of 10 kHz lowpass filtering, while this structure is lost at FM2 frequencies, especially above 60 kHz.

### Software of SCAT model system

The datasets developed along the SCAT model’s processing cascade (block diagram in Fig 3) have been explained in Fig 4. These illustrations are intended to assist the user in running the model and assessing how each stage’s routines are working. Fig 6 shows a close-up view of one particularly critical stage of processing—the enhancement of threshold-crossing timing that marks the frequencies of the interference nulls used to estimate the glint-delay values from Fig 4C. The problem addressed at this stage is to find the frequencies of the nulls from the longer threshold-crossing times and the loss of threshold detections at higher threshold levels in the vicinity of the nulls using the curved, scalloped shape of the rows of dot representing these time marks. Amplitude-latency trading increases the time disparities at null frequencies and makes estimation of these frequencies more precise (red horizontal arrows in Fig 4C).

**Fig 6.**
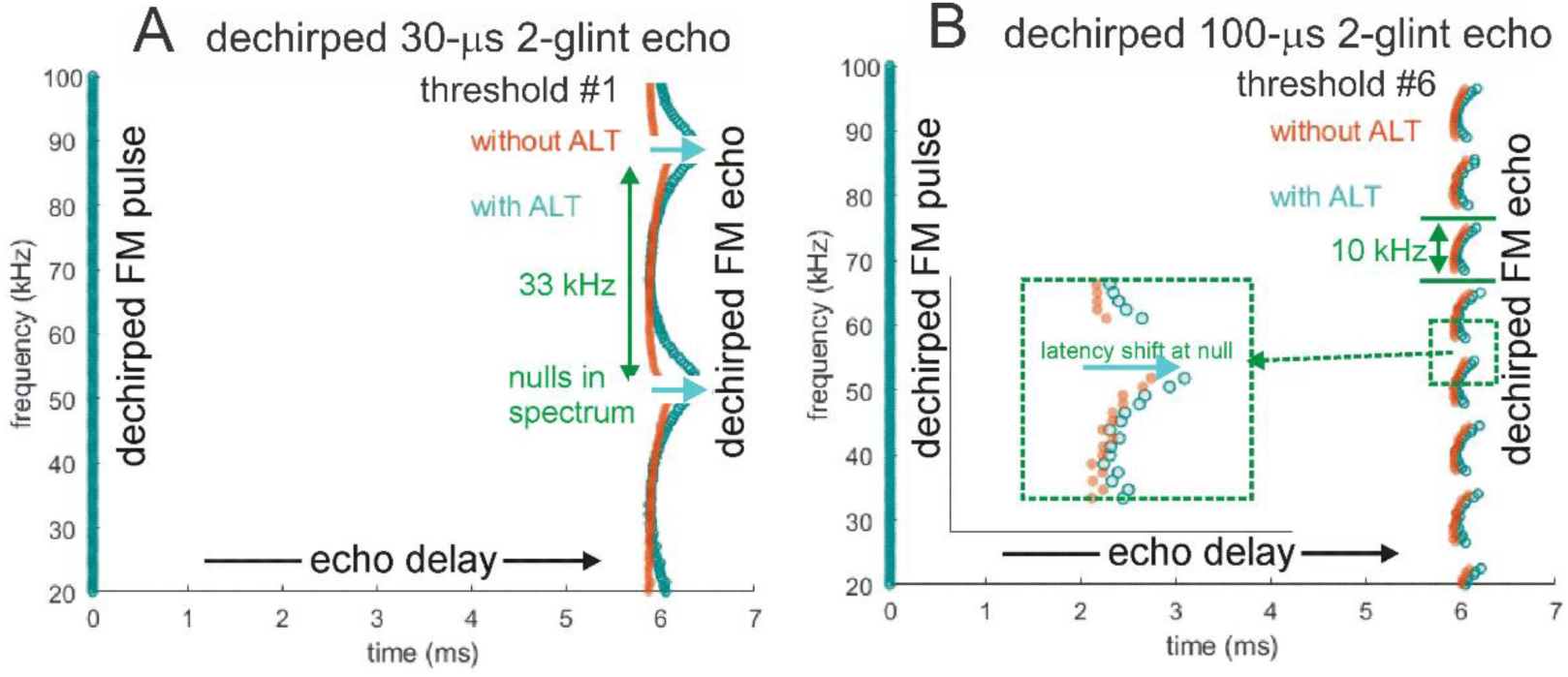
Enhanced registration of nulls by amplitude-latency trading. Dechirped SCAT spectrograms for FM pulse and two-glint echoes with (**A**) 30 µs glint delay separation and (**B**) 100µs glint delay separation. Threshold crossings (threshold #3) are marked by circles (orange for no amplitude-latency trading (*i.e*., just threshold crossing delay); blue for amplitude-latency trading (*i.e*., threshold crossing delay augmented to 25 μs longer latency per dB attenuation). Inset in B shows magnified view of one null in the 100-μs echo. The 6ms range delay is marked by the leading (left) vertical edge of the row of dots for each echo. Without amplitude-latency trading, the nulls appear as interruptions, or voids, in the orange spectrograms, with a slight rightward trend as frequency approaches the center of the null and the lower amplitude moves the threshold crossings to slightly longer times. The deviations from the range delay can be used to estimate each null’s frequency entirely using threshold-crossing timing information.

The model’s code was written in MATLAB (*ver. R2019a*) and is publicly accessible https://github.com/gomingchen/SCAT. We built four modules to process the big brown bat’s 2-harmonic FM chirp-echo sequences and the bottlenose dolphin’s click-echo sequences (see modifications below). The third and fourth modules are included to demonstrate clutter rejection and other capabilities. While those modules share many features, such as bandpass filtering and thresholding, they each call customized functions and use distinct parameters to serve their own unique signal sequences. Running each module can take up to several minutes in a standard laptop. The names of main programs of the modules are listed in Table 1.

**Table 1.** Main programs of each module

| Number | Name of the main program of each module | Module function |
| --- | --- | --- |
| 1 | bat2HFM.m | The module designed to process 2 harmonic FM broadcast-echoes sequences. |
| 2 | dolClick.m | The module used to process bottlenose dolphin click-echo sequences. |
| 3 | clutterRejection.m | These two programs can demonstrate how amplitude-latency trading rejects clutter off to the side from the beam aim. clutterRejection.m uses a 1 harmonic FM sweep, and clutterRejection_dolClick.m compares echoes from off-axis targets to those from targets in line with beam axis. |
| 4 | clutterRejection_dolClick.m |  |

The example signals for each module are in a folder named “exampleSignals” and are loaded into MATLAB at the beginning of the main programs. The parameters describing the properties of the corresponding sequence (see Table 2) are written in the main programs as well. SCAT combs through each frequency channel to find the right crossing between a threshold and every echo and broadcast. The interval, i.e., the signal width, needs to be identified prior to the process as a known parameter along with a few other parameters shown in Table 2. Each module has an example signal and a set of parameters paired to describe the signal. Most parameters are self-explanatory, and their values reflect computational convenience more than biological influence, but three of them deserve mention: The amplitude thresholds start at 3% of full-scale broadcast strength and proceed in ten equal steps to 98% full scale. In the current version, echoes can be set at any amplitude in this range, but our testing of the model typically involves echoes no more than half of the broadcast amplitude, in keeping with known properties of the big brown bat’s gain control mechanism [28].

**Table 2.**
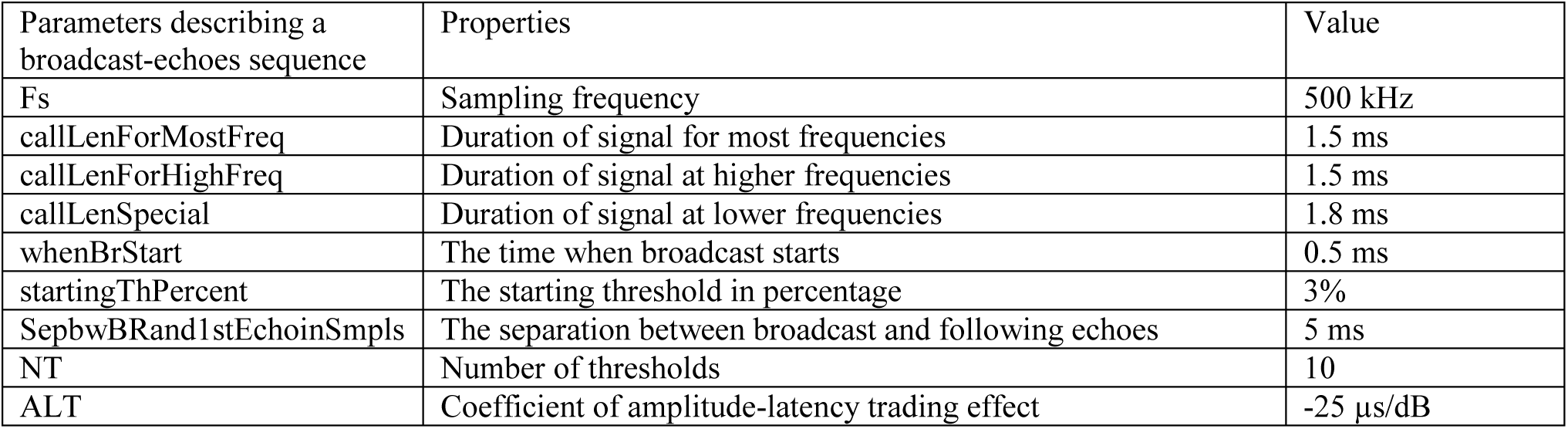
Parameters describing the properties of the example signal of 3^rd^ module

| Parameters describing a broadcast-echoes sequence | Properties | Value |
| --- | --- | --- |
| Fs | Sampling frequency | 500 kHz |
| callLenForMostFreq | Duration of signal for most frequencies | 1.5 ms |
| callLenForHighFreq | Duration of signal at higher frequencies | 1.5 ms |
| callLenSpecial | Duration of signal at lower frequencies | 1.8 ms |
| whenBrStart | The time when broadcast starts | 0.5 ms |
| startingThPercent | The starting threshold in percentage | 3% |
| SepbwBRand1stEchoInSmpls | The separation between broadcast and following echoes | 5 ms |
| NT | Number of thresholds | 10 |
| ALT | Coefficient of amplitude-latency trading effect | -25 μs/dB |

## Results from the SCAT Model

### Echo delay and glint estimates for FM chirps and clicks

To illustrate the output of the SCAT model, the information shown by the diagrams in Figs 3-6 is converted into a single, composite picture, the dechirped broadcast at its absolute spectrogram delay and the 100-μs 2-glint echo at its estimated glint spacing in Fig 7. Here, we use the term “image”, which is appropriate for describing the model’s output in this composite format because it is literally just a picture that shows the estimates of range delay (t) and glint delay (Δt). The arrangement of the X, Y, and Z axes is interlocking; it illustrates the magnifying effect of the spectrogram transformation from spectral ripples into the glint delay estimate alongside the range delay estimate derived from spectrogram correlation. Application of the term “image” to bat or dolphin biosonar perception rather than to display of SCAT outputs is a deeper matter to be discussed below.

**Fig 7.**
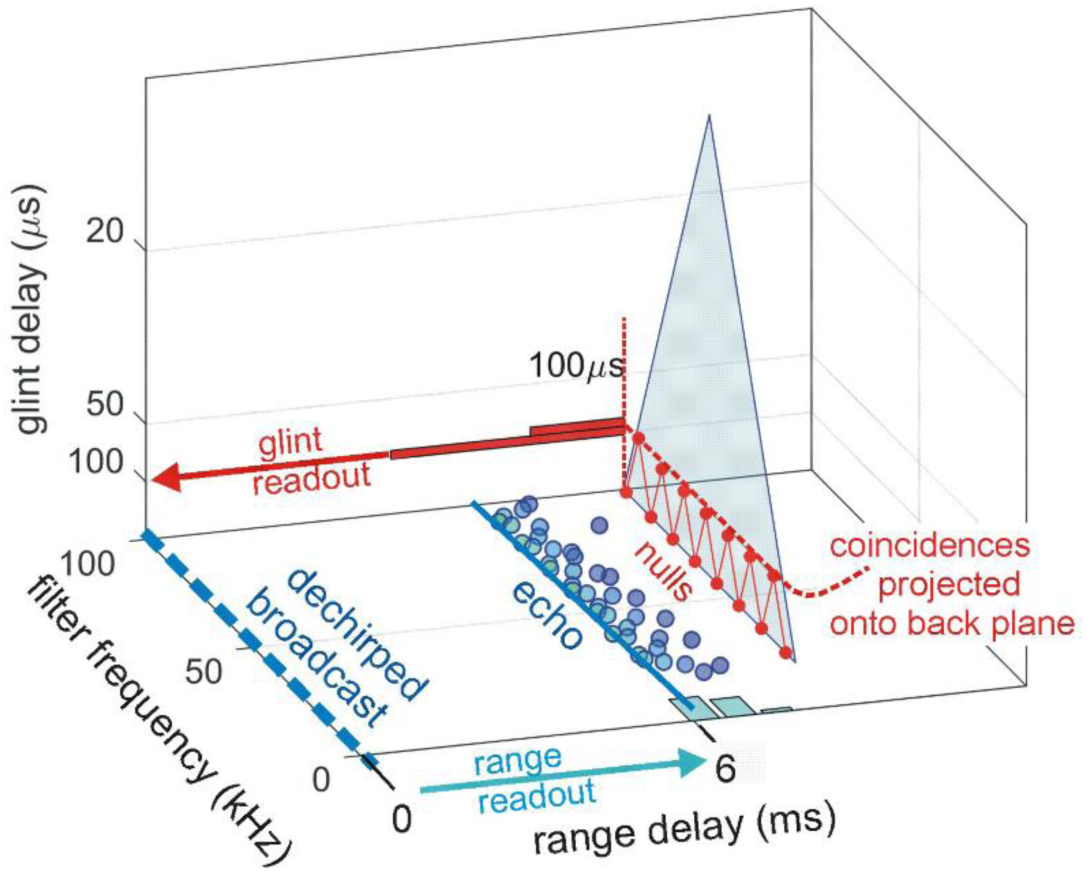
SCAT display format for 100 μs glint. Format of SCAT model output display for the twotwo-glint 100-μs test echo in Fig. 4. The horizontal (X axis) shows the range delay from Fig. 4B. The frequency scale (Y axis) shows center frequencies of the bandpass filters. The zero origin of the horizontal echo delay scale (X axis) is the dechirped FM pulse (dashed blue line). The echo’s dechirped spectrogram is traced on the X-Y plane by blue dots representing 2nd threshold level as an example. The threshold-crossings follow the scalloped pattern representing low amplitude values at the nulls with longer response latencies (amplitude-latency trading) (see Fig. 6). Range delay of the dechirped echo is marked by a light-blue histogram on the X axis. The leading edge of the histogram is taken as the delay because the tail of the histogram is lengthened by amplitude-latency-trading, which retards the delay estimate past the objective delay. The vertical scale (Z axis) shows the estimated time separation of the glints derived from the zig-zag coincidences plotted on the triangular null-detecting network (transparent blue). It is scaled nominally from 300 μs at the bottom (a limit set by the 300-μs filter integration-time where the spectrograms separate into two ridges; Fig. 1) to 12.5 μs at the top (a limit set by the maximum width of glint spacing capable of being registered on the 80 kHz wide base of the triangle). Spectral nulls are marked at 10 kHz intervals along the bottom Y-axis edge of the triangle. The back (X-Z) plane shows the glint separation extracted from the triangular null-spacing network from the frequencies of nulls and coincidences that trace a zig-zag pattern according to null separations, with the apex of the zig-zag triangles marking the glint spacing in microseconds. The glint spacing is displayed as a horizontal red histogram rotated sideways on the back plane. The height of the histogram is enlarged for better display. Numerical values of time on the horizontal (X) axis for target range and glint spacing on the vertical (Z) axis are combined into the bat’s perception of each target (Fig. 4D).

#### Display of SCAT output

Fig 7 shows the SCAT model’s complete output for the 100-μs 2-glint echo illustrated in Fig 4. This 3D format preserves the separation of the computational pathways that leads to the two delay estimates—for range delay and glint spacing. The X-Y plane in Fig 7 corresponds to the spectrogram plane in Fig 4B. The dechirped echo spectrogram is plotted on the X-Y plane by blue dots representing the same threshold level as for the broadcast. To show the target’s overall distance, the range delay compounded across different frequencies is marked by a light blue histogram aligned on the X axis. Note that this histogram is skewed, with an early sharp peak at the left and a longer, lower tail off to the right. The skew is caused by the longer latencies of threshold events at the frequencies of the nulls due to amplitude-latency trading (see Fig 6). The model determines the range delay from the leftmost rising edge of the histogram, which is derived from the short latency threshold-crossing at the peaks or left-hand segments of the scalloped curves tracing the dechirped echo spectrogram (Fig 4B). While the longer latency threshold marks are crucial for finding the frequencies of the nulls from their deviation to the right, they unacceptably bias the range-delay estimate overall and are discounted. The depths of the spectral nulls are often more than 50% less than the relative amplitude of the broadcast at the same frequencies. Consequently, the thresholds higher than half of the broadcast maximum amplitude won’t be crossed by the echo at the frequencies of the spectral notches. When a threshold fails to be crossed by the echo waveform at a certain frequency, the resulting voids will be recorded as a not-a-number (NaN) for that frequency in MATLAB. This equates to the responses of neurons in the big brown bat’s inferior colliculus when the stimulus is an FM sweep containing a spectral null at the neuron’s tuned frequency [106]. The swath of NaNs increases in width as the threshold becomes higher because more frequency channels around the notch would fail to contain crossings of the threshold, as shown in Fig 6. The MATLAB programs look for clusters of NaN from the crossings over all thresholds. The spread of later latencies is used to verify the locations of notches by comparing the latency of spectral notch vs. those of other frequency channels remote from the notch. After the null frequencies are entered on the frequency axis at the bottom of the triangular (transparent blue) network in Fig 7, the glint separations are displayed by the apex points on the zig-zag line in the triangular plane. The row of these apex points (red) on the triangular plane is projected onto the back (X-Z) plane, which has a vertical (Z) axis scaled in glint delay times in microseconds (the reciprocal of the frequency spacing of the nulls in kHz). This vertical axis is scaled from near 300 μs at the bottom (a limit set by the 300-μs filter integration time where the spectrograms separate into two ridges; see Fig 1) to 12.5 μs at the top (a limit set by the maximum width of glint spacings capable of being registered on the 80 kHz wide base of the triangular null-plotting network). This 3D display graphically depicts the target’s shape by its glint spacing as a horizontal red histogram on the back plane, using the vertical (Z) glint-delay scale for its bins. The target’s range, which is displayed by the numerical values of time on the horizontal (X) axis, is associated with its corresponding glint spacing on the vertical (Z) axis, now combined into a representation of both components that emerge in the bat’s perception of the target. As introduced in Fig 4C, the inversion of the SCAT representation from the peaks at different thresholds to the nulls located between the peaks reduces the density of the echo’s eventual perceived representation to the range delay (6 ms) and the glint delay (100 μs). The intervening spectrogram delays and the frequencies of the nulls are already a much sparser representation than the dense, multidimensional representation consisting of all the threshold events at each frequency and their individual spectrogram delays.

#### Display for three representative FM echoes

Using the 3D output display format described in Fig 7, Fig 8 illustrates the SCAT outputs for three 2-glint echoes that have glint separations of 50, 100, and 200 μs. They are arranged in a series at nominal range delays of 5, 12, and 18 ms, respectively. These delays are just used to package multiple echoes into the same signal stream to facilitate batch processing; other than being large enough separations to keep the processing epochs apart, we attach no particular significance to them in this and subsequent examples. Their glint-delay spacings correspond to target profiles of 0.85, 1.7, and 3.4 cm, respectively, in the middle for the sizes of insects hunted by big brown bats. Each echo’s dechirped spectrogram is traced on the X-Y plane by blue dots representing #2 threshold level, which follow the scalloped pattern representing low amplitude values at the nulls with longer response latencies due to amplitude-latency trading. Each dechirped delay is marked by a (black) histogram, which has the characteristic skew of threshold times being dragged to longer times by amplitude-latency trading. The time scale of these histograms is shown by the delay spacing of 2 ms between the successive dechirped delays. The glint separations from the nulls are displayed on the spectral to temporal networks (transparent blue triangles). The back (X-Z) plane shows the estimated glint separation projected from the triangular null-spacing network using the frequencies of nulls and coincidences that trace a zig-zag pattern according to null separations, with the apex of the zig-zag triangles marking the glint spacing in microseconds. As before, the vertical (Z) axis is scaled from 300 μs at the bottom to 12.5 μs at the top. The target shapes are displayed by the glint spacing as horizontal blue histograms and red number labels on the back plane.

**Fig 8.**
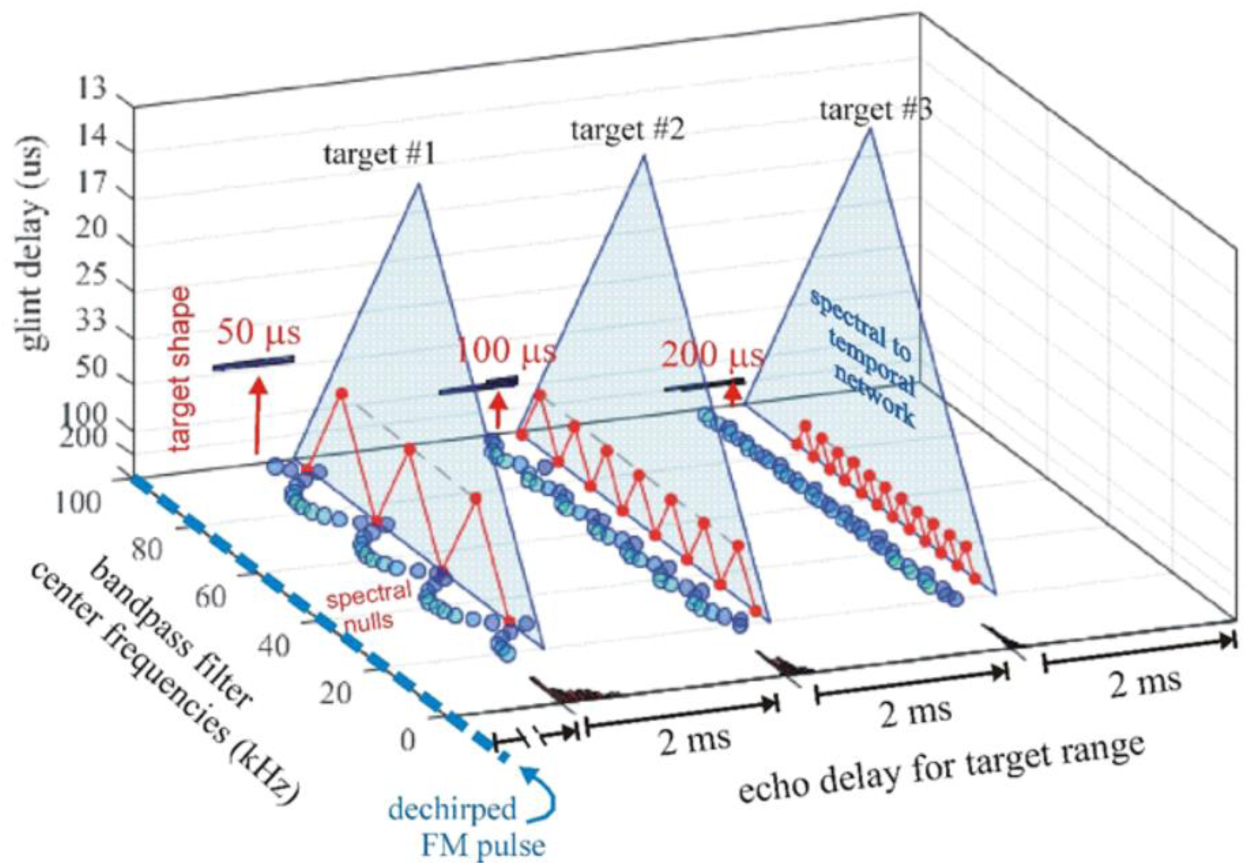
SCAT displays for different targets. The SCAT model outputs for three different two-glint echoes at different range delays are consolidated into a single 3D display using the format of Fig. 7. (No significance is attached to their range delays; the grouping is merely a way to set up multiple echoes for processing as a batch.) They have glint separations of 50, 100, and 200 μs, respectively (left to right). The frequency scale (Y axis) shows the center frequencies of the bandpass filters. The zero origin of the horizontal echo delay scale (Y axis) is the dechirped FM pulse (dashed blue line). Each echo’s dechirped spectrogram is traced on the X-Y plane by blue dots representing 2nd threshold level, which follow the scalloped pattern representing low amplitude values at the nulls with longer response latencies.). Each dechirped range delay is marked by a black histogram, with delay spaces of 2 ms along the X axis between the dechirped delays. The glint separations are displayed on the spectral to temporal networks (transparent blue triangles). The back (X-Z) plane shows the glint separation extracted from the triangular null-spacing network from the frequencies of nulls and coincidences that trace a zig-zag pattern according to null separations, with the apex of the zig-zag triangles marking the glint spacing in microseconds. The vertical (Z) axis is scaled from 300 μs at the bottom (a limit set by the 300-μs filter integration-time where the spectrograms separate into two ridges; Fig. 1) to 12.5 μs at the top (a limit set by the maximum width of glint spacings capable of being registered on the 80 kHz wide base of the triangle). The target shapes are displayed by the glint spacing as horizontal black histograms on the back plane. The height of each histogram is scaled for better display. Numerical values of time on the horizontal (X) axis for target range and glint spacing on the vertical (Z) axis are combined into the bat’s perception of each target.

#### Composite SCAT display for 2-glint echoes

To examine the performance of the SCAT model for different types of biosonar signals, we used a suite of 2-glint echoes derived from a dolphin click and a bat FM chirp. Differences in the model’s parameters reflect primarily the different frequency bands of bottlenose dolphins and big brown bats. There is one difference, however, that addresses how estimates of range delay are combined across frequencies and how glint delays are integrated into the range-delay images. Fig 1C has shown how the spectral nulls in 2- glint dolphin echoes are discernable as regularly-spaced ripples in the spectrogram for glint spacings of 35 and even 26 μs. The continuous vertical ridge of the click’s spectrogram enhances the appearance of the nulls as part of a ripple pattern. In contrast, as shown in Fig 1E, the dispersal of frequencies along the FM sweeps in the bat chirps, and even more the split of the spectrum into FM1 and FM2, obscures the regularity of the ripples until the glint delay increases to 100 μs, when two or more nulls fit into FM1 by itself. This difference accords with what has been established experimentally—that big brown bats treat the frequencies in FM1 and FM2 differently [33,43,65], and that the continuity of the spectrograms are correlated with a wide span of spectral ripple detection in dolphins [94–96] and a narrower span in bats [85].

#### Dolphin clicks

To model images for echoes of dolphin clicks, the SCAT filterbank is altered to use 121 parallel gammatone filters, center frequencies from 30 to 150 kHz, ERB 4 kHz. In this case, there is no explicit FM sweep to remove. This calibrating step removes differences across frequencies in the broadcast spectrum that lead to amplitude-latency trading in the timing of the threshold events triggered by the broadcast (see below). These perturb the SCAT model’s auditory spectrogram of the broadcast unless they are explicitly included when the zero time origin of delays are established independently at individual frequencies. In effect, the dechirping process “flattens” or “whitens” the transmitted spectrum, removing the small timing perturbations that amplitude-latency trading brings into the representation of the transmitted pulse. The SCAT model’s reliance on threshold-timing marks for all computations requires this precaution even for dolphin clicks.

To test the model with dolphin-like echoes, we used the same 2-glint delay separations illustrated in Fig 1A,B. These are 9, 18, 26, 35, 70, 100, 200, 300, 500, and 700 μs. This range of glint separations brackets the 250 μs integration-time for echo reception [3], which delineates spectro-temporal from purely temporal registration of the second glint. Conventional spectrograms for the echo sequence derived from the sample dolphin click are replotted in Fig 9A for comparison with the threshold-crossing spectrograms derived from the SCAT model’s bandpass filterbank in Fig 9B. Echoes from 26 to 200 μs contain multiple spectral nulls which appear as ripples along the vertical frequency axis, while for echoes at shorter separations of 9 and 18 μs the nulls are too wide and far apart to be characterized easily as ripples. Instead, they remove wide swaths from the spectrograms. Echoes with long separations from 300 to 700 μs pull apart to form two discrete click echoes. They have no spectral nulls because the glint spacing is longer than the 250-µs integration-time used to form the spectrograms. Fig 9B shows the stacks of threshold-crossing times for the outputs of the bandpass filters. The transmitted click (left) is represented by a smooth, continuous vertical stripe that follows each of the ten threshold levels (color bar in Fig 9B). For the echoes, with different placements of spectral nulls and peaks, the threshold crossings follow the scalloped pattern illustrated in Fig 5. The peaks are marked by the higher thresholds (orange, red) shifting to the right, while the nulls appear as voids between the peaks for echoes from 9 to 200 μs, where only the lowest thresholds (blues) are detected. Note that the nulls and peaks are discernable in the 100 μs echo but not the 200 μs echo; this is only because the dots used to mark the events are too large. In the 300, 500, and 700 μs echoes, each separate glint reflection is separated but not visible because the second in each pair is obscured by the spread of the dots that mark the threshold events for the first reflection in the pair.

**Fig 9.**
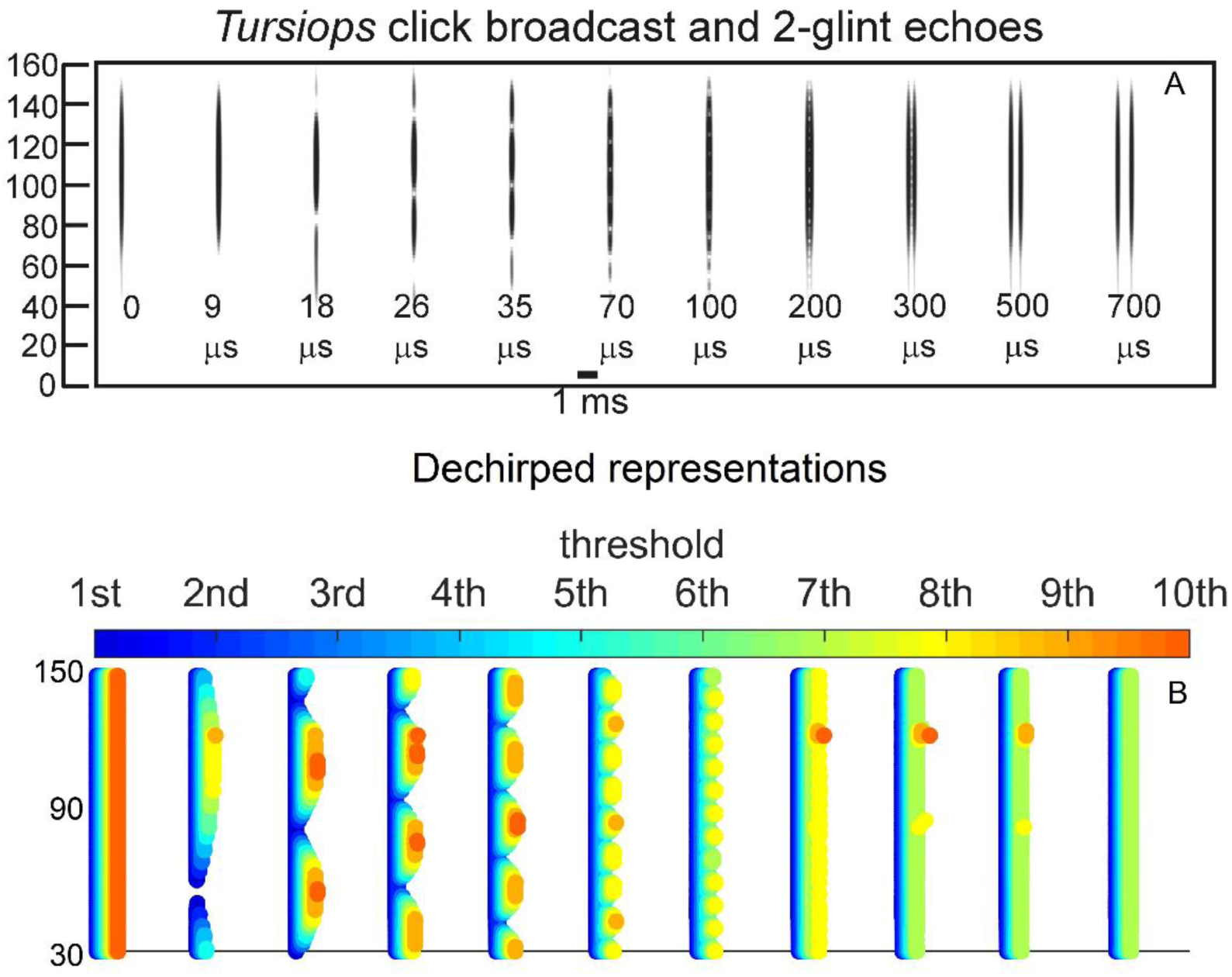
Conventional and SCAT spectrograms for two-glint click echoes. (**A**) The bottlenose dolphin click is formed into two-glint echoes (glint delays 0 to 70 μs) that appear to have separate highlights in the time waveforms (Fig. 1A) but merged clicks with spectral nulls in the spectrograms for glint delays of 9 to 200 μs (replotted from Fig. 1C). (**B**) After conversion into auditory spectrograms by the bandpass filterbank, the timing of the ten threshold-level detections (#1 to #10 in color bar) are nearly superimposed in the dechirped broadcast spectrogram (“0 glint,” left), sliding 0.12 ms slightly rightwards as the threshold level rises from lowest (blue) to highest (red). The glint interference patterns modulate the amplitude of the echo spectrograms so that the full range of threshold-crossings (from blue to red) appears at the peaks but not at the intervening nulls, where only the lowest thresholds yield any detections at all. Due to amplitude-latency trading, the time-axis re-represents the amplitudes of the peaks and nulls along the frequency axis by the scalloped pattern of threshold levels. Some echoes don’t have higher threshold crossings because the gain control in SCAT model is an approximate process to align the echo strength to be as close to the broadcast but lower.

To facilitate running batches of echoes in the SCAT model, the time-window parameters found in dolClick.m in https://github.com/gomingchen/SCAT have to be adjusted to allow detection of separate glints as events in time at short spacings such as 300-700 μs. Specifically, SCAT can batch-process a series of broadcasts followed by multiple click echoes (Fig 9A) with a parameter that describes the intervals between those echoes. The parameter is called “callLenForMostFreq” and tells the model when to look for the crossing from next echo. However, echoes with glint spacing 300, 500, and 700 µs consist of two discrete click echoes, and the goal is to recover both of them. The interval between the two closely spaced glint echoes is very short. When clipping the original broadcast and each of those three echoes, a shorter interval thus needs to be used to capture the crossing of the second glint echo. This capability is presented here just as a convenience for batch-processing of several echoes at a time. For realistic sonar scenes, multiple echoes often follow each broadcast, and the model normally is configured for this contingency.

Fig 10 shows the SCAT 3D display for the click echoes in Fig 9 using the format explained in Figs 6 and 7. The dechirped spectrograms are traced on the X-Y plane by threshold-crossing marks (blue dots, shown here just for threshold #3 so the complete vertical spectrogram stripe is visible without being obscured by overlay of dots that mark higher thresholds being crossed, as in Fig 9B. Range delays (t) are displayed on the horizontal X axis as a light-blue histogram for each echo. (The delay axis is segmented by breaks to magnify individual histograms, with each range delay assigned to the left-hand principal peak of the histogram). The broad, continuous spectrum of the click shows the interruption of the threshold crossings at the nulls, which are transferred from the X-Y plane onto the triangular networks for extracting the frequency spacing of the nulls in each echo (see Figs 4C, 6). The null spacing is determined from the apex points of the red zig-zag lines that connect the null frequencies, which are projected onto the back X-Z plane to register the approximate spacing of the nulls in terms of their inverse, the time separation of the glint reflections. The continuous nature of the click spectrum makes it easy to register all of the glint separations from the 18-μs echo to the 200 μs echo. However, there is only one null in the 9-μs echo, at 83 kHz, and no second null inside the band of the click. Consequently, the 9-μs echo does not fit two nulls into the width of the triangular network’s base and the network cannot find a null spacing for that echo. Beginning at the 18-μs glint spacing, the network finds adjacent pairs of nulls and successfully estimates the glint delays (Δt) from the frequency spacing of the nulls (Δf). For the 300, 500, and 700 μs echoes, the two glint reflections pull apart to be registered as separate range delays (blue histograms on X axis), with no null-based glint-delay estimates because the nulls are too close together to be detected as separate in frequency. The combination of glint delay and range delay estimates effectively characterizes the 2-glint echoes on both sides of the integration time window that defines whether the second glint evokes spectral nulls or a separate range delay.

**Fig 10.**
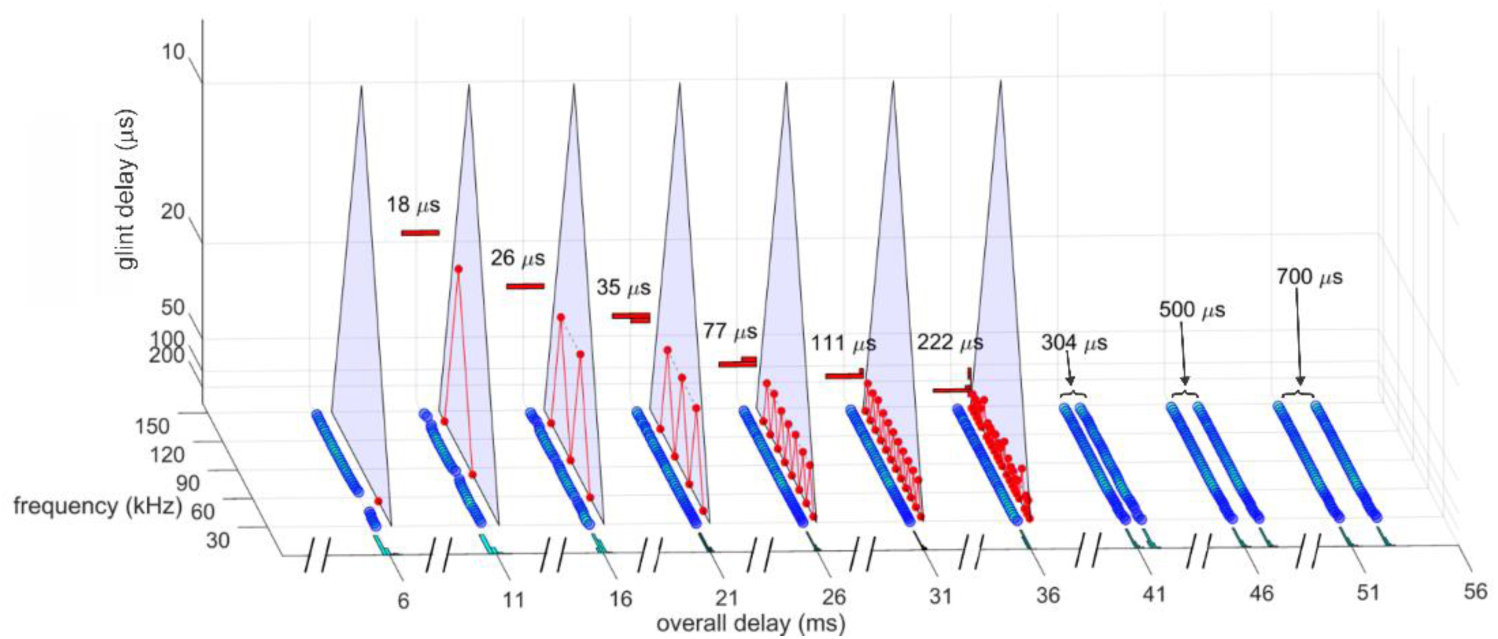
SCAT displays for two-glint click echoes. The 3D SCAT plots of the whole series of two-glint click echoes from Fig 9 portray the range delay of each target on the horizontal X axis and the glint delay of each target on the X-Z plane. Using the display format defined in Fig 7, the dechirped spectrograms are plotted on the X-Y plane. For glint delays from 18 to 200 μs in Fig 9, the triangular network extracts the glint delay for each target from the apex points of the red zig-zag line on the triangular network. Combining the whole series of echoes into a batch for processing yields a graph of glint spacing on the X-Z plane. When the glint spacing reaches 300 μs, the two glint reflections pull apart in the spectrograms and no longer have an interference spectrum to be plotted on the triangular network or projected onto the back plane. Both glints then appear entirely on the horizontal overall delay axis.

#### Bat chirps

Fig 11A shows conventional spectrograms for the same family of 2-glint echoes illustrated for dolphin clicks but instead derived from a 2-harmonic FM bat chirp as broadcast. Fig 11B traces threshold-crossing times for the FM sweeps in the same echoes as in Fig 10A with the frequency axis broken into two bands, one for FM1 and the other for FM2. The dots are for threshold #3, again, chosen to show the separation of the sweeps with no threshold dots hidden behind others due to different times for other thresholds. Fig 11C,D shows the corresponding dechirped SCAT spectrograms, now completely separated into two frequency bands for FM1 and FM2. Both the conventional spectrograms in Fig 11A and the complete SCAT spectrograms in Fig 12 illustrate the problem posed by the two harmonics: Spectral peaks are identified by the scalloped pattern of the threshold crossings, which are marked by the spread of the dots to the right, while the nulls are identified with spaces between the peaks, where only the lowest thresholds are crossed. Unlike for the click echoes in Fig 9, however, where the succession of peaks and nulls is easily seen, the split between the two parts of the spectrum for FM1 and FM2 obscures the orderly appearance of the nulls until the glint spacing is large enough that two or more nulls are present just in one of the harmonics. At glint spacings from 9 to 35 μs, the break between the two harmonics adds a discontinuity to the dechirped spectrograms. Only at 70 μs does the rippled appearance of the nulls begin to appear across the harmonics, and at 100 μs it is unmistakable. The ripples then continue to be visible for spacing of 200 and 300 μs, but at 500 and 700 μs, there are no ripples because the reflections have pulsed apart to appear as separate spectrogram ridges.

**Fig 11.**
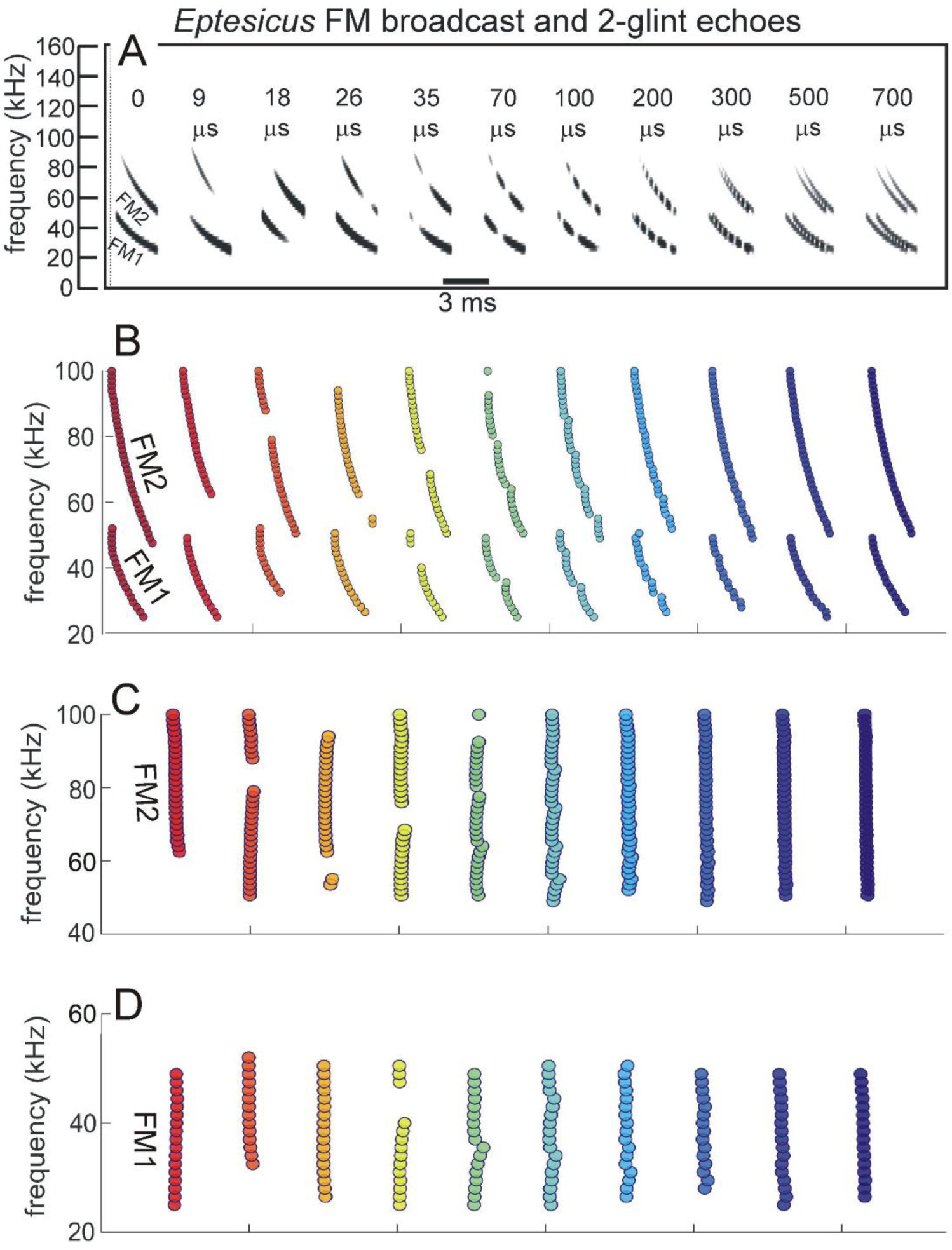
SCAT spectrograms for two-glint FM echoes. (**A**) Conventional spectrograms for the series of two-glint echoes (0 to 700 μs glint delays) of a bat chirp replotted from Fig. 1E. The broadcast has two harmonics (FM1, FM2) (left). For glint delays of 9-300 μs, the spectrograms of the glint reflections are merged and have spectral nulls that appear as patterned ripples for 70-300 μs. At short glint delays of 9-35 μs, the nulls are too far apart to appear as ripple; instead, they obliterate wider segments of the spectrograms. At longer delays of 500-700 μs, the reflections pull apart to form separate ridges that trace the two harmonics. (**B**) SCAT spectrograms showing threshold crossings as points marking the instantaneous frequency of the sweeps by their threshold crossings (to avoid overlap of dots representing different thresholds, only threshold #3 is illustrated here). The nulls are represented by voids and the scalloped shape of the curves tracing the FM sweeps. (**C,D**) Dechirped SCAT spectrograms with FM1 and FM2 plotted separately. Segregation of harmonics into two bands is done to take into account the way the bat separates them and assigns FM1 as necessary for delay processing, with FM2 only used if its frequencies mirror those in FM1 [64].

**Fig 12.**
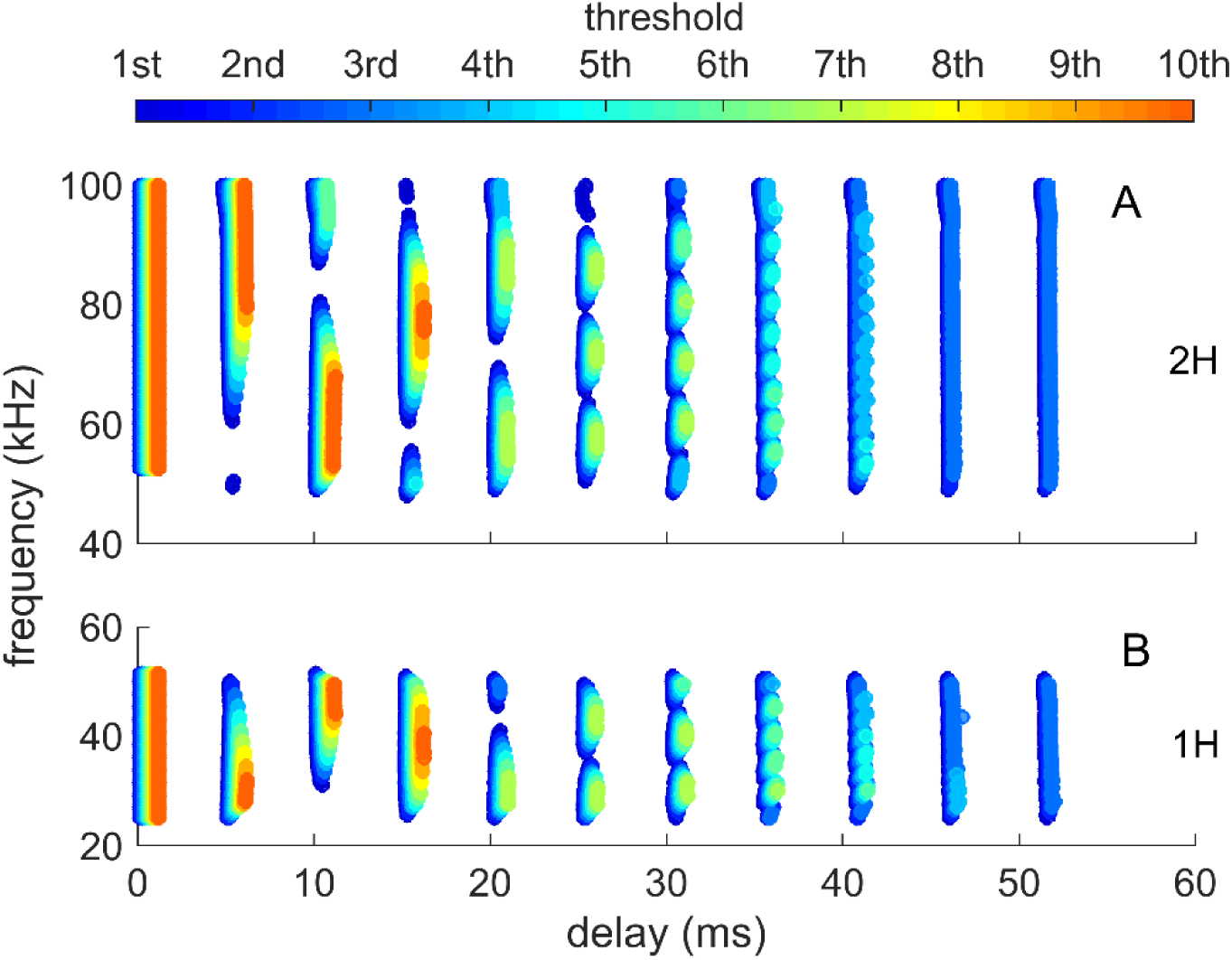
Dechirped SCAT spectrograms for two-glint FM echoes. Dechirped spectrograms of FM echoes at all ten threshold levels (color bar in the top row) show the characteristic scalloped appearance due to the movement to longer latencies at higher thresholds (yellow to red) for frequencies of peaks and the restriction of threshold crossings to the lowest thresholds at the nulls (blue). FM1 and FM2 are plotted separately, with both harmonics in the broadcast at left. The threshold crossings for the broadcast follow vertical stripes, but the amplitude variations in the two-glint echoes break up the stripes into segments that appear as islands centered on the peaks (see also Fig 4C). The nulls are recognizable by the holes in the spectrograms.

Fig 13 illustrates the sequence of FM echoes with the same 3D display already employed to describe the SCAT model’s outputs. The range delays are shown by the blue histograms along the X axis, again with the scale having breaks that separate the 2 ms segments that magnify the range-delay histograms. Glint delays are extracted from 18 to 300 μs echoes, although with more variability than for the click echoes in Fig 10 due to the effect of the boundary between FM1 and FM2, which interrupts the spectrograms. This variability originates in the null-detecting coincidence points on the triangular network. The zig-zag lines are more stable for the 100, 200, and 300 μs echoes, and they do produce a well-defined glint-delay histogram for each echo on the X-Z back-plane. All of the echoes with 300-μs or less glint separation register as a single range-delay histogram on the X axis, while the 500 and 700 μs echoes separate to form two independent returns (rows of blue threshold dots) and are registered separately by their range delay histograms on the X axis. Despite the evident awkwardness of the harmonic separation in Figs 10-12, it is in keeping with the bat’s asymmetric treatment of FM1 and FM2. Instead of combining the two harmonic bands into a single, continuous spectrum from 25 to 100 kHz, the bat prioritizes frequencies in FM1 and only uses frequencies in FM2 if the corresponding FM1 frequencies are also carried in the echoes [33,43,64,65,85,121]. The bat’s emphasis on the lowest frequencies exploits the greater reflectivity of the surrounding scene at these frequencies; echoes arriving from off-axis or distant objects are always lowpass-filtered compared to echoes arriving from nearby and on-axis [11].

**Fig 13.**
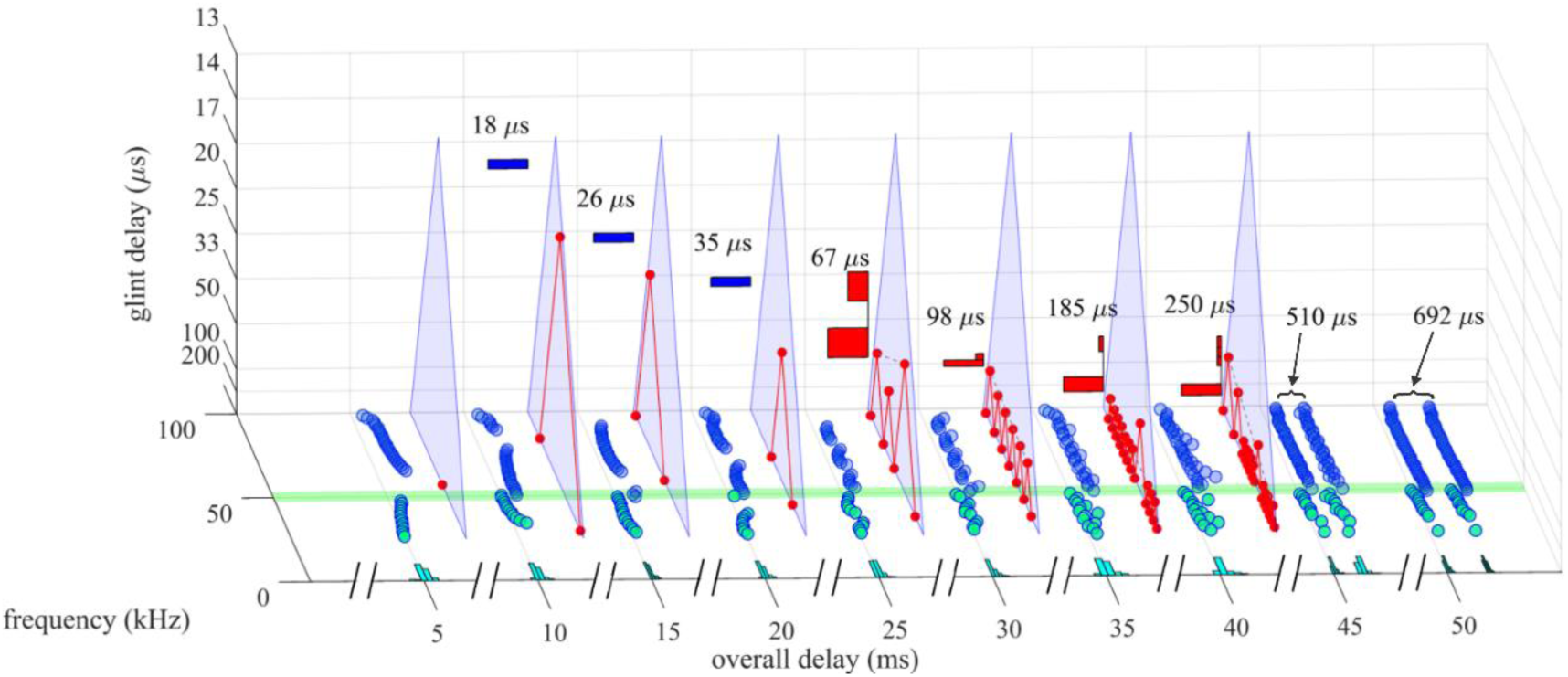
SCAT display for two-glint FM echoes. The 3D SCAT plot for the series of FM echoes in Fig. 12 portrays glint spacing of each target on the X-Z plane associated with its corresponding overall echo delay on the X-Y plane. The spectrograms are trace by blue and green dots for threshold crossings (#3 is illustrated). The dots (blue) tracing FM2 are moved (“dithered”) slightly to the right to avoid overlap with FM1 (green). Estimates for glint spacings from 18 to 300 μs are determined from the triangular null-spacing network and projected onto the back (X-Z plane). For echoes with recognizable spectral ripples (glint delays of 70, 100, 200, and 300 µs) that are traced by red zig-zag lines on the triangular network, the intersection of FM1 and FM2 introduces perturbations in the estimates. When the glint spacing exceeds 300 μs, the two glint reflections pull apart to form separate dechirped spectrograms and no longer have an interference spectrum to be plotted on the back plane. Both glints then appear entirely on the horizontal overall delay axis.

### Clutter Rejection

In most biosonar scenes, multiple objects are present, located in different directions and at different distances. Echoes returning from a target of interest (*e.g*., insect, fish) have to compete with echoes from the other objects, called clutter, to be located and perceived. As described above, the SCAT model uses inversion of the acoustic pattern of spectral ripples from frequency to delay to extract information about a target’s shape while determining range delay for localization. The mechanism for displaying shape is the triangular-shaped network that finds the frequency spacing of spectral nulls (Figs 4, 7). The SCAT image of a target is “in focus” when the acoustic factors affecting the auditory spectrogram are distilled down to just the effects of reflection from the object’s glints, with no spectral coloration—usually lowpass filtering— caused by off-axis or distant location. Then, the X-Z plane of the 3D SCAT display portrays only the object itself. If the object is off to the side of the emitted sound beam or far enough away, its echoes necessarily are weakened overall. Additionally, the spectrum of these echoes is affected by direction and distance; it will contain not only the local spectral ripple related to any object’s multiple glints but also a more global lowpass reduction in strength at all of the higher frequencies due to the frequency dependence of both broadcast beam width and atmospheric absorption. When the bat aims its transmitted beam on the target of interest, it ensures that all of the frequencies in the sound are projected onto that object. However, the consequence is that clutter located off to the sides is not ensonified by the full broadcast spectrum. While the lower frequencies impinge on the clutter as well as the target, the incident sound contains progressively weaker high frequencies as the off-side angle increases. Lowpass filtering thus is the dominant characteristic of clutter echoes, and the SCAT model treats it in a unique way inspired by experimental results from bats [11].

#### Lowpass dolphin clutter

Due to their continuous, broad spectrum, the dolphin clicks highlight how the SCAT model treats lowpass echoes as a by-product of the glint-delay mechanism. Fig 14A shows spectrograms for a transmitted dolphin click and for two simulated echoes; one is a 100-μs 2-glint echo (Fig 9) from a near target located on the broadcast beam, and the other from the same 2-glint object but mimicking a location off the beam’s axis, where it is ensonified with lower strength at higher frequencies. In Fig 14A, the regular pattern of spectral ripples at intervals of 10 kHz is easily seen in the 2–glint echo and these ripples are visible in the lowpass echo, but the added lowpass filtering truncates the upper end of the spectrogram so the ripples disappear into the larger reduction in amplitude. Nevertheless, in Fig 14B, the dechirped SCAT spectrogram, which shows all ten thresholds (color bar), resolves the spectral ripple across the entire band of frequencies. The dechirped spectrogram of the lowpass echo follows the same pattern in that the 10 kHz ripples are visible across the spectrum, but here the role of amplitude-latency trading becomes especially obvious. In the region of the lowpass effect, the threshold crossings are limited to only the lowest levels (blues), and the timing of the threshold crossings shifts to longer latencies, which curves the SCAT spectrogram to the right at higher frequencies. The dropout of many threshold events and the longer latencies makes the lowpass region resemble a single, broad null about 40 kHz in width, mixed with the narrower, glint-related nulls spaced 10 kHz apart.

**Fig 14.**
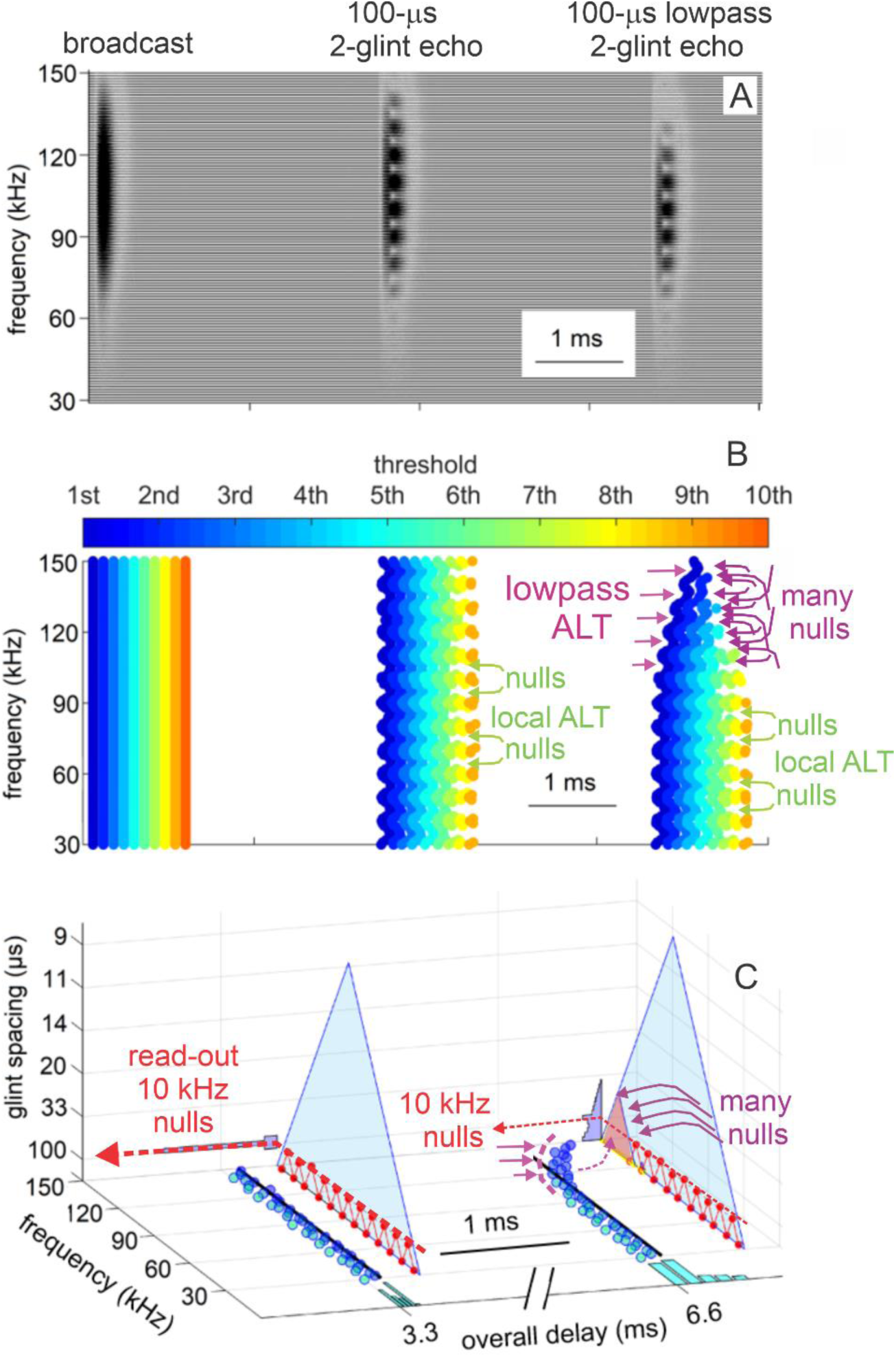
3D SCAT display for lowpass click echo. The continuous spectrum of the dolphin clicks helps to illustrate how the SCAT model processes clutter echoes. (**A**) Periodic spectral ripple for the 100-μs two-glint echo is visible at 10 kHz intervals in the spectrogram from the bandpass filters (see Fig. 9), while both the ripple and the broader reduction in strength across high frequencies are visible in the lowpass 100-μs echo. (**B**) The dechirped SCAT spectrograms show the same pattern of peaks and nulls across the echo spectrum for the two-glint echo (stack of ten thresholds in color bar), but the lowpass echo’s broadly lower amplitude at high frequencies reduces the threshold crossings to only the lowest (blue) and retards their latencies due to amplitude-latency trading. The lowpass roll-off in amplitude is translated into the rightward-curved trajectory of threshold crossings, which brings them into the same latency format used to detect nulls (longer latencies at some frequencies relative to others). (**C**) The triangular network of null-detecting coincidence nodes readily extracts the 10 kHz null spacing in the two-glint echo and displays it as a 100-μs glint delay on the X-Z plane. In the lowpass echo, the 10 kHz spacing of the nulls also is extracted and displayed, but the wide region of lower amplitudes and longer latencies between 100 and 140 kHz resembles an additional, very wide null. The triangular network registers it as a multiplicity of narrower nulls with a range of different widths from the minimum null spacing of about 3 kHz to about 15 kHz. The resulting nulls all are entered onto the triangular network, which converts them into numerous glint-delay estimates ranging in size from the inverse of 3 kHz (300 μs) to the inverse of 15 kHz (70 μs) (see Fig 15).

The presence of the single, broad null runs up against a limitation of the SCAT model as derived from the big brown bat. Neurons in the bat’s central auditory system are tuned to frequencies from 10 to 100 kHz, with frequency tuning widths from about 1-2 kHz to 15 kHz [93]. The 10 kHz ripples in the spectrogram are within the range of these tuning widths and can be registered as a series of discrete nulls at individual frequencies in the spectrogram. However, even the broadest tuning curves are too narrow to register a single spectral null as wide as the null implied by the region of lowpass filtering in Fig 14B. Consequently, the broad null created by lowpass filtering is not capable of being represented as such. Instead, it has to be approximated as a series of narrower nulls of different center frequencies and widths that fill up the 40 kHz wide gap.

Fig 14C shows how this approximation is done. The 3D SCAT display for the 100 μs 2- glint click echo traces the registration of the spectral nulls through the triangular network, which determines their frequency spacing and projects it onto the glint-delay image on the X-Z plane. With no other features of the echo spectrum, the read-out of the 10 kHz ripples is an unambiguous 100-μs glint spacing. Fig 15A (red curve) plots this glint-delay estimate by means of the histogram oriented sideways and projected onto backplane of Fig 14C. For the lowpass echo, the same process in Fig 14C finds the ripples for the two glints and projects the 100-μs glint-delay estimate onto the X-Z plane, too. However, this estimate is now accompanied by another, less sharp estimate derived from the set of narrow nulls used to fill in the nominal, wide null created by the lowpass filtering. Those nulls are found in SCAT specifically because their arrivals are later than the latest of the majority arrivals of all crossings. Collectively, these nulls are shown by a purple triangle superimposed on the high-frequency corner of the larger triangular network in Fig 14C. There are many of these nulls with multiple frequency spacings that extend from about 3-4 kHz to about 15 kHz in the vertical dimension of the triangle. The corresponding glint-delay estimates (Δt) tied to each of the pieces that fit into this highlighted triangle, which are the inverse of the null spacings (Δf), fill in the gap from over 300 μs down to about 60 μs. The distribution of these glint delays is shown in Fig 15A (blue curve). The peak in this curve corresponding to the 100-μs glint separation is visible, but it is obscured because it rides on top of the broader plateau of glint-delay estimates created by filling in the wide lowpass region with multiple narrower nulls. Suppose, however, that this display of the glints is treated as a unitary image with an overall image “intensity” distributed across all of the glint delays the image contains. Then, the pure glint image for the focused echo has its primary content only the 100 μs peak itself, with negligible further contributions from glint delays that occupy positions remote from the 100 μs point. The lowpass image contains not only the 100 μs peak, but also the “fill” contributed by all of the glint delay values from 50 to 400 μs, which contributes much more to the image’s total intensity than does the 100-μs peak. This blurs the image, defocusing the representation for any of the actual glints in the object that returns the lowpass echo, which is part of the clutter. In Fig 15, if red and blue curves are normalized by their image “energy” – the area under the curve, the obscuring effect of the lowpass “fill” can be mitigated (Fig 15B). The fill spreads the content of the clutter’s image over such a wide span that it has relatively little content at any one putative glint delay within this span. The much more condensed content of the image for the target that is in focus (the 100 μs glint spacing) protrudes above the scaled-down content of the image of the clutter at the same glint delay. The overall effect is to widen the glint image of the clutter over much of the range of glint delays that the triangular network is capable of decoding. Glint delays for clutter that are widely spread across the triangular network contribute to the image at numerous locations away from the 100-μs glint, and normalization spreads their influence wide enough to improve the contrast surrounding the concentrated 100 μs peak.

**Fig 15.**
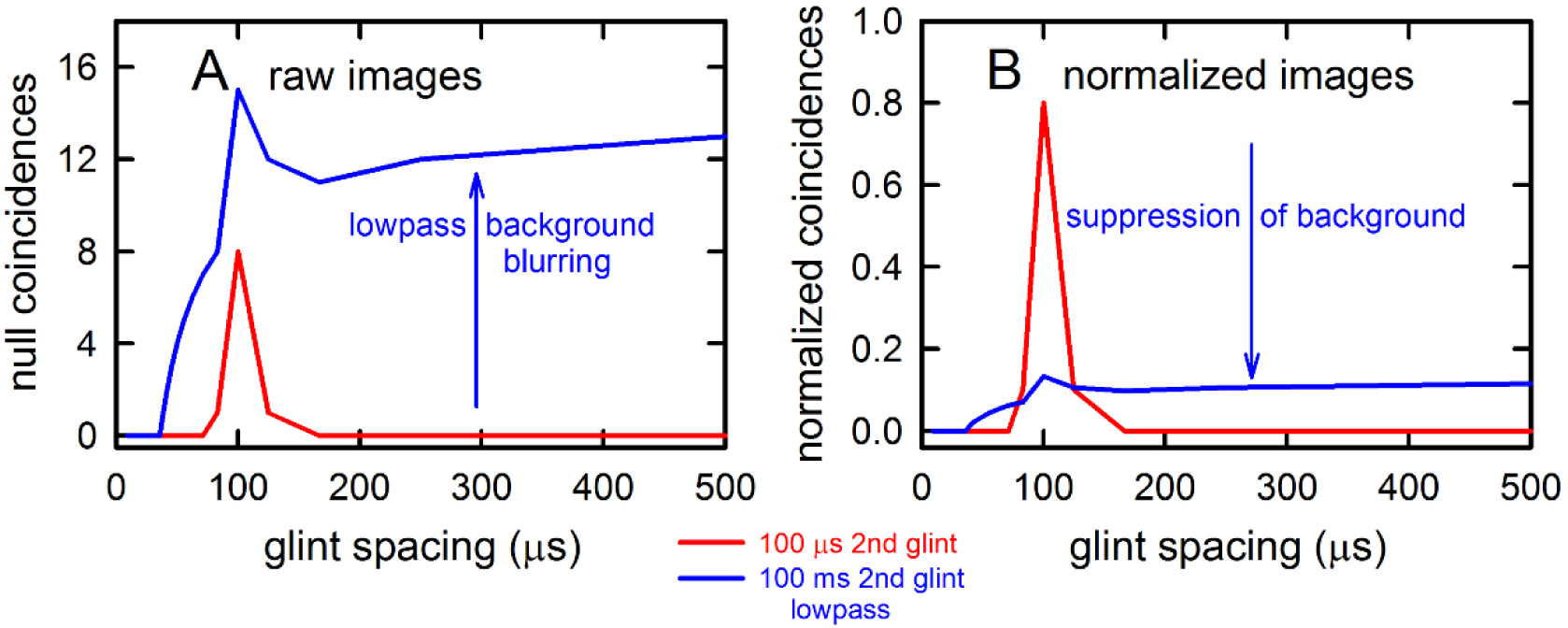
Glint-delay display for click echoes. (**A**) Graph showing the glint-delay distributions displayed in Fig. 14C on the X-Z plane and derived by the triangular network for the 100-μs two-glint echo (red) and the lowpass 100-μs two-glint echo (blue). The 100-μs peak is the salient feature of the two-glint echo from its 10 kHz null spacing, while the lowpass echo shows the 100-μs peak largely swamped by the broad set of nulls from the lowpass region of the triangular network. The lowpass echo represents off-axis clutter, and the blurring effect of the many extraneous nulls detected across the lowpass region illustrates the masking effect of clutter interference with perception of the second glint. (**B**) The blurring effect that obscures the 100-μs glint in A is mitigated by using the total image “energy,” or the area under each curve, to normalize the display of glint delay, which becomes unmasked by the spread of energy across the glint-delay axis.

#### Lowpass bat clutter

To explain how lowpass filtering effects the SCAT images derived from FM echoes, we created a sequence of signals that mimic the frequencies used by big brown bats but have only one harmonic sweep, from 100 kHz down to 20 kHz. Fig 16A shows the spectrograms created by the FM broadcast and a series of seven numbered echoes that have been subjected to increasing amounts of lowpass filtering. The 2-glint reference echo for this series has a 100 μs glint separation (echo #1), which creates the expected spectral ripple composed of nulls spaced at 10 kHz intervals. Different amounts of lowpass filtering are added to the basic 2- glint echo to illustrate what happens when the attenuation caused by the lowpass filter gradually overcomes the glint interference nulls at the higher frequencies. Initially, the filtering affects only the highest frequencies around 95-100 kHz, but gradually attenuation spreads downward to lower frequencies from one echo to the next (shaded red triangle in Fig 16A). At the strongest lowpass condition (echo #7), attenuation spreads down to 50-60 kHz. In effect, just as for the dolphin click echoes, the lowpass filtering is manifested as a broad null located at the upper end of the spectrum, a null that expands downward in frequency to become more and more difficult to register as a single null by itself.

**Fig 16.**
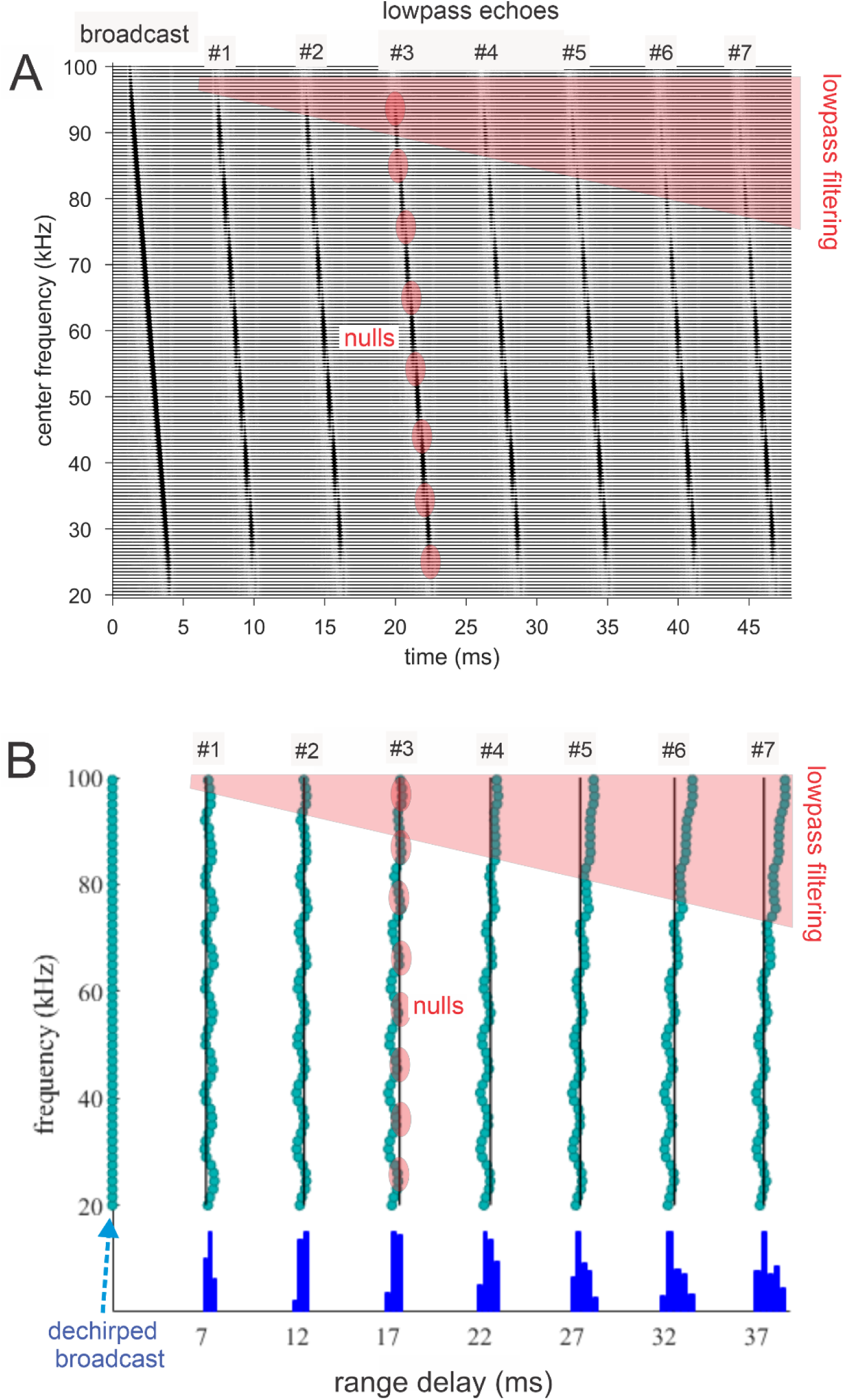
Effects of progressively increasing lowpass filtering on FM echoes. (**A**) Bandpass filter spectrograms of FM broadcast (for simplification of illustration, signals have just 1 harmonic sweep from 100 down to 20 kHz) and series of seven numbered 100-μs two-glint echoes that have increasing amounts of lowpass filtering (shaded triangle for filtering; shaded ovals for nulls). As lowpass filtering cutoff spreads downward in frequency, the nulls are obscured in the affected high-frequency band. (**B**) Dechirped SCAT spectrograms (just threshold #1 at 3% amplitude) for the broadcast (at zero on the range delay axis) and the seven numbered lowpass-filtered echoes from A. The spectral nulls (shaded ovals) are visible from the scalloped shape of the spectrograms due to the longer latencies at the individual nulls’ frequencies from amplitude-latency trading. The gradually increasing lowpass filtering (shaded triangle) causes the threshold crossings to shift to longer latencies over higher-frequency region from amplitude-latency trading. It transposes the lower amplitudes from lowpass filtering into longer latencies but also creates the appearance of a single, wide null over the 70-100 kHz high end of the spectrum. How this spurious null is registered turns out to be the key to coping with clutter.

The dechirped SCAT spectrograms in Fig 16B show the broadcast (at left) and the series of 100-μs echoes as they are registered by threshold crossings in the frequency channels of the model. The spread of lowpass filtering downward along the frequency axis is illustrated by the expanding shaded triangle. For clarity, these dechirped spectrograms are traced by only one of the model’s ten amplitude thresholds (threshold #1, for 3% amplitude; blue dots). They show the transformation of the glint nulls at 10 kHz intervals into the scalloped pattern of latencies, and beginning at the highest frequencies the added latency increases due to lowpass filtering from the highest frequencies in echoes #2 to #4 to more substantial latency shifts covering frequencies all the way down to 60 kHz in echoes #5 to #7. The range-delay distributions for each echo are plotted in Fig 16B along the X-axis as dark blue histograms. As was the case for the lowpass click echo (Fig 14B), the region of lower amplitudes from lowpass filtering is manifested as a single, broad null in the spectrum that is transposed into its corresponding latency increases due to amplitude-latency trading. The frequencies encompassed by this null spread downward from 100 kHz to around 60 kHz as lowpass filtering is made stronger. The fidelity of this transformation from numerical amplitudes to numerical latencies is particularly evident here.

Fig 17 shows the 3D SCAT display for the dechirped broadcast and the series of seven echoes with increasing amounts of lowpass filtering. Each echo has a 100-µs glint spacing that feeds into the blue triangular network to find the frequency spacing of the nulls. The red zig-zag lines on the networks are anchored at the base of each triangle on the frequencies of the nulls, where the scalloped pattern of the threshold crossings (again, just threshold #1 is illustrated) shifts along the curved trajectory of the lowpass-affected latencies but still are recognized by their local latency patterns. The apex of each converging segment on these lines marks the size of the null spacing (10 kHz), which is projected onto the X-Z plane to register the 100-μs glints (horizontal dashed red arrow). The glint nulls are registered for all of the echoes (see below), even though the encroaching lowpass filtering bends the SCAT spectrograms to the right (Fig 15B, so that) the glint nulls are successfully detected. The lowpass-filtering effect in the echoes combined with an addition of amplitude-latency trading to create a wide zone of latencies that resembles a single null covering all of the lowpass frequencies. As was the case for the lowpass click echo (Fig 14B,C), the limited frequency resolution of the triangular network from about 3 kHz at the short end to about 30-35 kHz at the long end means that the single wide null associated with lowpass filtering is not registered as one null. Instead, it is approximated by filling it in with numerous nulls having narrower widths that activate significant segments of the blue triangular networks (shown as purple triangles on the blue networks in Fig 17). These numerous, filled-in nulls are projected onto the X-Z plane as additional glint-delay values along with the 100-μs glint delay derived from the 10 kHz nulls. The latency shifts required to detect nulls are small. The minimal shifts associated with the 10 kHz nulls are extracted for all of the frequencies below 60-70 kHz, where the expanding lowpass filtering region does not prevent their being detected. The latency shifts produced by the lowpass filtering also are detected and used to generate the nulls that fill in the lowpass region of frequencies. Only a very small region of fill-in nulls is found for echo #1 (tiny shaded triangle on blue triangular network), but this region expands rapidly for echoes #2 and #3, and becomes saturated for echoes #4--#7. These fill-in nulls appear in the glint-delay histograms on the X-Z backplane.

**Fig 17.**
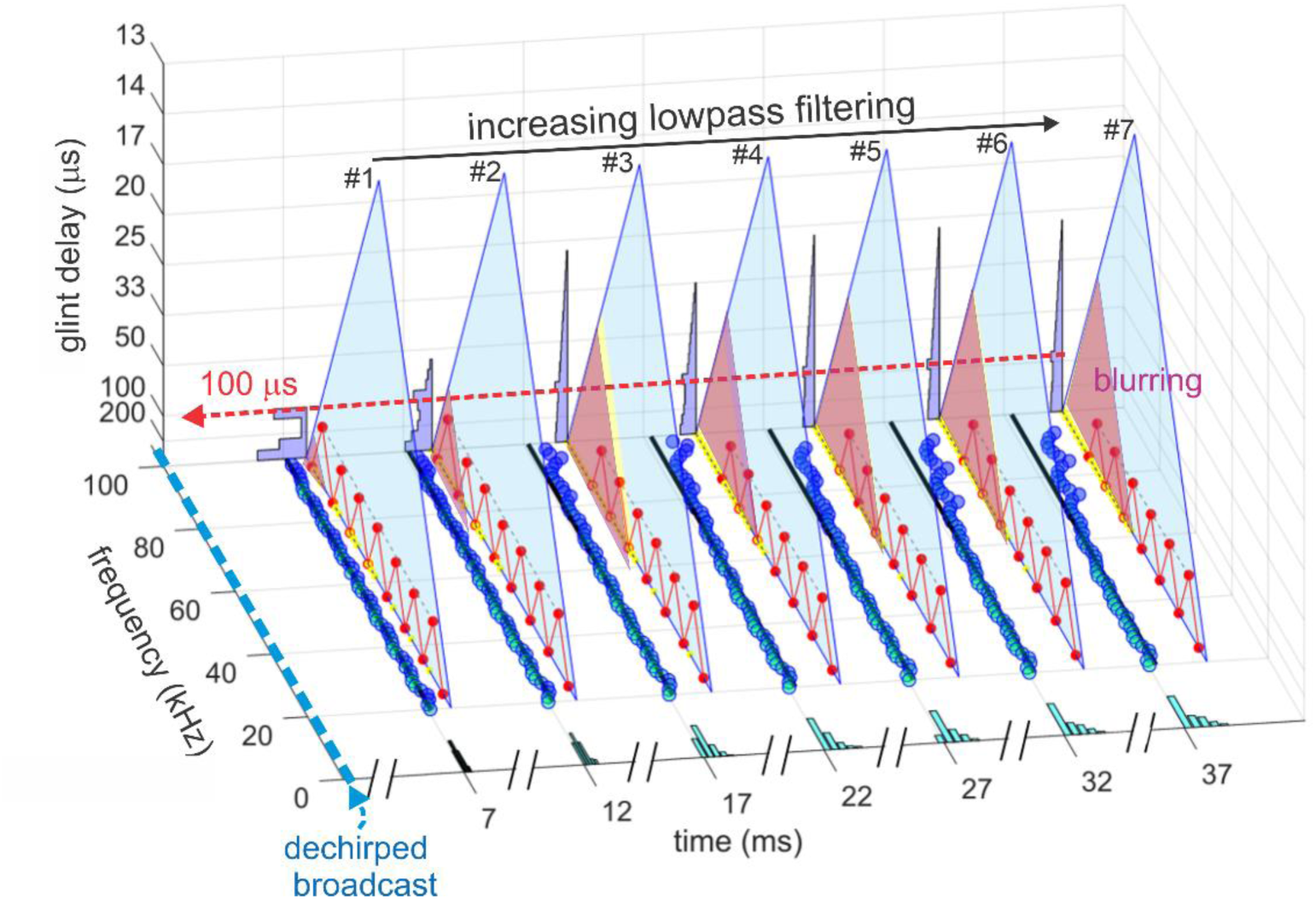
3D SCAT display for lowpass FM echoes. 3D SCAT display of range-delay and glint-delay estimates for the seven numbered 100-μs lowpass FM echoes in Fig 16. Range-delay histograms along the horizontal X axis here are also shown in Fig 16B. The glint-delay estimates derived from the triangular network are projected onto the X-Z backplane. For the echo with minimal lowpass filtering (echo #1; see Fig 16B), the 10-kHz null spacing is easily seen as the corresponding 100-μs glint delay (red arrow), although even the slight amount of lowpass filtering in echo #1 is enough to start producing a few additional nulls at glint-delays around 250-300 μs (very small shaded purple triangle in corner of large triangular network), making a second peak in the glint-delay histogram. As lowpass filtering becomes stronger (echoes #2-#3), the 100-μs glint in the glint-delay histogram becomes swamped by the spread of the additional nulls that fill up the lowpass region (increasing size of shaded purple triangle in corner of large triangular network). These spurious nulls (and the corresponding shaded triangles) stop spreading across more frequencies for echoes #4-#7. The 100-μs glint is barely discernible in the glint-delay histograms projected onto the X-Z plane. Progressively more lowpass filtering expands the single broad null caused by amplitude-latency trading, which grows and fills more of the large triangular network, but it does not spread below 50-60 kHz, so the addition of further spurious nulls (and the increasing size of the shaded triangle) thereafter stops.

#### Suppression of clutter by normalizing to image “energy.”

Fig 18A provides a closer view of the glint-delay histograms from the X-Z plane of Fig 17, replotted with glint delay on the horizontal axis (as in Fig 15A for dolphin click lowpass echoes). The series of lowpass FM echoes from Fig 17 are represented by the 100-μs glint spacing for echo #1, with increasing lowpass filtering from echoes #2-#6. As more of the fill-in nulls come to dominate the glint-delay mechanism, the 100-μs glint rides up to be on top of the wider histogram representing all the fill-in nulls. This contrast enhancement gets larger until the presence of the actual 100 μs glint becomes obscured by the effects of lowpass filtering. This progression illustrates the difficulty of representing echoes arriving from objects located off the broadcast beam’s axis as clutter. Note that even a slight degree of lowpass filtering is sufficient to create the fill-in nulls, which blur the glint-delay image by spreading its added glint estimates from about 50 to 400 μs to surround the actual second glint at 100 μs.

**Fig 18.**
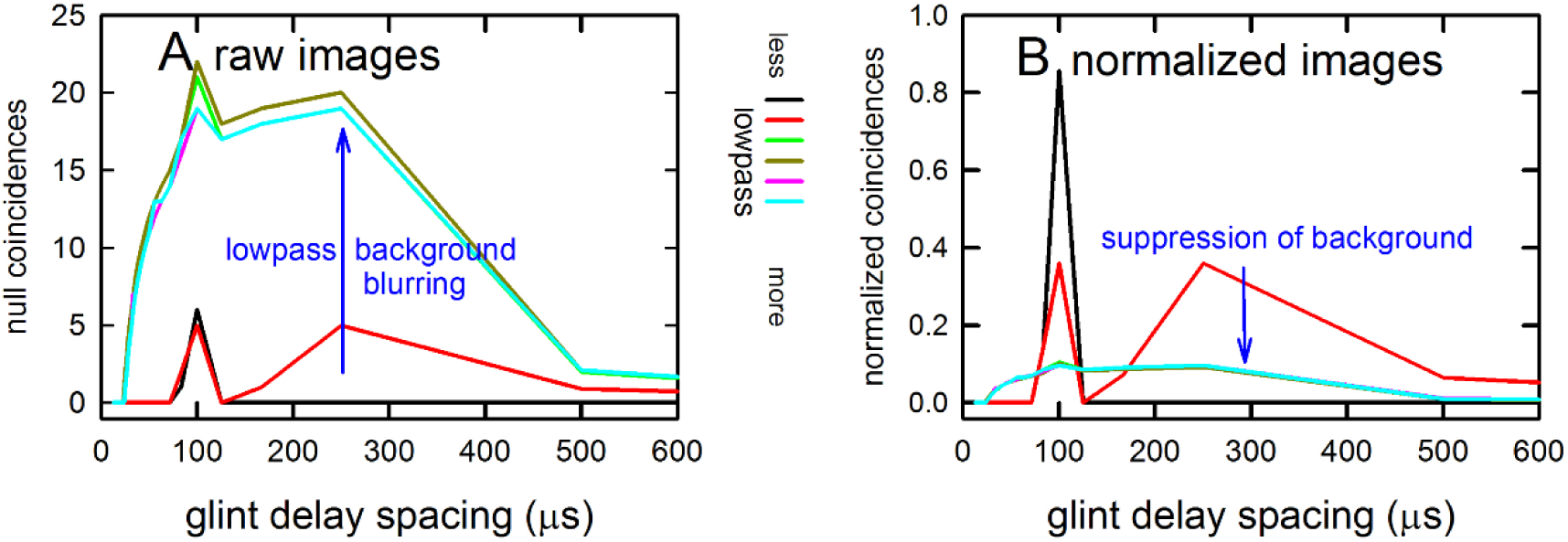
Glint-delay display for FM echoes. (**A**) Graph showing the glint-delay distributions displayed in Fig 17 on the X-Z plane and derived by the triangular network for the 100-μs two-glint FM echo (black) and the series of lowpass 100-μs two-glint FM echoes (red to green). The 100-μs peak is the salient feature of the two-glint echo from its 10 kHz null spacing, while the series of lowpass echoes show the 100 μs peak increasingly swamped by the broad set of nulls from the lowpass region of the triangular network in Fig 17. The lowpass echoes represent increasingly large angles of off-axis clutter, and the blurring effect of the many extraneous nulls detected across the lowpass region illustrates the masking effect of clutter interference with perception of the second glint. (**B**) The progressively stronger blurring effect that obscures the 100 μs glint in A is mitigated by using the total image “energy,” or the area under each curve, to normalize the display of glint delay, which becomes unmasked by the spread of energy across the glint-delay axis.

Blurring of the echo-delay image has been observed experimentally in big brown bats [33, 66]. The emergence of the blurring effect towards the long end of the glint-delay scale in Fig 18 coincides with the build-up of the perceived location of the artificially generated glint delays produced when the multiple, spurious nulls appear to fill in the wide lowpass null. The blurring mitigates clutter interference. Nevertheless, big brown bats successfully fly between rows of obstacles [67] and even follow along a row of obstacles forming a screen [122], explicitly using the locations of offside elements of the scene for steering flight. So, notwithstanding the blurring, the clutter is in fact perceived by the bat insofar as it can at least participate in sensorimotor control. In flight tests with big brown bats flying between two rows of obstacles, the bat can detect and turn away from a weakly-reflective net surrounded by strongly-reflective clutter [123]. The approximate signal-to-clutter ratio for detecting the net is −20 dB. How does the bat overcome the seeming disadvantage of such strong blurring of clutter images to reduce the interfering effects of the clutter? We intend to use the SCAT model in a binaural setting to examine how this might be done.

### SCAT model for naturalistic target echoes

Besides producing images for simulated two glint echoes, the SCAT model can also process echoes from naturalistic targets which contain multiple glints. We analyzed a series of previously-described echoes from a fluttering army worm moth (*Pseudaletia unipuncta*, body length 2 cm, wingspan 3.5 cm) [53]. The moth was attached to fine threads and placed in the center of a hoop with 1 m diameter, with its left body side facing the impinging sounds. The speaker and microphone were placed 34 cm from the insect, resulting a 2 ms delay between the pulse and received echo. The insect was ensonified with ultrasonic pulses that mimic the echolocation calls of big brown bats in approach phase of insect pursuit. The pulse is 1.25 ms long and has two harmonics, with the first from 25-50 kHz and the second 50-100 kHz. The ensonification rate is 33 Hz (*i.e.*, the pulse was played every 30 ms), which fortuitously matches the actual wingbeat cycle of the moth in this experiment. Echoes from the moth are cut out of the continuous recording and strung together to compress them into a 25 ms sequence shown in Fig 19A. Both the time waveforms and the spectrograms of the echoes are illustrated. The overall amplitude of the echoes is modulated by the moth’s wingbeats. In the first half of the sequence, echoes become stronger and then weaker with no distinct spectral notches, indicating the reflections are mainly from instants when the nearest (left) moth wing is oriented perpendicular to the incident sounds, before it flaps back toward the body and away from being acoustically perpendicular. The second half of the sequence consists of weaker echoes with prominent spectral notches (red arrows) emerging in the spectrograms. When the left wing has flapped into a position where both wings and the moth’s body received similar amount of sound and return glint reflections of similar amplitudes (shown in Fig 19A), the reflections interfere more equally with each other, producing those spectral notches. The SCAT display of echoes #14 – 18 from Fig 19A is shown in Fig 19B, with three of the ten threshold crossings (thresholds #2 - #4) plotted on X-Y plane, and the glint delay estimates on the X-Z plane. The current SCAT model provides a crude picture of the moth’s geometry, primarily under the assumption that there are just two prominent glints. The configuration of echoes overall is consistent with a small number of glints that characterize the insect [53].

**Fig 19.**
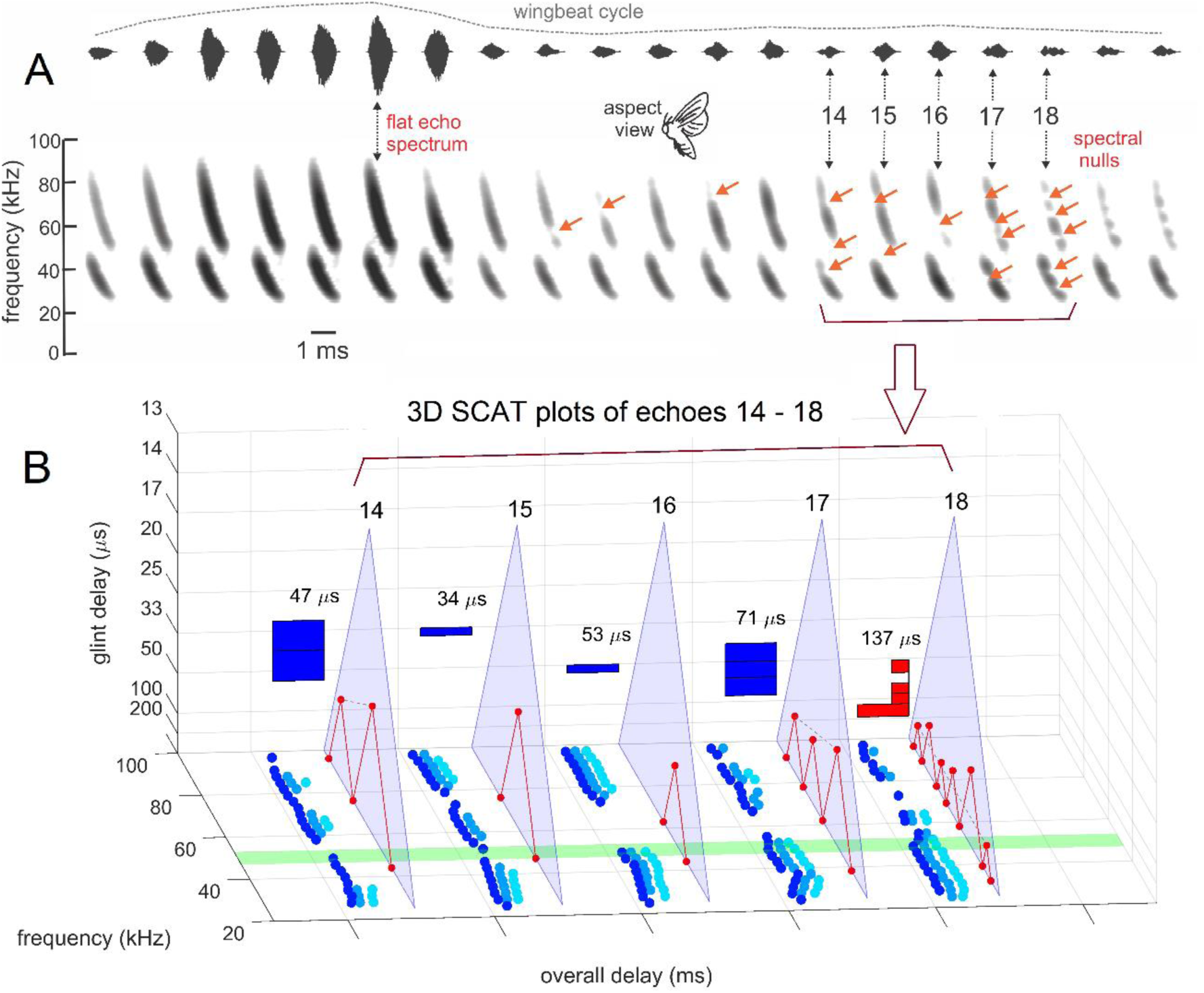
SCAT displays for fluttering moth echoes. (**A**) Horizontal stream of echoes (top) and spectrograms (bottom) recorded during a wingbeat cycle from a fluttering army worm moth ensonified from the side (aspect view insert) [53]. The 1.25 ms echoes from successive ensonifications were extracted and strung together to form this sequence. Echo amplitude is modulated by wingbeats, largely due to a wing moving into a posture perpendicular to the incident sound and creating a specular reflection with a flat spectrum. At other orientations, the weaker wing reflections bring up the glint reflections from other moth body parts so that interference becomes stronger and spectral nulls (red arrows) are more obvious (echoes 15-18). (**B**) 3D SCAT plots for echoes 14 – 18 from A. Three of the ten threshold crossings (#2 to #4) are displayed on X-Y plane, marked in dark blue (left) to light blue (right), respectively. For echoes that contain prominent notches, the triangular network finds and displays glint delays that change according to the wingbeats to portray the acoustic version of target shape.

## Discussion

### The SCAT model of biosonar perception

The SCAT model acquires broadband biosonar broadcasts and echoes, represents them as time-frequency spectrograms using parallel bandpass filters, translates the filtered signals into ten parallel amplitude threshold levels, and then operates on the resulting time-of-occurrence values at each frequency to estimate overall echo range delay (t) and the structure of the echo spectrum by depicting it as a series of local frequency nulls arranged regularly along the frequency axis of the spectrograms after dechirping them relative to the broadcast (Figs 1,2). The regular frequency spacing of the nulls (Δf) is explicitly transposed into its inverse, the time-delay separation (Δt) of two closely-spaced reflections from a target that contains these glints. Both range delay and glint delay, the principal outputs of the model, are known from experiments with big brown bats to be perceived as numerical values of time equated with distance from the bat, even though the source of one part of the perception is direct estimation of echo delay and the other part is indirect estimation obtained by transforming the spectrum of echoes—the pattern of ripples—into the glint delay spacing inferred from the spectrum.

The SCAT model is described here in largely in terms of its ability to dissect two-glint echoes into their component delays by bridging the gap between direct estimation of delay from the time domain and transformation of delay from the frequency domain. This keeps the explanation of the algorithms and their associated displays easy to follow. The mechanism reaps an unexpected benefit for suppressing clutter interference in that it is driven to an extreme representation of spurious glints by the lowpass filtering that characterizes offside or distant clutter. We defer a systematic examination of real three- and four-glint echoes to future work because further progress depends on understanding how information about the phase of wideband echoes is acquired from the initial, rich bandpass representation (Fig 5) and then carried through subsequent stages of processing to appear in the SCAT displays [22, 101]. This question is related to binaural processing and reconstruction of target shape from multiple aspect views [97]. Experimental results show that both bats and dolphins perceive the phase periodicity of echoes [28–32,121], but it is unclear how this is possible from exiting knowledge of auditory responses [21, 124]. The SCAT model provides a tool for examining how the parameters of sound reception and early auditory representation allow the equivalent of phase information to pass through the processing stream and appear in the resulting images.

### The SCAT display and the bat’s perception

The outstanding theoretical question posed by the SCAT 3D displays (Figs 7, 8, 10, 11, 13, 14, 15, 17, 19) concerns their relation to what the bat itself actually perceives. Psychophysical experiments show that big brown bats perceive the distance to targets from the delay of its echoes, with longer delays directly perceived as longer ranges [84]. Extending neural responses latencies by amplitude-latency trading directly leads to perception of longer ranges, so the latencies of neural responses convey the arrival time of echoes relative to broadcasts upward through the auditory system to be manifested in perception. Big brown bats also perceive the time delay of the second glint in terms of the slightly longer range to the second glint (84). If the glint separation is shorter than the integration-time for echo reception, the spectral interference nulls created when the two glint reflections arrive to be transduced by the inner ear are the acoustic cue for the second glint, but that glint is perceived unambiguously *as its delay*, not just *as a spectral coloration akin to timbre*. This is the spectrogram transformation.

The displays derived from the SCAT model are *pictures* drawn on 3D plots having axes presented as visual images for human interpretation. The question is this: When the bat perceives an object by echolocation, does it form an “image” constructed on neural scales arranged as spatial neural maps with dimensions of distance and glint separation analogous to the dimensions of the pictures? Or, does it perceive echo delays on a scale from near to far without any corresponding near and far in topographic neural space? In part, this is a question of fundamental concern for perceptual psychology and cognitive science. Do sharply registered spatial percepts require correspondingly sharp spatial neural substrates, or instead can the pattern of timing in neural responses distributed across a population of neurons, with at most a poorly-defined topography, evoke the perception of a sharply-registered perception of distance? How does perceptual sharpness depend on the sharpness of neuronal registration? In conjunction with behavioral and neural experimental results, the SCAT model provides a tool for addressing this question. The core problem is that individual neurons in the big brown bat’s auditory system responding to FM chirps and echoes do so with on average only one neuronal action potential, or spike, for each sound [8]. The bat’s perceptions thus might arise from the greater precision implicit in the timing of spikes *distributed across groups of neurons, not in topographic place defined by the less precise tuning to delay or frequency*. This issue would not arise if it were not for the existence of psychophysical results that point to the reality of biosonar percepts of greatly higher acuity than can be explained from existing neurophysiological results [21]. It seems inescapable that a defining role for perception resides in the timing of neural responses—that is, *when* some neurons respond, not predominantly *which* neurons express these responses.

### Technological implications of the SCAT model

The chief technological question follows from the conceptual question described above. Manifestly it is desirable to have the precision, quality, and real-time capabilities so clearly demonstrated by echolocating bats and dolphins somehow incorporated into man-made sonar and radar systems. To reach this goal, how much of the internal perceptual machinery has to be recreated explicitly in hardware and software? Contemporary systems use digital signal processing, which is based on multiple-bit arithmetic and the computational bottlenecks created by recursive algorithms and the serial nature of computer programs more generally. In such systems, higher speeds are achieved by employing multiple digital processors in a parallel design. The lesson for technology offered by the auditory system in the context of biosonar is that virtually total parallel design is possible to achieve, with the accompanying advantages for real time performance. The types of neuronal circuits invoked for auditory modeling are coincidence comparisons—essentially one-bit AND operations that keep track not only of the occurrence of coincidences by the times at which they occur [35]. If in fact bats do perceive targets as spatially-dimensioned images without developing similarly sharp topographically-dimensioned neural displays, how do we proceed to develop bio-inspired sonar systems that at least try to adopt this principle? Does the future of biomimetic sonar have to wait on further progress in the area of machine hearing to shape the selectivity of neural responses to different features of pulses and echoes?

### Limitations and specific future work

The interference between the echoes of two glints disappear when the spacing between the two is more than 300 µs for short clicks (see Fig 10), and it is present for longer pulses like bat FM broadcasts, but at 500 µs, spectral nulls are so close together that the ridge patterns which SCAT model based on to find notches are lost. To find the glint spacing in this case, SCAT model adopted another approach to treat the two glints as separate targets, and then reduced the echo interval to find the crossings of both glint echoes with thresholds. In this case, the glint spacing was found not through the spectral notches but the delay estimate derived from dechirped representations of both glints. This results in a display of both target location and target shape. The bat seems able to fuse both temporally measured delays and spectrally estimated delay differences onto the same perceptual dimension. The SCAT model can be a tool for further modeling to determine how perceived dimensions of the scene are coupled to guidance for steering the head and sonar beam along with steering flight.

Accuracy and efficiency are both important in engineered sonar, but enhancement of one may compromise the other. For example, the accuracy of SCAT model may be improved by using smaller frequency subbands in the filterbank; however, because the program loops through each band to find threshold crossings, the processing time will increase accordingly. We propose a potential solution in engineering applications that using two rounds to pinpoint the glint spacing. Specifically, the first round will use 1 kHz increment and infer the geometry of the targets. For those with longer glint delays than 200 µs, the SCAT model can use smaller frequency increments (*e.g*., 0.5 kHz) to pinpoint more precisely the spectral notches and thus provide a more accurate estimate of glint spacing. This can not only avoid using one universal small subband that diminish the general efficiency of SCAT model, but also can give relatively accurate results based on the estimate of target geometry from the first round of processing. This change does not apply to range delay or distance estimation, since even with a 1 kHz increment in filterbank frequencies, there are enough channels to make a close prediction.

As a final point, the SCAT model described here is a one-eared, or monaural, model, whereas echolocating animals use both ears in a synchronized manner. It is essential to develop a binaural version of the SCAT model to locate multiple objects, at the very least in the range-crossrange plane. The reason goes beyond just localization, however, because the stereo auditory view not only of the plane but of each object’s glints offers a way to advance target-based synthetic-aperture imaging for reconstructing shape from multiple aspect views, using the leading glint’s range delay in each ear and disparities in the trailing glint’s glint delays at the two ears.

## Acknowledgments

This research was supported by Office of Naval Research Grant (N00014-14-1-05880) to JAS, and Office of Naval Research Multidisciplinary University Research Initiative (N00014-17-1-2736) to AMS and JAS.

## References

1. Griffin DR. Listening in the dark. New Haven: Yale University Press; 1958.

2. Neuweiler G. Biology of bats. Oxford: Oxford University Press; 2000.

3. Au WWL. The sonar of dolphins. New York: Springer; 1993.

4. Thomas JA, Moss CF, Vater M, editors. Echolocation in bats and dolphins. Chicago: Univ. of Chicago Press; 2004.

5. Surlykke A, Nachtigall PE, Fay RR, Popper AN, editors. Biosonar. New York: Springer; 2014.

6. Fenton MB, Grinnell AD, Popper AN, Fay RR, editors. Bat bioacoustics. Springer: New York; 2016.

7. Teeling EC, Jones G, Rossiter SJ. Phylogeny, genes, and hearing: Implications for the evolution of echolocation in bats. In: Fenton MB, Grinnell AD, Popper AN, Fay RR, editors. Bat bioacoustics. Springer: New York; 2016. p. 25–54.

8. Ballieri A, editor. Biologically inspired radar and sonar: Lessons from nature. London: IET Press; 2017.

9. Simmons JA. Theories about target ranging in bat sonar. Acoustics Today;13: 43–51; 2017.

10. Simmons JA, Gaudette JE. Biosonar signal processing by frequency-modulated bats. IET Radar Sonar Navig. 2012; 6: 556–565.

11. Simmons JA. Bats use a neuronally implemented computational acoustic model to form sonar images. Curr Opin Neurobiol. 2012; 22:311–319.

12. Simmons JA, Gaudette JA, Warnecke M. Biosonar inspired signal processing and acoustic imaging from echolocating bats. In: Balleri A, editor. Biologically inspired radar and sonar: lessons from nature. London: IET Press; 2017. p5.

13. Fenton MB, Jensen FH, Kalko EKV, Tyack PL. Sonar signals of bats and toothed whales. In: Surlykke A, Nachtigall PE, Fay RR, Popper AN, editors. Biosonar. New York: Springer; 2014. p. 11–59.

14. Madsen PT, Surlykke, A. Echolocation in air and water. In: Surlykke A, Nachtigall PE, Fay RR, Popper AN, editors. Biosonar. New York: Springer; 2014. p. 257–304.

15. Au WWL, Suthers RA. Production of biosonar signals: Structure and form. In: Surlykke A, Nachtigall PE, Fay RR, Popper AN, editors. Biosonar. New York: Springer; 2014. p. 61–105.

16. Metzner W, Müller, R. Ultrasound production, emission, and reception. In: Fenton MB, Grinnell AD, Popper AN, Fay RR, editors. Bat bioacoustics. Springer: New York; 2016.p. 55–91.

17. Henson OW Jr. The ear and audition. In: Wimsatt WA, editor. Biology of bats vol 2. Academic Press: New York; 1970. p. 181–264.

18. Jen PHS. Electrophysiological analysis of the echolocation system of bats. In: Contributions to sensory physiology, 6. New York: Academic Press; 1982; p. 115–168.

19. Pollak GD, Casseday P. The neural basis of echolocation in bats. New York: Springer; 1989.

20. Covey E, Casseday JH. Timing in the auditory system of the bat. Ann Rev Physiol. 1999; 61: 457–476.

21. Pollak G. Organizational and encoding features of single neurons in the inferior colliculus of bats. In: Busnel R-G, Fish JF, editors. Animal Sonar Systems. Plenum Press: New York; 1980. p. 549–587.

22. Saillant PA, Simmons JA, Dear SP, McMullen TA. A computational model of echo processing and acoustic imaging in frequency-modulated echolocating bats: The spectrogram correlation and transformation receiver. J Acoust Soc Am. 1993; 94: 2691–712.

23. Wiegrebe L. An autocorrelation model of bat sonar. Biol Cybern. 2008; 98: 587–595. doi: 10.1007/s00422-008-0216-2.

24. Sanderson MI, Neretti N, Intrator N, Simmons JA. Evaluation of an auditory model for echo delay accuracy in wideband biosonar. J Acoust Soc Am. 2003; 114: 1648–1659.

25. Simmons JA, Ferragamo MJ, Moss CF. Echo-delay resolution in sonar images of the big brown bat, *Eptesicus fuscus*. Proc Nat Acad Sci USA. 1998; 95: 12647–12652.

26. Neretti, N., Sanderson, N. I., Intrator, N., and Simmons, J. A. Time-frequency computational model for echo-delay resolution in sonar images of the big brown bat, *Eptesicus fuscus*. J Acoust Soc Am. 2003; 113: 2137–2145.

27. Au WWL, Simmons JA (2007) Echolocation in dolphins and bats. Physics Today. 2007; 60: 40–45. doi: 10.1063/1.2784683.

28. Simmons JA. Perception of echo phase information in bat sonar. Science. 1979; 204: 1336–1338.

29. Simmons JA, Ferragamo M, Moss CF, Stevenson SB, Altes RA. Discrimination of jittered sonar echoes by the echolocating bat, *Eptesicus fuscus*: the shape of target images in echolocation. J Comp Physiol A. 1990; 167: 589–616.

30. Simmons JA. Evidence for perception of fine echo delay and phase by the FM bat, *Eptesicus fuscus*. J Comp Physiol A. 1993; 172: 533–547.

31. Finneran JJ, Jones R, Mulsow J, Houser DS, Moore, PW. Jittered echo-delay resolution in bottlenose dolphins (*Tursiops truncatus*). J Comp Physiol A. 2019; 205: 125–137.

32. Finneran JJ, Jones R, Guazzo RA, Strahan MG, Mulsow J, Houser DS, Branstetter BK, Moore PW. Dolphin echo-delay resolution measured with a jittered-echo paradigm. J Acoust Soc Am. 2020; 148: 374–388.

33. Bates ME, Simmons JA, Zorikov TV. Bats use echo harmonic structure to distinguish targets from clutter and suppress interference. Science. 2011; 333: 627–630.

34. Cohen L. Time-Frequency Analysis. New York: Prentiss Hall; 1995.

35. Lyon RF. Human and machine hearing. Cambridge UK: Cambridge University Press; 2017.

36. Skolnik MI. Introduction to radar systems, 2nd ed. New York: McGraw-Hill, 1980.

37. Woodward PM. Probability and information theory with applications to radar. 2nd ed. New York: Pergamon Press; 1953.

38. Simmons JA. Noise interference with echo delay discrimination in bat sonar. J Acoust Soc Am. 2017; 142: 2942–2952. doi.org/10.1121/1.5010159.

39. Avissar M, Wittig JH Jr, Saunders JC, Parsons TD. Refractoriness enhances temporal coding by auditory nerve fibers. J Neurosci. 2013; 33: 7681–7690.

40. Heil P, Peterson AJ. Spike timing in auditory-nerve fibers during spontaneous activity and phase locking. Synapse. 2016; 71: 5–36.

41. Peterson AJ, Heil P. Phase locking of auditory-nerve fibers: the role of lowpass filtering by hair cells. J Neurosci. 2020; 10.1523/JNEUROSCI.2269-19.2020.

42. Joris PX, Smith PH. The volley theory and the spherical cell puzzle. Neurocience 2008; 154: 65–76.

43. Bates ME, Simmons JA. Effects of filtering of harmonics from biosonar echoes on delay acuity by big brown bats (*Eptesicus fuscus*). J Acoust Soc Am. 2010; 128: 936–946.

44. Simmons JA, Neretti N, Intrator N, Altes RA, Ferragamo MJ, Sanderson MI. Delay accuracy in bat sonar is related to the reciprocal of normalized echo bandwidth, or Q. Proc Natl Acad Sci. USA. 2004; 101: 3638–3643.

45. Stilz WP, Schnitzler HU. Estimation of the acoustic range of bat echolocation for extended targets. J Acoust Soc Am. 2012; 132: 1765–1775.

46. Simmons JA, Houser D, Kloepper L. Localization and classification of targets by echolocating bats and dolphins. In: Surlykke A, Nachtigall PE, Fay RR, Popper AN, editors. Biosonar. New York: Springer; 2014. p. 169–193.

47. Kick SA, Simmons JA. Automatic gain control in the bat’s sonar receiver and the neuroethology of echolocation. J Neurosci. 1984; 4: 2705–2737.

48. Wahlberg M, Surlykke A. Sound intensities of biosonar signals from bats and toothed whales. In: Surlykke A, Nachtigall PE, Fay RR, Popper AN, editors. Biosonar. New York: Springer; 2014. p. 107–141.

49. Nachtigall PE, Schuller G. Hearing during echolocation in whales and bats. In: Surlykke A, Nachtigall PE, Fay RR, Popper AN, editors. Biosonar. New York: Springer; 2014. p. 143–167.

50. Simmons JA, Chen L. Acoustic basis for target discrimination by FM echolocating bats. J Acoust Soc. Am. 1989; 86: 1333–1350.

51. Murchison AE. Detection range and range resolution of echolocating bottlenose porpoise (*Tursiops truncatus*). In: Busnel RG, Fish JF, editors. Animal Sonar Systems. New York: Plenum. 1980. p. 43–70.

52. Møhl B. Target detection by echolocating bats. In: Nachtigall PE, Moore PWB, editors. Animal Sonar: Processes and performance. New York: Plenum; 1988. p. 435–450.

53. Moss CF, Zagaeski M. Acoustic information available to bats using frequency modulated sonar sounds for the perception of insect prey. J Acoust Soc Am. 1994; 95: 2745–2756.

54. Houston RD, Boonman AM, Jones G. Do echolocation signal parameters restrict bats’ choice of prey? In: Thomas JA, Moss CF, Vater M, editors. Echolocation in bats and dolphins. Chicago: Univ. of Chicago Press; 2004. p.339–345.

55. Hartley DJ, Suthers RA. The sound emission pattern of the echolocating bat, *Eptesicus fuscus*. J Acoust Soc Am. 1989; 85:1348–1351.

56. Saillant PA, Simmons JA, Bouffard F, Lee DN, Dear SP. Biosonar signals impinging on the target during interception by big brown bats, *Eptesicus fuscus*. J Acoust Soc Am. 2007; 21: 3001–3010.

57. Simmons JA. Temporal binding of neural responses for focused attention in biosonar. J Exp Biol. 2014; 217: 2834–2843.

58. Masters WM, Moffat AJM, Simmons JA. Sonar tracking of horizontally moving targets by the big brown bat, *Eptesicus fuscus*. Science. 1985; 228: 1331–1333.

59. Ghose K, Moss CF. The sonar beam pattern of a flying bat as it tracks tethered insects. J Acoust Soc Am. 2003; 114: 1120–1131.

60. Ghose K, Moss CF. Steering by hearing: a bat’s acoustic gaze is linked to its flight motor output by a delayed, adaptive filter law. J Neurosci. 2006; 26: 1704–1710.

61. Falk B, Jakobsen L, Surlykke A, Moss CF. Bats coordinate sonar and flight behavior as they forage in open and cluttered environments. J Exp Biol. 2014; 217: 4356–4364.

62. Moss CF, Bohn K, Gilkenson H, Surlykke A. Active listening for spatial orientation in a complex auditory scene. PLoS Biol. 2006; 4: 615–626. doi:10.1371/journal.pbio.0040079.

63. Grinnell AD. Hearing in bats: an overview. In: Popper AN, Fay RR, editors. Hearing by bats. New York: Springer; 1995. p. 1–36.

64. Ming C, Bates ME, Simmons JA. How frequency hopping suppresses pulse-echo ambiguity in bat biosonar. Proc Natl Acad Sci USA. 2020; 117: 17288–17295. doi/10.1073/pnas.200110511.

65. Stamper SA, Bates ME, Benedicto D, Simmons JA. Role of broadcast harmonics in echo delay perception by big brown bats. J Comp Physiol A. 2009; 195: 79–89.

66. Warnecke M, Simmons JA. Target shape perception and clutter rejection use the same mechanism in bat sonar. J Comp Physiol A. 2016; 202, 371–379.

67. Wheeler AR, Fulton KA, Gaudette JE, Matsuo I, Simmons JA. Echolocating big brown bats, *Eptesicus fuscus*, modulate pulse intervals to overcome range ambiguity in cluttered surroundings. Front Behav Neurosci. 2016; 10:125. doi: 10.3389/fnbeh.2016.00125.

68. Simmons JA, Hiryu S, Shriram U. Biosonar interpulse intervals and pulse-echo ambiguity in four species of echolocating bats. J Exp Biol. 2019; 222. doi: 10.1242/jeb.195446.

69. Hiryu S, Bates ME, Simmons JA, Riquimaroux H. FM echolocating bats shift frequencies to avoid broadcast-echo ambiguity in clutter. Proc Natl Acad Sci USA. 2010; 107: 7048–7053.

70. Altes RA. Sonar for generalized target description and its similarity to animal sonar systems. J Acoust Soc Am. 1976; 59: 97–105.

71. Griffin DR. Discriminative echolocation by bats. In: Busnel R-G, editor. Animal sonar systems: biology and bionics. Jouy-en-Josas, France: Laboratoire de Physiologie Acoustique; 1967. p. 273–300.

72. Griffin DR, Friend JH, Webster FA. Target discrimination by the echolocation of bats. J Exp Zool. 1965; 158: 155–168.

73. Airapetyants E Sh, Konstantinov AI. Echolocation in nature. Jerusalem: Israel Program for Scientific Translation; 1973.

74. Simmons JA, Lavender WA, Lavender BA, Doroshow CA, Kiefer SW, Livingston R, Scallet AC, Crowley DE. Target structure and echo spectral discrimination by echolocating bats. Science 1974; 186(4189): 1130–1132.

75. Habersetzer J, Vogler B. Discrimination of surface-structured targets by the echolocating bat *Myotis myotis* during flight. J Comp Physiol. 1983; 152: 275–282.

76. Schmidt S. Evidence for a spectral basis of texture discrimination in bat sonar. Nature 1988; 331: 617–619.

77. Simmons JA, Moss CF, Ferragamo M. Convergence of temporal and spectral information into acoustic images of complex sonar targets perceived by the echolocating bat, *Eptesicus fuscus*. J Comp Physiol A. 1990; 166: 449–470.

78. Mogdans J, Schnitzler H-U. Range resolution and the possible use of spectral information in the echolocating bat, *Eptesicus fuscus*. J Acoust Soc Am. 1990; 88: 754–757.

79. Mogdans J, Schnitzler H-U, Ostwald J. Discrimination of 2-wavefront echoes by the big brown bat, *Eptesicus fuscus*: Behavioral experiments and receiver simulations. J Comp Physiol A. 1993; 172: 309–323.

80. Schmidt S. Perception of structured phantom targets in the echolocating bat, *Megaderma lyra*. J Acoust Soc Am. 1992; 91: 2203–2223.

81. Grunwald, J-E, Schornich S, Wiegrebe L. Classification of natural textures in echolocation. Proc Nat Acad Sci USA. 2004; 101: 5670–5674.

82. Fitzlaff J, Schuchmann M, Grunwald JE, Schuller G, Wiegrebe L. Object-oriented echo perception and cortical representation in echolocating bats. PLoS Biol. 2007; 5: e100. doi:10.1371/journal.pbiol.0050100.

83. Yovel Y, Matthias OF, Stilz P, Schnitzler U-U. Complex echo classification by echolocating bats: a review. J Comp Physiol A. 2011; 197: 475–490. DOI 10.1007/s00359-010-0584-7

84. Simmons JA, Ferragamo MJ, Saillant PA, Haresign T, Wotton JM, Dear SP, Lee DN. Auditory dimensions of acoustic images in echolocation. In: Popper AN, Fay RR, editors. Editors. Hearing by bats. New York: Springer; 1995. p. 146–190.

85. Shriram U, Simmons JA. Echolocating bats perceive natural-size targets as a unitary class using micro-spectral ripples in echoes. Behav Neurosci. 2019; 133: 297–304. doi.org/10.1037/bne0000315.

86. Matsuo I, Imaizumi T, Akamatsu T, Furusawa M, Nishimori Y. Analysis of the temporal structure of fish echoes using the dolphin broadband sonar signal. J Acoust Soc Am. 2009; 126: 444. doi: 10.1121/1.3147505.

87. Au WWL, Branstetter BK, Benoit-Bird KJ, Kastelein RA. Acoustic basis for fish prey discrimination by echolocating dolphins and porpoises. J Acoust Soc Am. 2009; 126: 460–467.

88. Masters WM, Jacobs SC, Simmons JA. The structure of echolocation sounds used by the big brown bat, *Eptesicus fuscus*: Some consequences for echo signal processing. J Acoust Soc Am. 1991; 89: 1402–1413.

89. Au WWL, Benoit-Bird KJ. Automatic gain control in the echolocation system of dolphins. Nature. 2003; 423: 861–863.

90. Surlykke A, Moss CF. Echolocation behavior of big brown bats, *Eptesicus fuscus*, in the field and the laboratory. J Acoust Soc Am. 2000; 108: 2419–2429.

91. O’Neill WE. The bat auditory cortex. In: Hearing by bats, Popper AN, Fay RR, editors. New York: Springer; 1995. p. 416–480.

92. Suga N, Olsen JF, Butman JA Specialized subsystems for processing biologically important complex sounds: cross-correlation analysis for ranging in the bat’s brain. Cold Spring Harbor SympQuant Biol. 1990: 55: 585–597.

93. Simmons JA, Saillant PA, Ferragamo MJ, Haresign T, Dear SP, Fritz JB, McMullen TA. Auditory computations for acoustic imaging in bat sonar. In: Hawkins HL, McMullen TA, Popper AN, Fay RR, editors. Auditory Computation. New York: Springer; 1996. p. 401–468.

94. Zorikov TV, Dubrovsky NA. Echo processing procedure in bottlenose dolphins. Oceans 2003 Proceedings (IEEE Cat. No. 03CH37492), 2003: 7 pp. doi: 10.1109/Oceans.2003.178577.

95. Branstetter, BK, Van Alstyne KR, Strahan MG, Tormey MN, Wu T, Breitenstein RA, Houser DS, Finneran JJ, Xitco MJ Jr. Spectral cues and temporal integration during cylinder echo discrimination by bottlenose dolphins (*Tursiops truncatus*). J Acoust Soc Am. 2020; 148: 614–626.

96. Accomando AW, Mulsow J, Houser DS, Finneran JJ. Classification of biosonar target echoes on coarse and fine spectral features in the bottlenose dolphin (*Tursiops truncatus*). J Acoust Soc Am. 2020; 148: 1642–1646.

97. DeLong CM, Bragg R, Simmons JA. Evidence for spatial representation of object shape in echolocating bats (*Eptesicus fuscus*). J Acoust Soc Am. 2008; 123: 4582–4598.

98. Mertens A, Mertens A. (1999) Signal Analysis: wavelets, filter banks, time-frequency transforms and applications. Wiley: New York.

99. Boashash B. Time frequency signal analysis and processing. 2^nd^ ed. New York: Elsevier; 2016. pp.577–625 doi.org/10.1016/B978-0-12-398499-9.00014-5.

100. Gaunaurd GC, Strifors HC. Signal analysis by means of time-frequency (Wigner-type) distributions-applications to sonar and radar echoes. Proc IEEE—Special issue on: Time-frequency analysis. 1996:84: 1231–1248.

101. Peremans H, Hallam J. The spectrogram correlation and transformation receiver, revisited. J Acoust Soc Am. 1998; 104: 1101–1110.

102. Matsuo I, Kunugiyama K, Yano M. An echolocation model for range discrimination of multiple closely spaced objects: Transformation of spectrogram into the reflected intensity distribution. J Acoust Soc Am. 2004; 115: 920–928.

103. Park M, Allen R. Pattern-matching analysis of fine echo delays by the spectrogram correlation and transformation receiver. J Acoust Soc Am. 2010; 128: 490–1500.

104. Georgiev K, Balleri A, Stove A, Holderied MW. Bio-inspired processing of radar target echoes. Radar Sonar & Navigation IET. 2018; 12: 1402–1409.

105. Altes RA. Texture analysis with spectrograms. IEEE Transactions Sonics and Ultrasonics. 1984; 31: 407–417. doi: 10.1109/T-SU.1984.31521.

106. Sanderson MI, Simmons JA. Neural responses to overlapping FM sounds in the inferior colliculus of echolocating bats. J Neurophysiol. 2000; 83: 1840–1855.

107. Sanderson MI, Simmons JA. Selectivity for echo spectral shape and delay in the auditory cortex of the big brown bat, *Eptesicus fuscus*. J Neurophysiol. 2003; 87: 2823–2834.

108. Macías S, Bakshi K, Garcia-Rosales F, Hechavarria JC, Smotherman, M. Temporal coding of echo spectral shape in the bat auditory cortex. PLoS Biol. 2020; 18(11):e3000831. doi.org/10.1371/journal.pbio.3000831.

109. Bodenhamer R, Pollak GD. Time and frequency domain processing in the inferior colliculus of echolocating bats. Hear Res. 1981; 5: 317–335.

110. Pollak GD, Marsh DS, Bodenhamer R, Souther A. Characteristics of phasic-on neurons in the inferior colliculus of anaesthetized bats with observations related to mechanisms for echoranging. J Neurophysiol. 1977; 40, 926–941.

111. Jen PHS, Schlegel, PA. Auditory physiological properties of neurons in the inferior colliculus of the big brown bat, *Eptesicus fuscus*. J Comp Physiol. 1982; 147: 351–363.

112. Haplea S, Covey E, Casseday JH. Frequency tuning and response latencies at three levels in the brainstem of the echolocating bat, *Eptesicus fuscus*. J Comp Physiol A. 1994; 174: 671–683.

113. Ferragamo MJ, Haresign T, Simmons JA. Frequency tuning, latencies, and responses to FM sweeps in the inferior colliculus of the echolocating bat, *Eptesicus fuscus*. J Comp Physiol A. 1998; 182: 65–79.

114. Ashmore J, Avan P, Brownell WE, Dallos P, Dierkes K, Fettiplace R, Grosh K, Hackney CM, Hudspeth AJ, Jülicher F, Lindner B, Martin P, Meaud J, Petit C, Santos Sachi JR, Canlon B. The remarkable cochlear amplifier. Hear Res. 2010; 266: 1–17.

115. Heil P, Peterson AJ. Basic response properties of auditory nerve fibers: a review. Cell Tissue Res. 2015; 361:129–158.

116. Klug A, Khan A, Burger RM, Bauer EE, Hurley LM, Yang L, Grothe B, Halvorsen MB, Park TJ. Latency as a function of intensity in auditory neurons: influences of central processing. Hear res. 2000:148: 107–123.

117. Burkard R, Moss CF. The brain-stem auditory-evoked response in the big brown bat (*Eptesicus fuscus*) to clicks and frequency-modulated sweeps. J Acoust Soc Am. 1994; 96: 801–810.

118. Ma X, Suga N. Corticofugal modulation of the paradoxical latency shifts of inferior collicular neurons. J Neurophysiol. 2008; 100: 1127–1134.

119. Luo J, Simmons AM, Beck QM, Macías S, Moss CF, Simmons JA. Frequency-modulated up-chirps produce larger evoked responses than down-chirps in the big brown bat auditory brainstem. J Acoust Soc Am. 2019; 146(3): 1672–1684. doi: 10.1121/1.5126022.120.

120. Simmons JA, Yashiro H, Kohler AL, Riquimaroux H, Funabiki K, Simmons AM. Long-latency optical responses from the dorsal inferior colliculus of Seba’s fruit bat. J Comp Physiol A. 2020; 206(6): 831–844. doi: 10.1007/s00359-020-01441-7.

121. Moss CF, Schnitzler H.-U. Accuracy of target ranging in echolocating bats: acoustic information processing. J Comp Physiol A. 1989; 165: 383–393.

122. Warnecke M, Lee W-J, Krishnan A, Moss CF. Dynamic echo information guides flight in the big brown bat. Front Behav Neurosci. 2016; 10: 1–11. https://doi.org/10.3389/fnbeh.2016.00081

123. Knowles JM, Barchi JR, Gaudette JE, Simmons JA. Effective biosonar echo-to-clutter rejection ratio in a complex dynamic scene. J Acoust Soc Am. 2015; 138: 1090–1101

124. Schörnich S, Wiegrebe L. Phase sensitivity in bat sonar revisited. J Comp Physiol A. 2008; 194: 61–67.

